# Age and infectious dose significantly affect disease progression after RHDV2 infection in naïve domestic rabbits

**DOI:** 10.1101/2021.05.26.445897

**Authors:** Robyn N. Hall, Tegan King, Tiffany O’Connor, Andrew J. Read, Jane Arrow, Katherine Trought, Janine Duckworth, Melissa Piper, Tanja Strive

## Abstract

Rabbit haemorrhagic disease virus 2 (RHDV2 or GI.2, referring to any virus with lagovirus GI.2 structural genes) is a recently emerged calicivirus that causes generalised hepatic necrosis and disseminated intravascular coagulation leading to death in susceptible lagomorphs (rabbits and hares). Previous studies investigating the virulence of RHDV2 have reported conflicting results, with case fatality rates ranging from 0% to 100% even within a single study. Lagoviruses are of particular importance in Australia and New Zealand where they are used as biocontrol agents to manage wild rabbit populations; wild rabbits threaten over 300 native species and result in economic impacts in excess of $200 million AUD to Australian agricultural industries. It is critically important that any pest control method is both highly effective (i.e., virulent, in the context of viral biocontrols) and has minimal animal welfare impacts. To determine whether RHDV2 might be a suitable candidate biocontrol agent, we investigated the virulence and disease progression of a naturally occurring Australian recombinant RHDV2 in both 5-week-old and 11-week-old New Zealand White laboratory rabbits after either high or low dose oral infection. Objective measures of disease progression were recorded through continuous body temperature monitoring collars, continuous activity monitors, and twice daily observations. We observed a 100% case fatality rate in both infected kittens and adult rabbits after either high dose or low dose infection. Clinical signs of disease, such as pyrexia, weight loss, and reduced activity, were evident in the late stages of infection. Clinical disease, i.e., welfare impacts, were limited to the period after the onset of pyrexia, lasting on average 12 hours and increasing in severity as disease progressed. These findings confirm the high virulence of this RHDV2 variant in naïve rabbits. While age and infectious dose significantly affected disease progression, the case fatality rate was consistently 100% under all conditions tested.

## Introduction

Rabbit haemorrhagic disease is a peracute, typically fatal, hepatitis of European rabbits (*Oryctolagus cuniculus*) culminating in generalised hepatic necrosis and disseminated intravascular coagulation in susceptible individuals [1]. It is caused by two lagoviruses in the family *Caliciviridae*. RHDV (or GI.1) was first reported as a cause of mass mortalities in farmed rabbits in China in 1984 [2] and subsequently became endemic in most of Europe, parts of Asia and Africa, Cuba, Australia, and New Zealand [3]. In Australia and New Zealand, it was deliberately released as a viral biocontrol agent to manage wild rabbit populations. It arrived in other regions through natural and/or anthropogenic transmission routes, and also caused sporadic disease in Mexico, USA, Uruguay, Canada, and elsewhere [1].

In 2010, the novel lagovirus RHDV2 (RHDVb or GI.2), which is genetically and antigenically distinct from RHDV, was reported from wild and domestic rabbits in France and Spain [4, 5]. RHDV2 can affect rabbits as young as 11 days old. In contrast, rabbits younger than approximately 8 weeks of age show an innate age-dependent resistance to disease caused by the closely related RHDV (or GI.1), although they are still permissive to infection [4]. Additionally, RHDV2 causes disease in several hare (*Lepus*) species and can overcome RHDV vaccinal protection [6-9]. After its emergence, RHDV2 rapidly spread via natural transmission pathways throughout Europe, parts of Africa and the Middle East, Australia, and Canada, and has recently been reported in wild *Lepus* and *Sylvilagus* species in the USA [10, 11]. It is now the dominant pathogenic lagovirus in these geographic regions.

Within the RHDV2 designation, there are several variants, which comprise non-structural (NS) coding sequences from other lagoviruses along with the GI.2 structural protein [12]. These variants arise through recombination at the junction between the viral polymerase and capsid protein (VP60) via a template switching mechanism shared by other caliciviruses [13]. While all RHDV2 variants reported appear to be pathogenic and have similar antigenic properties [12, 14-19], genetic diversity in the non-structural proteins between different variants may explain the widely discrepant results reported in previous RHDV2 pathogenesis studies. For example, mortality rates in field outbreaks reportedly ranged from 20% to 90% [4, 5, 20-27]. Additionally, RHDV2 mortalities after experimental infection occurred later and over a longer period than RHDV1 in some studies but not in others, and case fatality rates (CFR) were highly variable (0% to 100%), even between experiments within a single study [9, 14, 28-33]. There have also been conflicting reports of age-dependent mortality after RHDV2 challenge, suggesting in some cases that young rabbits may in fact be more susceptible than adults [4, 24, 33]. This information has been summarised in Table 1.

**Table 1.**
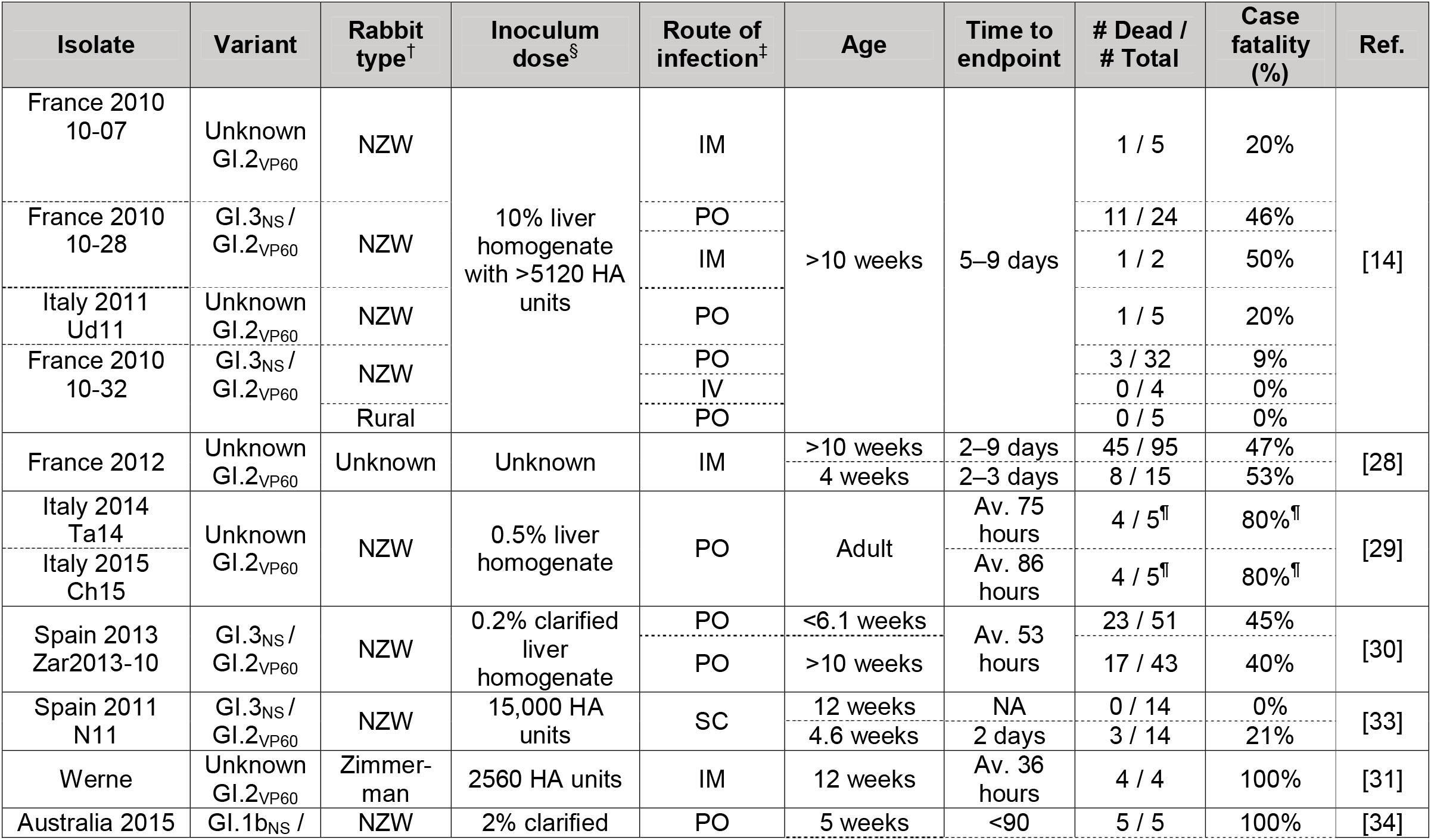
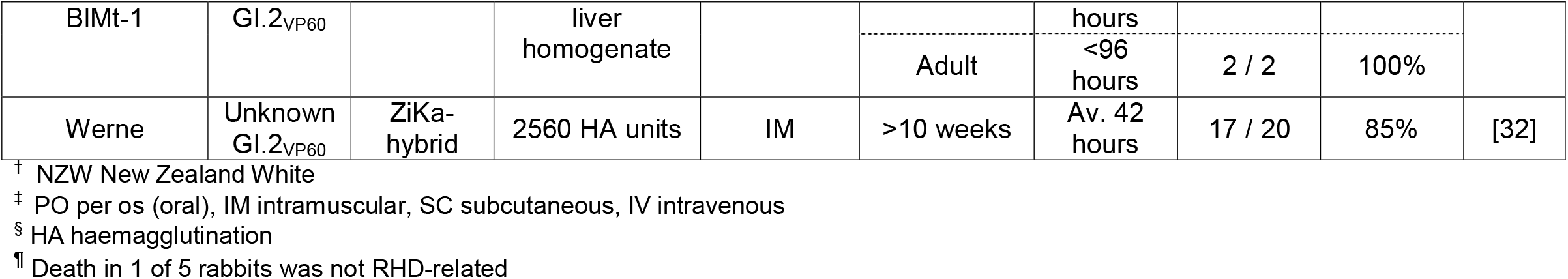
Summary of published RHDV2 experimental infection studies.

While some authors have suggested that RHDV2 has increased in virulence since its emergence [29], this doesn’t explain the variation observed within single studies. For many published studies, the full genome sequence of the isolate used for infection was not reported, so the specific RHDV2 variant used is unknown (Table 1). Additionally, the infectious dose is either not given or is difficult to compare between studies. Importantly, determination of infectious dose for lagoviruses must be measured by titration in rabbits since these viruses cannot be cultivated *ex vivo*. Taken together, the discordant findings reported in the literature suggest that unexplained factors may affect disease progression after RHDV2 infection, for example, virus variant, age at infection, infectious dose, and perhaps host genetics and route of infection.

This study aimed to thoroughly assess the virulence of an RHDV2 variant, a recombinant GI.1bP-GI.2 that is widely circulating in Australia, the Iberian peninsula and the Azores, Morocco, France, and Poland [12, 15, 16, 18, 19, 35-38]. We investigated the influence of age and infectious dose on disease progression in naïve laboratory rabbits using the natural oral route of infection. Although RHDV2 is already actively circulating in Australia, the ability to control the timing of releases through registration of this virus as an additional rabbit biocontrol agent would improve the effectiveness of biocontrol programs by allowing landholders to exploit synergistic interactions between different lagovirus variants.

## Materials and methods

### Animal trials

Five-week-old kitten (range 33–35 days at infection, weight 519–985 g [x□= 701 g], 18 male, 10 female) and 11-week-old adult (range 75–77 days at infection, 1909–2636 g [x□= 2177 g], 17 male, 11 female) New Zealand White rabbits were obtained from a laboratory rabbit breeding colony that was known to be free of benign and pathogenic lagoviruses. All rabbits were confirmed to be seronegative to the benign lagovirus RCV-A1 [39], as well as to RHDV1 [40] and RHDV2 [41], prior to infection. All work was conducted in accordance with the ‘Australian code for the care and use of animals for scientific purposes’ and was approved by the ‘CSIRO Wildlife, Livestock and Laboratory Animal’ animal ethics committee (permit #2018-06). Rabbits were housed individually in laboratory rabbit housing (Tecniplast, Italy) within a climate-controlled, insect-proof room. Independent trials were conducted using entire litter groups until the required sample size was met (Table 2).

**Table 2.** Experimental design for the assessment of RHDV2 virulence.

| Trial | Number of animals<sup>†</sup> | Age (weeks) | Sex (F:M) | Target inoculum dose (RID<sub>50</sub>)<sup>‡</sup> | Actual inoculum dose (capsid gene copies) |
| --- | --- | --- | --- | --- | --- |
| 1 | 4 + 1 control | 11 | 2:3 | 1000 | 1×10<sup>7</sup> |
| 2 | 6 + 1 control | 11 | 2:5 | 1000 | 7×10<sup>7</sup> |
| 3 | 2 + 1 control | 11 | 1:2 | 1000 | 6×10<sup>6</sup> |
|  | 4 |  | 2:2 | 50 | 3×10<sup>5</sup> |
| 4 | 8 + 1 control | 11 | 4:5 | 50 | 4×10<sup>5</sup> |
| 5 | 6 + 1 control | 5 | 1:6 | 1000 | 1×10<sup>7</sup> |
| 6 | 6 + 1 control | 5 | 4:3 | 1000 | 1×10<sup>7</sup> |
|  | 1 |  | 0:1 | 50 | 7×10<sup>5</sup> |
| 7 | 6 + 1 control | 5 | 4:4 | 50 | 3×10<sup>5</sup> |
| 8 | 5 + 1 control | 5 | 1:5 | 50 | 7×10<sup>5</sup> |

<sup>†</sup> Each trial included one uninfected control animal from the respective litter group.
<sup>‡</sup> RID<sub>50</sub> 50% rabbit infectious dose

Sample sizes were estimated based on demonstrating lethality in both 5-week-old and 11-week-old animals at two infectious doses. Calculations were performed using the following parameters: two independent study groups, dichotomous endpoint, anticipated incidence of 70% in infected animals (based on previously published studies) and 5% in uninfected animals (to account for potential adverse events [29]), alpha of 0.05, 80% power, and an enrolment ratio of 3:1 (to minimise the number of control animals required). This resulted in infection groups of 12 animals per age and infectious dose (four treatments: adult/high dose, adult/low dose, kitten/high dose, kitten/low dose) and uninfected control groups of 4 animals per age.

### Virus inoculum

Rabbits were infected orally (to mimic a natural route of infection) with a GI.1bP-GI.2 virus, MEN-1. This virus was first isolated from a dead wild rabbit found in Menangle, New South Wales (NSW), in April 2016. Virus stocks were prepared by the Elizabeth Macarthur Agricultural Institute (Menangle, NSW); briefly, virus was amplified in New Zealand White rabbits, liver homogenates were semi-purified, and freeze-dried virus stocks were prepared and titrated to derive the 50% rabbit infectious dose (RID_50_) of the virus stock. The genome sequence of this isolate is available under Genbank accession MW467791. For titration, groups of six rabbits were inoculated intramuscularly with 10-fold serial dilutions of the concentrated virus stock and the Reed-Muench method was used to determine the 50% endpoint [42]. Individual vials of freeze-dried virus stock (Elizabeth Macarthur Agricultural Institute, NSW, Australia) were reconstituted in sterile water immediately prior to each trial and diluted to the required dose. Rabbits were infected orally with either a high dose (1000 RID_50_) or low dose (50 RID_50_) of virus in a final volume of 0.7 ml, where the low dose was estimated as the amount of virus that a rabbit may reasonably consume on carrot bait during a biocontrol program. The high dose was selected to be 20 times the low dose. Each inoculum was quantified by RT-qPCR [17] to ensure reconstitution was reproducible between trials (Table 2).

### Monitoring

Rabbits were fitted with cat collars comprising a 3D-accelerometer-based activity monitor (a wearable device designed for animals) and a temperature datalogger (Figure 1).

**Figure 1.**
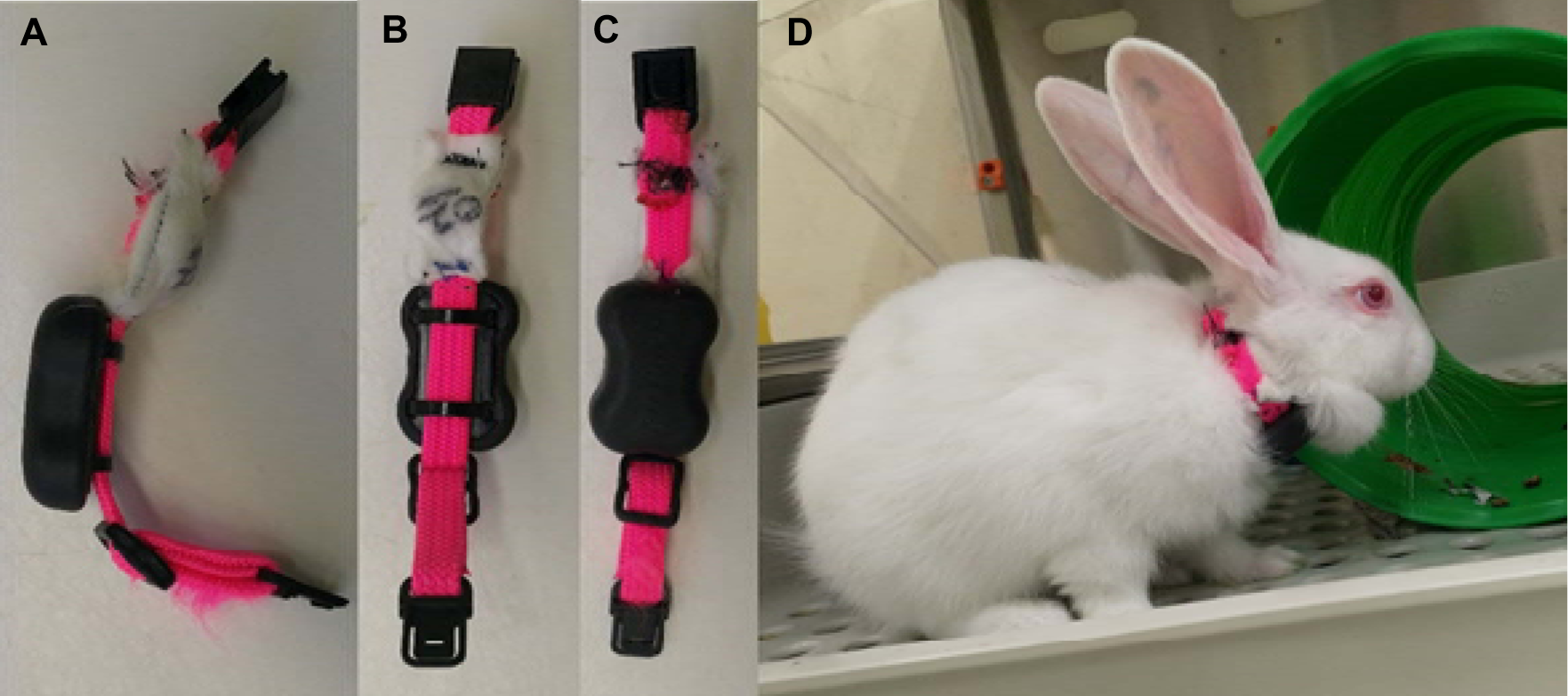
Continuous temperature and activity monitoring. A cat collar was fitted for each rabbit, which comprised a SubCue-Mini temperature datalogger within a fabric pouch and a 3D accelerometer to measure activity levels. A) Side view; B) Skin-apposing view; C) External-facing view; D) Collar fitted on rabbit.

In trials 1 to 3, Tuokiy pet fitness trackers and the associated Joyful Pet app (Zhuhai Tuokiy Electronics, China) were used, which generated a reading every 20 minutes. Due to technical issues, no activity data were recorded for trial 4. Activity trackers were subsequently switched to FitBark2 devices and the associated mobile and web app (FitBark Inc, Kansas City, USA) for trials 5 to 8. The FitBark2 monitors generated an activity reading every minute. SubCue-Mini temperature dataloggers (Canadian Analytical Technologies Inc, Alberta, Canada) were placed in a thin gauze pouch affixed to the collar and held firmly in place against the dewlap to record surface body temperature. Since we were interested in the trend in temperature over time (rather than the absolute core body temperature) surgical implantation of these devices was considered unnecessarily invasive. Temperature and activity monitoring was initiated at least two days prior to infection to obtain baseline recordings for each individual. Death as an endpoint was not used in this experiment; instead, humane endpoints were established to minimise disease and suffering while still observing as complete a disease course as possible to evaluate the welfare impacts of infection. Rabbits with terminal disease, most obviously assessed by a rectal temperature less than 38°C with concurrent lethargy or those experiencing seizures, were humanely killed by intravenous barbiturate overdose after sedation with xylazine 5mg/kg and ketamine 30 mg/kg administered intramuscularly. Trials were terminated once humane endpoints were reached or at 10 days post-infection (dpi). Despite these intervention criteria, due to the acute nature of rabbit haemorrhagic disease some infected animals died peracutely between monitoring timepoints. To account for this, each rabbit was monitored continuously using an Annke 8CH wireless 1080P HDMI NVR 2500TVL camera (Annke, California, USA) with infrared night vision to obtain a precise time of death and to observe any clinical signs prior to death.

### Sample collection

The temporal dynamics of viraemia were monitored by quantifying viral RNA in whole blood samples (collected into EDTA tubes) at the following time points: time of infection, 1 dpi, 1.5 dpi, 2 dpi, 2.5 dpi, 3 dpi, 4 dpi, 7 dpi, 8 dpi, 9 dpi, and at necropsy. Serum was collected prior to infection and at necropsy to investigate early antibody responses to infection. Liver samples were collected at necropsy to estimate virus RNA load in the liver at endpoint.

### Laboratory analyses

RNA was extracted from 50–100 mg of EDTA blood or 20 mg of liver using the SimplyRNA Tissue kit and Maxwell RSC16 instrument (Promega, Wisconsin, USA), as per the manufacturer’s instructions. This protocol was previously validated on whole blood samples in our laboratory. Virus loads in blood and liver were determined using a previously described universal RT-qPCR assay targeting a conserved region of VP60 and utilising a standard curve for absolute quantification [17]. Blood glucose measurements were obtained at the time of blood sampling using an Accu-chek handheld glucometer (Roche Diabetes Care, NSW, Australia). GI.2-specific IgM responses were estimated in pre-infection and necropsy serum samples using a previously described assay [41].

### Data analyses

SubCue-Mini temperature data, Tuokiy activity data, and Fitbark2 activity data were downloaded from their respective applications and analysed in Microsoft Excel and R version 4.0.3 [43]. Plots were generated using ggplot2 [44] and ggpubr [45]. Survival analyses were conducted in survminer [46]. Other R packages used during this study for data manipulation and statistical analyses include readxl [47], plyr [48], pracma [49], emmeans [50], cowplot [51], RColorBrewer [52], scales [53], and those in the tidyverse [54].

## Results

To experimentally assess the virulence and disease progression of RHDV2 in naïve rabbits 5-week-old (kitten) and 11-week-old (adult) New Zealand White laboratory rabbits were orally infected with either a high (1000 RID_50_) or low (50 RID_50_) dose of a GI.1bP-GI.2 RHDV2. We also orally inoculated four control rabbits for each age group with sterile distilled water. We observed a 100% CFR in all infected rabbits regardless of age or infectious dose, while all the control animals survived until the end of the experiment (10 dpi) (Figure 2A).

**Figure 2.**
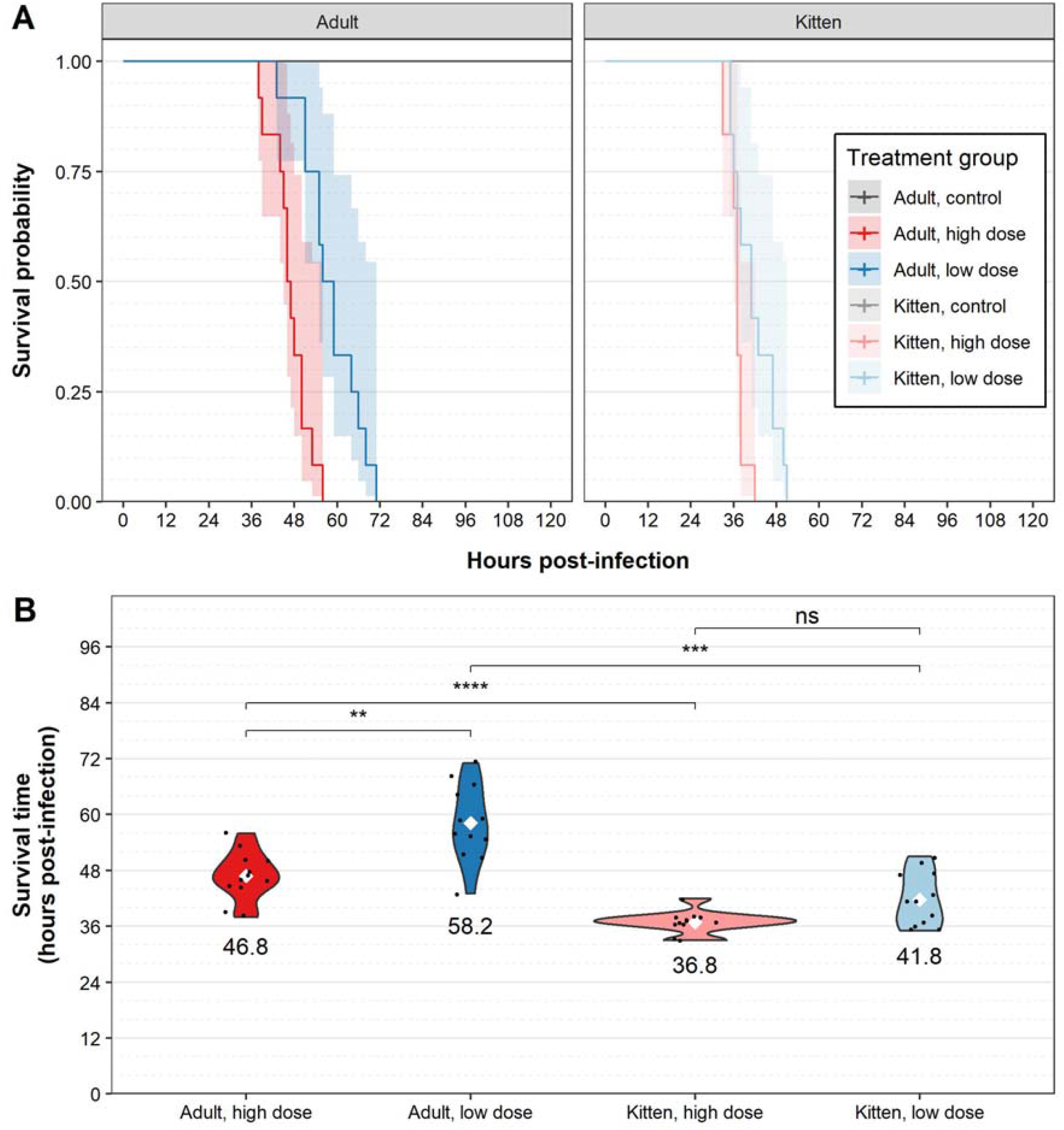
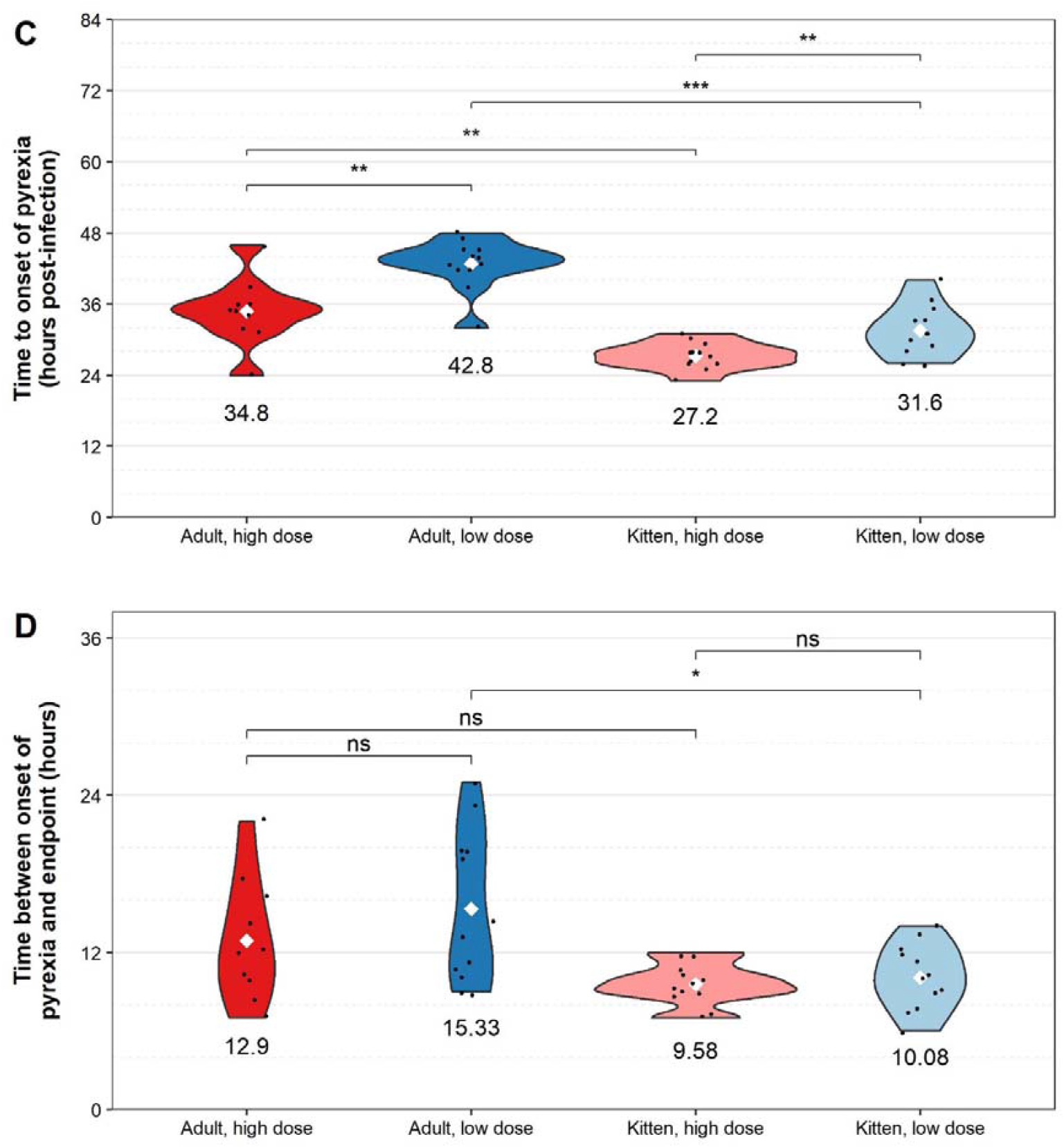
Survival and pyrexia following RHDV2 challenge. Adult (11-week-old) rabbits and kittens (5-week-old) were challenged with either a high virus dose (1,000 RID_50_; *n* = 12 per age) or a low virus dose (50 RID_50_; *n* = 12 per age), or monitored as uninfected controls (*n* = 4 per age). The precise survival time after virus challenge was derived from continuous video camera monitoring. (A) Survival analysis was performed using survminer. Transparent shaded areas represent 95% confidence intervals. Plots are right-censored at 120 hours post-infection; all control animals survived until the end of the experiment (i.e., 10 days post-infection). (B) Survival time was derived by combining data from time to euthanasia (humane endpoint) and time to death obtained from continuous camera monitoring. (C) Time to onset of pyrexia after RHDV2 challenge was estimated based on plots from continuous temperature monitoring collars. (D) The duration of clinical disease was taken to be the time between the onset of pyrexia and endpoint. Violin plots were constructed for each treatment group. Individual data points are shown as black dots. The mean for each treatment group is shown as a white diamond and the value of the mean (in hours) is given below each plot. Note that temperature collars did not record properly for two animals in the ‘adult, high dose’ group so *n* = 10 for this group in panels C and D. Significance was calculated using a two-sided Wilcoxon signed rank test in ggpubr. ns not significant, * p <= 0.05, ** p <= 0.01, *** p <= 0.001, **** p <= 0.0001.

The mean survival time following infection was 46 hours, and infected animals died or reached humane endpoints between 33 and 71 hours post infection (hpi). Kittens had a significantly shorter survival time (range 33–51 hpi [x□ 39 hpi]) than adult rabbits (range 38–71 hpi [x□ 53 hpi]) infected with the same virus dose (Figures 2A&B, Table 3). Additionally, within the adult rabbit groups, survival time was significantly shorter in rabbits infected with a high virus dose (range 38–56 hpi [x□ 47 hpi]) than in those infected with a low virus dose (range 43–71 hpi [x□ 58 hpi]) (Figures 2A&B, Table 3). This dose-dependent difference in survival time was not significant in kittens; notably, the effect size was small in this age group relative to the population variance (Figures 2A&B, Table 3).

**Table 3.** Statistical analyses of disease progression after RHDV2 infection. Significance testing was conducted using the two-sided Wilcoxon signed rank test. All p values are given to two significant figures. The effect size refers to the difference between the means of the comparison groups.

| Comparison groups | p | Effect size (hrs) | Standard deviation (hrs) |  |
| --- | --- | --- | --- | --- |
|  |  |  | Group 1 | Group 2 |
| *Survival time* |  |  |  |  |
| High dose, adult vs kitten | 0.000079 | 10.0 | 5.2 | 2.4 |
| Low dose, adult vs kitten | 0.00013 | 16.4 | 8.1 | 5.8 |
| Adults, high dose vs low dose | 0.0015 | 11.4 | 5.2 | 8.1 |
| Kittens, high dose vs low dose | 0.062 | 5.0 | 2.4 | 5.8 |
| *Time to onset of pyrexia* |  |  |  |  |
| High dose, adult vs kitten | 0.0015 | 7.6 | 5.6 | 2.2 |
| Low dose, adult vs kitten | 0.00015 | 11.2 | 4.2 | 4.3 |
| Adults, high dose vs low dose | 0.0050 | 8.0 | 5.6 | 4.2 |
| Kittens, high dose vs low dose | 0.0089 | 4.4 | 2.2 | 4.3 |
| *Duration of clinical signs of disease* |  |  |  |  |
| High dose, adult vs kitten | 0.059 | 3.3 | 4.7 | 1.6 |
| Low dose, adult vs kitten | 0.030 | 5.2 | 5.7 | 2.4 |
| Adults, high dose vs low dose | 0.35 | 2.4 | 4.7 | 5.7 |
| Kittens, high dose vs low dose | 0.60 | 0.5 | 1.6 | 2.4 |

Infected rabbits exhibited clinical signs such as pyrexia, lethargy, weight loss, and terminal seizures. Pyrexia (observed as a clear increase from baseline temperature using collar temperature monitors [Supplementary figures—red line]) was the most consistently observed clinical sign, developing between 23–48 hpi in all infected individuals. In line with the trends observed for survival time, the mean time to onset of pyrexia (estimated from the collar temperature monitors [Supplementary figures—red line]) was significantly shorter in kittens and in those animals infected with a high virus dose (Figure 2C, Table 3). In contrast to the findings for survival time, the effect of virus dose on mean time to onset of pyrexia was statistically significant in both kittens as well as adult rabbits (Figure 2C, Table 3). Pyrexia peaked (based on maximum smoothed conditional mean of collar temperature monitors [Supplementary figures—thick red line]) on average 7 hours prior to endpoint (range 1–17 hours before endpoint) in both adults and kittens. This peak was followed by a continual decline leading to hypothermia and terminating with death or euthanasia of the animal (Figure 3).

**Figure 3.**
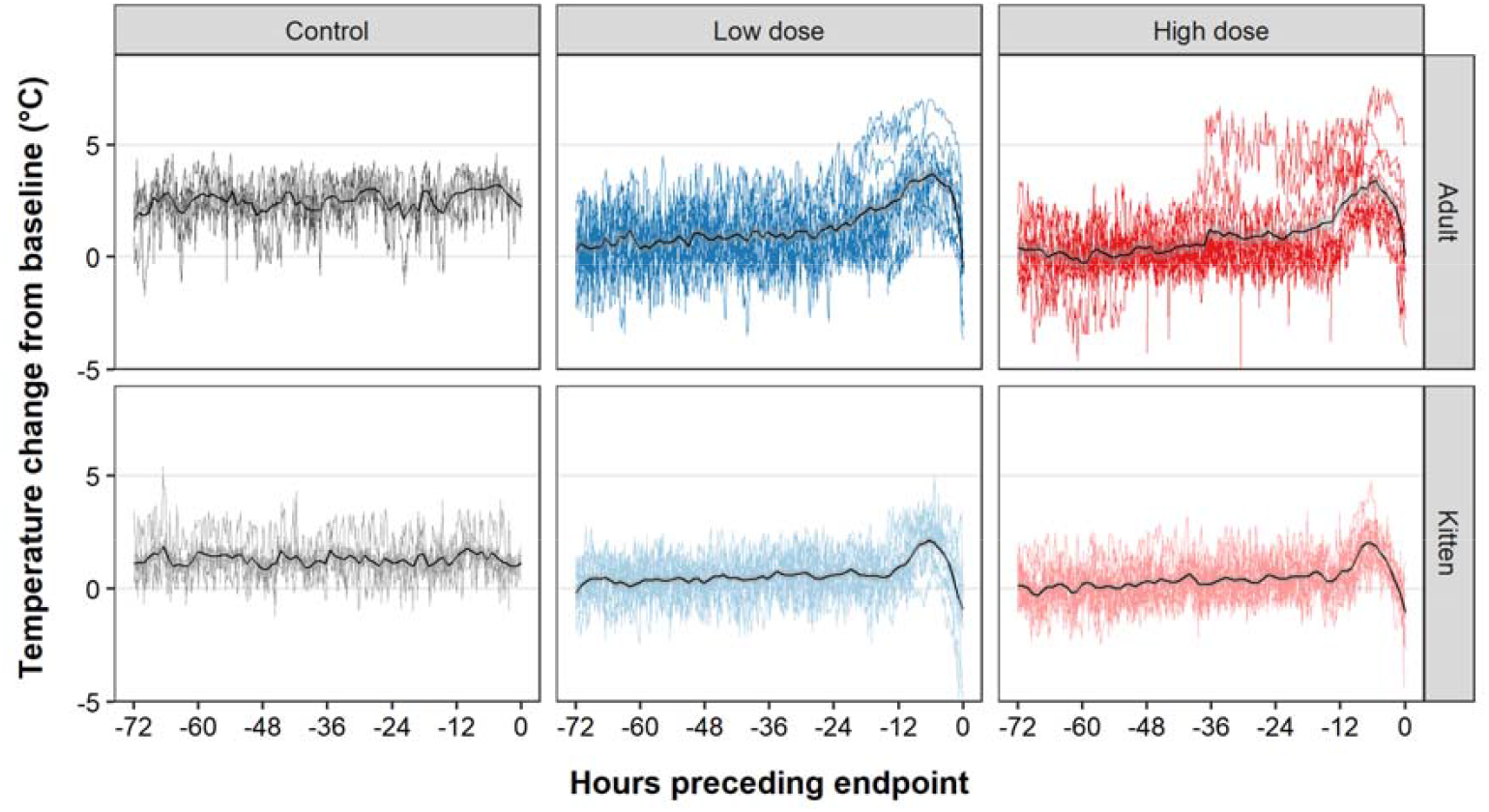
Temperature changes preceding death following RHDV2 infection. For each animal, the baseline temperature was calculated by averaging the temperature monitoring collar readings over a 1-hour period prior to infection. This baseline was then subtracted from each reading to give the temperature change from baseline, and this was plotted against the time preceding endpoint. Data were excluded where technical errors were encountered with the collars (see supplementary figures). Time pre endpoint was used instead of hours post infection because the decline from the peak of pyrexia was very consistent between animals. Plots are faceted by age and treatment group. Overlaid coloured lines represent temperature profiles from individual animals. The solid black lines represent the smoothed conditional mean for each treatment group. This was calculated using local polynomial regression fitting as implemented in ggplot2 (geom_smooth(method=“loess”) with span = 0.05). Transparent grey shaded areas represent 95% confidence intervals.

Lethargy was observed both through subjective monitoring and through objective monitoring using activity trackers (Supplementary figures). There was substantial noise in the activity tracker data and complete datasets were not obtained for all animals; however, generally, control animals maintained regular cyclical activity patterns over the duration of the experiment while infected animals showed a gradual reduction in activity levels coincident with the onset of pyrexia (Supplementary figures—orange line). This was most obvious in kittens, where fewer technical difficulties with the activity monitors were experienced. Notably, the reduction in activity was only evident in the final hours of disease (Supplementary figures—orange line).

Although we observed fluctuations in bodyweight in some individuals, weight loss was typically observed in infected animals, while in contrast, uninfected rabbits tended to gain weight over the duration of the experiment (Figure 4, Supplementary figures—green line). Of the 48 infected rabbits, weight loss was identified in 65% prior to endpoint, weight loss was only identified at the final monitoring point in 33% of animals (i.e., at a humane endpoint or after the rabbit had died), and weight loss was not observed in one kitten (Supplementary figures—green line). The percent weight loss (from maximum bodyweight) experienced in infected rabbits ranged from 0–7%, averaging 3% in kittens and 4% in adults. Weight loss was first observed between 24–66 hpi in adults and between 29–48 hpi in kittens, typically after the onset of pyrexia (Figure 4, Supplementary figures—green line).

**Figure 4.**
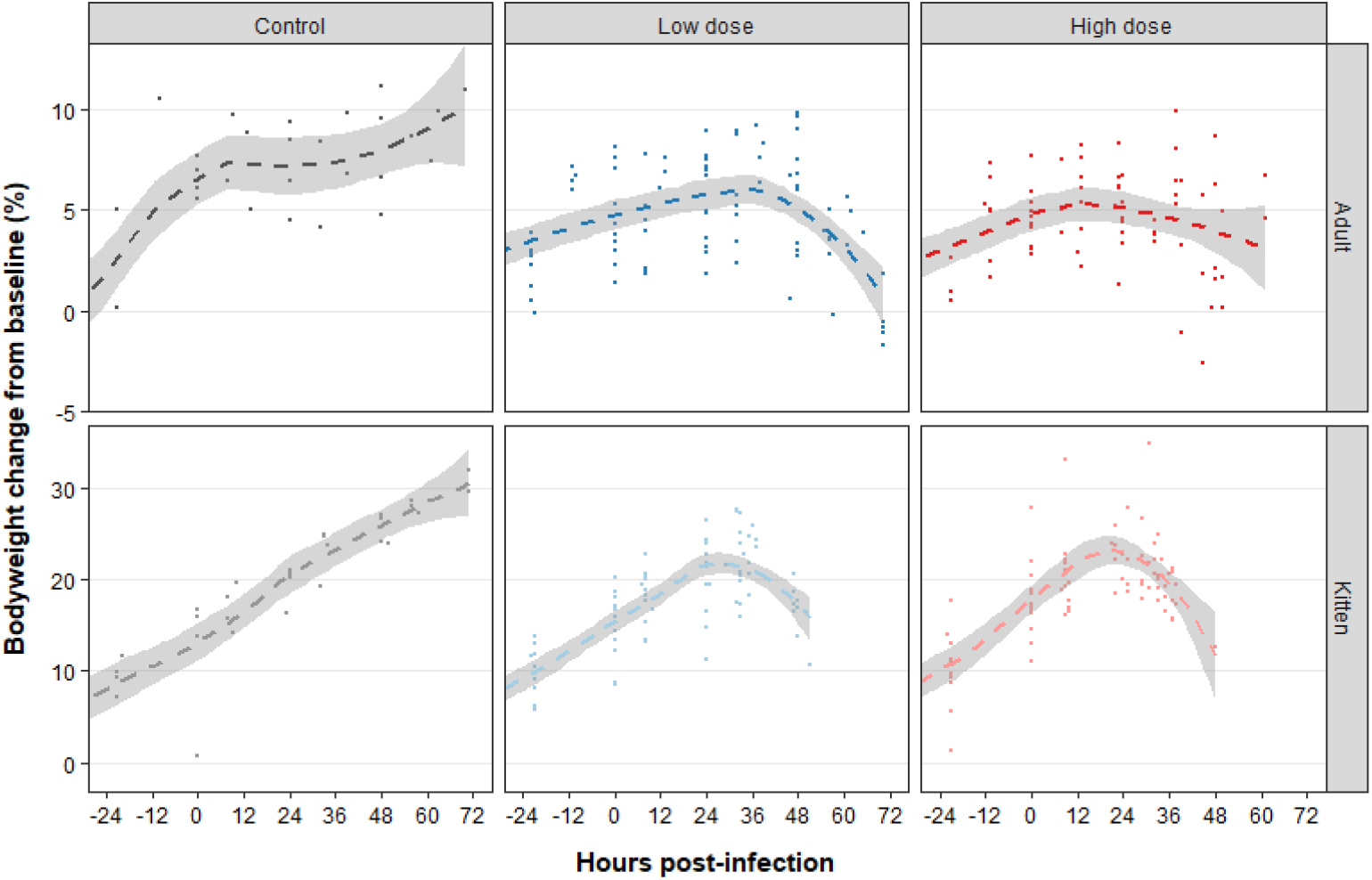
Bodyweight changes following RHDV2 infection. Bodyweight was recorded at least daily until endpoint, commencing 24 hours prior to virus challenge. Plots are right-censored at 72 hours post-infection (i.e., after all infected animals reached humane endpoints or died). The change in bodyweight from baseline (in percent) at each measurement point and for each animal is shown as an individual dot. Plots are faceted by age and treatment group. The dashed lines represent the smoothed conditional mean for each treatment group. This was calculated using local polynomial regression fitting as implemented in ggplot2 (geom_smooth(method=“loess”) with default span). Transparent shaded areas represent 95% confidence intervals. Note that the y-axis scales differ for the two age groups.

For animals that died between monitoring points, the video footage was reviewed to look for additional clinical signs in the terminal phase of disease. Seizures were observed in all infected animals that died (*n* = 30). These were characterised as intermittent episodes of generalised tonic-clonic seizure activity and commenced between 1 hour to 5 minutes prior to death. To further investigate the pathological mechanism precipitating these seizures, we obtained paired blood glucose measurements from 10 infected and 3 control kittens at the time of infection and at necropsy. Blood glucose concentration is known to decrease rapidly after death due to continued cellular glucose consumption; therefore, we restricted blood glucose measurements to animals that were humanely killed, and measurements were acquired immediately after barbiturate overdose. We observed post-mortem hyperglycemia in the three control animals when compared to baseline, which was presumably induced by the xylazine/ketamine combination used for sedation (Figure 5). This anaesthetic combination is known to affect glucoregulatory hormones, leading to decreased levels of serum insulin and subsequent secondary increases in blood glucose [55]. In contrast, all infected rabbits showed a decrease in blood glucose concentration between the paired baseline and post-mortem readings (Figure 5). Importantly, these animals were not yet experiencing seizures. Blood glucose measurements at necropsy in infected animals immediately after euthanasia ranged from 2.6–7.0 mmol/L (xJ 4.3 mmol/L). Notably, these measurements are likely to be artificially elevated due to the administration of the xylazine/ketamine sedation, as observed in the control animals.

**Figure 5.**
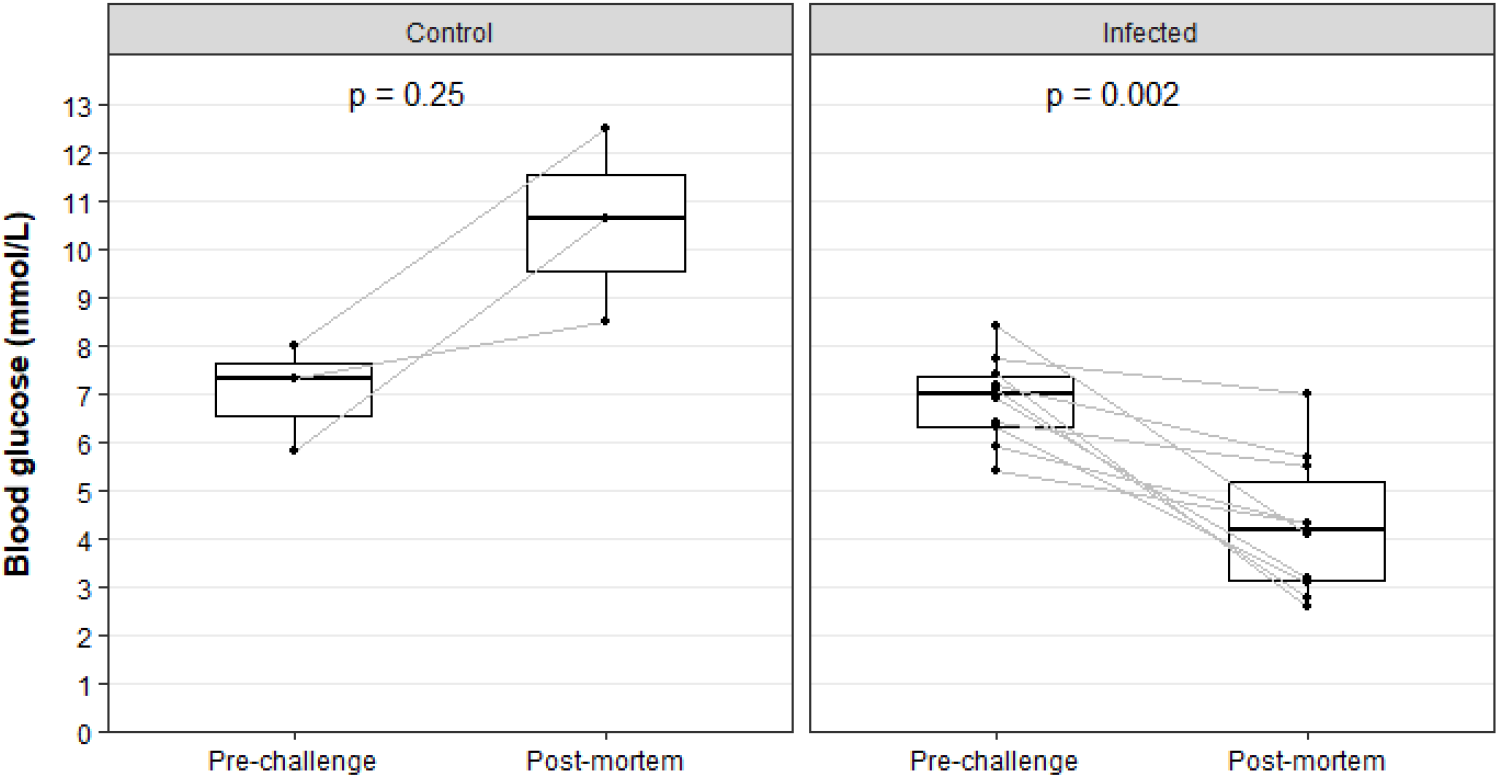
Blood glucose concentration in kittens pre- and post- RHDV2 infection. Blood glucose concentration was measured in a subset of kittens using an ‘Accu-chek Performa’ glucometer before infection and at post-mortem. Data were restricted to those animals that were euthanised (*n* = 13 kittens). Paired box plots were generated for each timepoint. Individual data points are shown as black dots, and paired samples are linked by grey lines. Significance was tested using paired, two-sided Wilcoxon signed rank tests as implemented in ggpubr, and exact p-values are given.

To investigate the welfare impacts of RHDV2 infection, we determined the duration of clinical disease in infected animals (i.e., the time during which clinical signs were detectable). Since pyrexia was 1) typically the first clinical sign to be observed (Supplementary figures), 2) was reliably observed in all infected animals, and 3) was continuously measured via the collar loggers, we considered welfare impacts to be experienced from the time of onset of pyrexia to the time of death/humane endpoint. Kittens experienced welfare impacts for 6–14 hours after RHDV2 infection (Figure 2D). Consistent with the longer disease duration observed in adults, welfare impacts were experienced for a duration of 7–25 hours in older rabbits (Figure 2D). Importantly, the gradual decline in activity that we observed suggests that welfare impacts were mild at the time of onset of pyrexia and increased in severity as disease progressed (Supplementary figures—orange line).

To study the *in vivo* replication kinetics of the GI.1bP-GI.2 RHDV2 variant, we monitored the temporal dynamics of viraemia after infection. Sporadic low-level RT-qPCR contamination was detected in some uninfected control and baseline samples (Figure 6). However, infected animals showed an obvious exponential increase in viral RNA in blood over the course of infection (Figure 6). Viraemia was detectable by 24 hpi in most infected animals (Supplementary figures—blue line). At this 24 hpi time point all kittens infected with a high virus dose were viraemic, correlating with the rapid disease progression observed in these animals. Viraemia peaked in the post-mortem samples, reaching an average of 1×10^8^ capsid gene copies per mg blood (range 3×10^5^–2×10^9^). The viral RNA load was significantly lower in blood than in the liver (range 5×10^6^–7×10^8^ [x□ 2×10^8^]) at the same time point (Figure 7A). Virus load in the liver (at necropsy) was significantly higher in kittens compared to adults (Figure 7B); there was no significant difference between animals that received a high vs low infectious dose (two-sided Wilcoxon signed rank test, p = 0.48).

**Figure 6.**
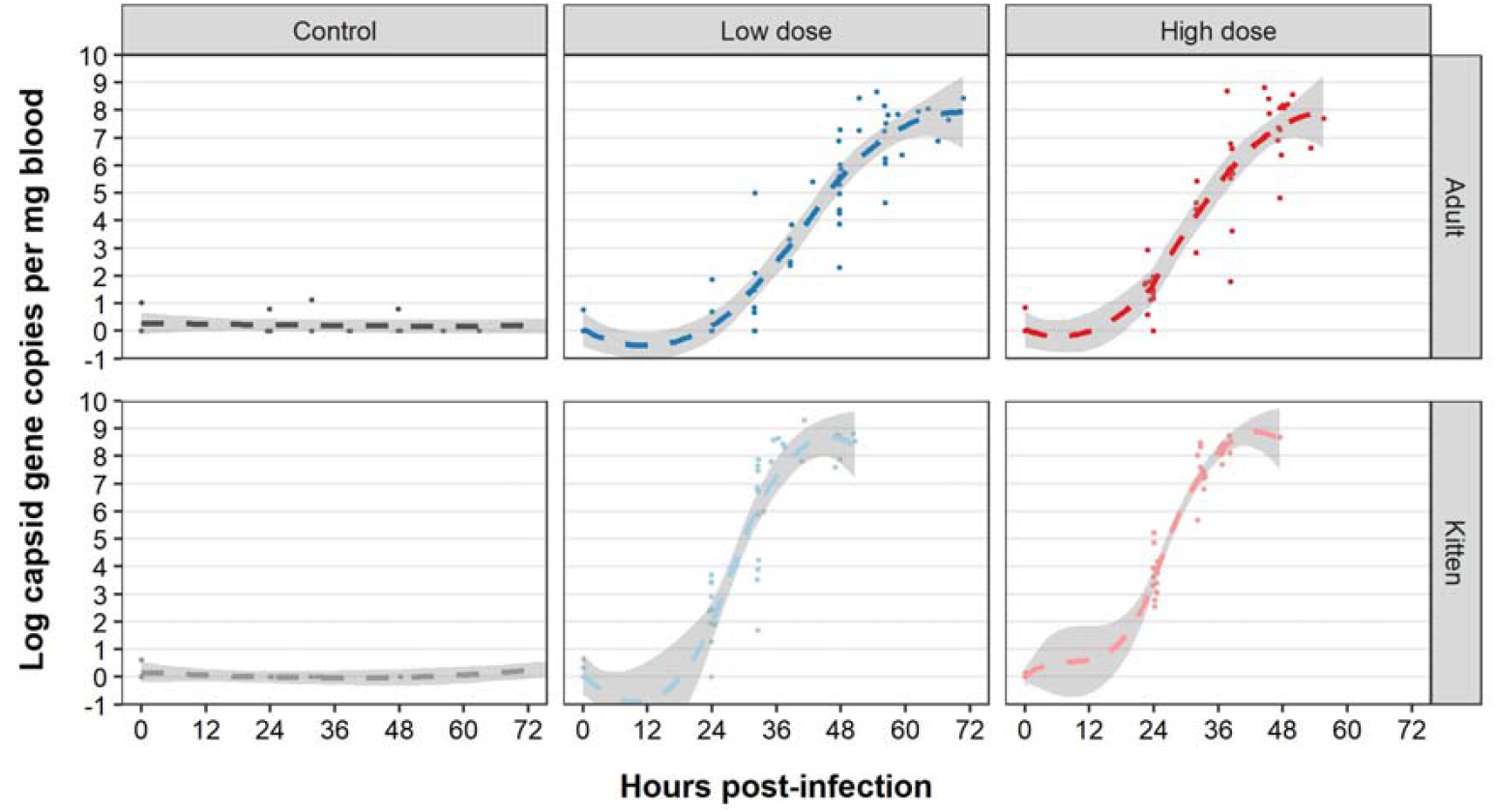
Kinetics of viraemia after RHDV2 challenge. Blood samples were collected at 0- and 24-hours post-infection and up to twice daily thereafter. Total RNA was extracted from whole blood and virus capsid gene copies per mg blood were quantified by RT-qPCR. Individual measurements are shown as dots. Plots are faceted by age and treatment group. The dashed lines represent the smoothed conditional mean for each treatment group using local polynomial regression fitting as implemented in ggplot2 (geom_smooth(method=“loess”) with default span). Transparent shaded areas represent 95% confidence intervals. Plots are right-censored at 72 hours post-infection for control animals.

**Figure 7.**
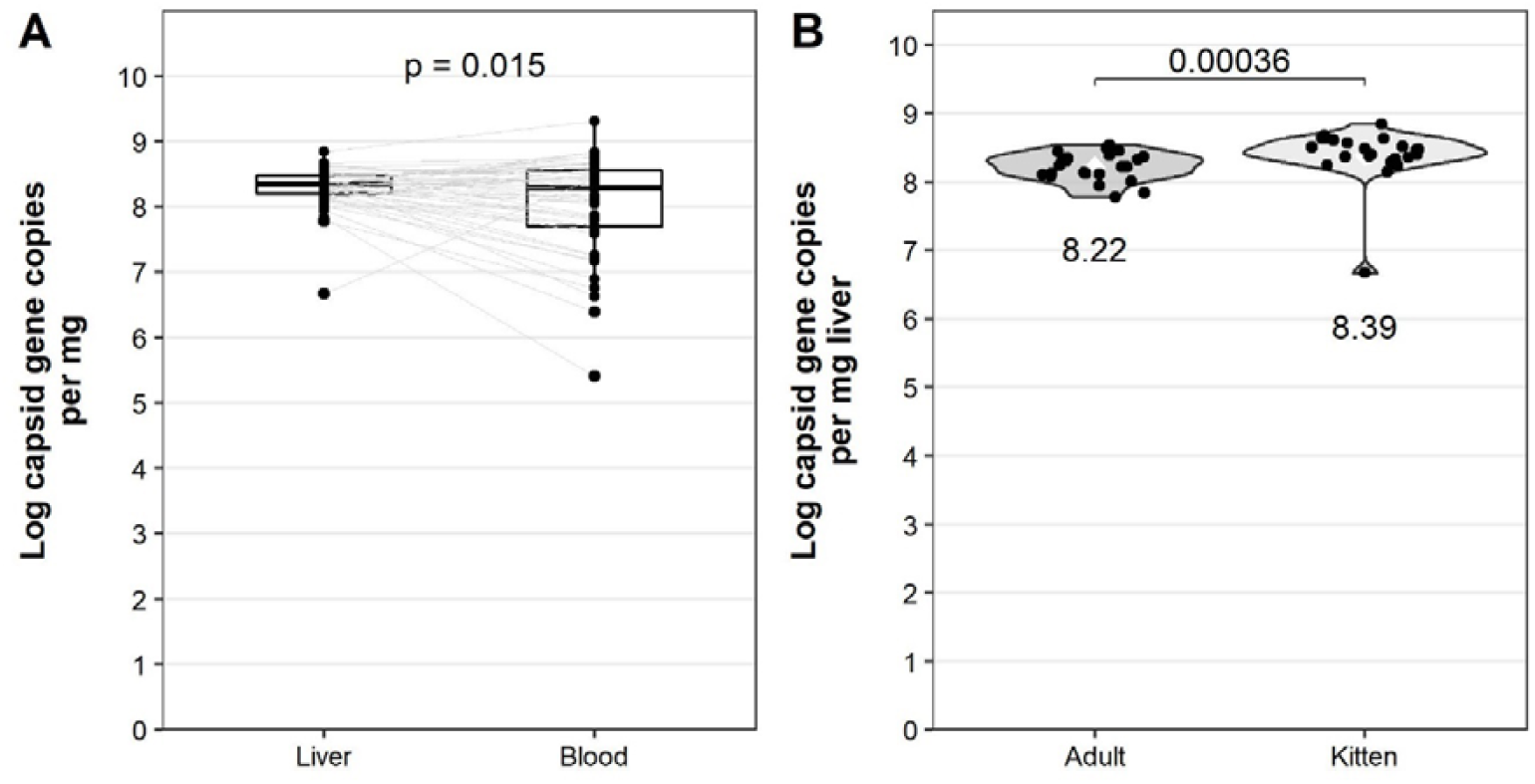
Virus loads in the liver and blood after RHDV2 infection. Total RNA was extracted from whole blood and liver post-mortem samples and log_10_ virus capsid gene copies per mg blood were quantified by SYBR-based RT-qPCR. Significance was tested using paired, two-sided Wilcoxon signed rank tests as implemented in ggpubr, and the p-value is given. (A) Paired box plots were generated to compare post-mortem virus loads in the liver and whole blood. Individual data points are shown as black dots, and paired samples are linked by grey lines. (B) Virus loads in the liver were plotted as violin plots and compared between adult rabbits and kittens. Individual data points are shown as black dots. The mean virus load in the liver for each group is shown as a white diamond, and the value of the mean is given below each plot.

Despite the rapid progression of disease, infected animals showed a significant increase in RHDV2-specific IgM reactivity in post-mortem sera when compared to paired pre-infection sera (Figure 8A). This IgM response was not detected in control animals. IgM was detectable as early as 36 hours post-infection in one animal (kitten, high dose; Figure 8B). The magnitude of IgM reactivity did not correlate with either age or infectious dose (two-sided Wilcoxon signed rank test, p = 0.30 for age and p = 1.00 for infectious dose). A simple linear regression model showed a significant positive correlation between IgM reactivity and survival time (p = 0.011; Figure 8B)). When this association was investigated for each age and infectious dose, the relationship was only significant for adult rabbits infected with a low dose (p = 0.0022, R^2^ 0.63). Notably, this group also had the longest survival times.

**Figure 8.**
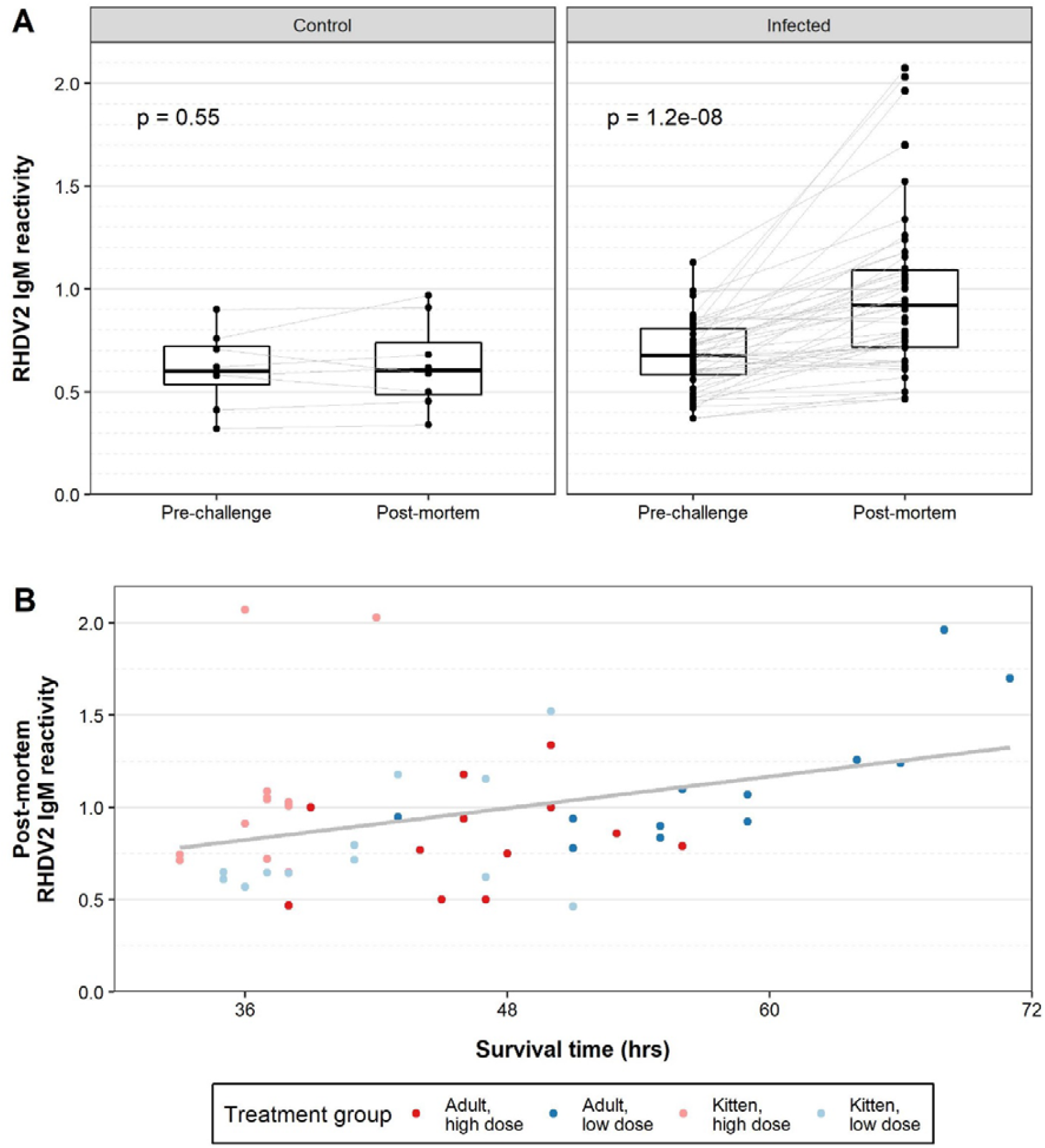
RHDV2-specific IgM reactivity after RHDV2 challenge. RHDV2-specific IgM reactivity was determined in pre-challenge and post-mortem sera by isotype-specific ELISA. Reactivity was calculated as the optical density of the sample at 1/40 dilution divided by the optical density of the negative control serum at 1/40 dilution. (A) Paired box plots were generated pre- and post-challenge. Individual data points are shown as black dots, and paired samples are linked by grey lines. Significance was tested using paired, two-sided Wilcoxon signed rank tests as implemented in ggpubr, and exact p-values are given. (B) The post-mortem RHDV2-specific IgM reactivity of infected rabbits was plotted against survival time. A simple linear regression model was plotted in grey.

## Discussion

The pathological changes observed at post-mortem examination after RHDV2 infection have been widely reported elsewhere [30, 33, 34]. In contrast to several previous studies, we found RHDV2 to be highly virulent in naïve laboratory rabbits. In our experiments, the CFR was 100% in both adult rabbits and kittens, regardless of infectious dose. We suggest that the discrepancies in the observed virulence of RHDV2 described in the existing literature are likely due to the different virus variants used for challenge studies. Here we used a GI.1bP-GI.2 virus that is widely circulating in Australia, the Iberian peninsula, the Azores, Morocco, France, and Poland [12, 15, 16, 18, 19, 35-38]. A previous study using this variant also reported a 100% CFR in laboratory rabbits [34], while studies reporting lower CFR using GI.3P-GI.2 variants or variants where the NS genotype was unknown [14, 28, 30, 32, 33].

The importance of recombination in generating genetic diversity in lagoviruses [19, 56], and the fitness benefits conferred by the NS proteins [57], are only beginning to be understood, and functional analyses remain challenging without a robust cell culture system. It is critically important for future RHDV2 pathogenesis and epidemiological studies that the full variant genotype is specified.

Disease progression was influenced by both age and infectious dose. We found that kittens and animals infected with a high virus dose succumbed to terminal disease faster than adult rabbits or those infected with a low virus dose. This trend was also reflected in the time to onset of pyrexia and in the progression of viraemia. Other studies have also suggested that disease severity may be increased in kittens. Dalton, Balseiro [33] observed that lesions appeared to be more frequent and more severe in kittens and that viraemia was detectable earlier in kittens than in adults. Interestingly, we also found that the virus load in the liver at death was significantly higher in kittens compared to adults. These results emphasise the importance of controlling for confounding factors, such as age and infectious dose, when conducting virulence comparisons between lagoviruses.

Haemagglutination (HA) units were used to quantify the amount of virus present in the inoculum in previous challenge studies, however HA units are a measure of antigen concentration and can only indirectly indicate infectious dose. Since there is currently no robust cell culture system for lagoviruses, accurate determination of the infectivity of virus stocks must be performed by titration in rabbits. The finding that disease progression is indeed affected by infectious dose highlights that titration of the inoculum is necessary to accurately compare disease phenotype between variants. Moreover, strict biosecurity measures are required to prevent cross-contamination between animals, which may inadvertently increase the infectious dose given. In this study the control animals remained uninfected, confirming that cross-infection did not occur.

We observed that the onset of pyrexia typically correlated with an exponential increase in viraemia, which frequently coincided with the onset of weight loss and lethargy. Weight loss in diseased animals could be due to decreased feed intake or could be an indicator of dehydration, however, the speed at which weight loss occurred may suggest that dehydration was more likely. Hydration status was not assessed objectively in this study. Transient weight loss that was not associated with disease was also seen in many animals (6/8 control animals); this was observed during the acclimatisation period or immediately after infection. We suggest that this weight loss is due to the stress of transport, initially, and then extended handling at the infection time point. Since weight loss was readily triggered by stress, we found that pyrexia was a more reliable indicator of infection.

Viraemia was detectable by RT-qPCR in all infected animals, and was first detected between 24 and 48 hpi, depending on age and infectious dose. This suggests that ante-mortem diagnostic testing for RHDV2 is technically feasible; however, given the rapid disease progression, RT-qPCR testing is likely to be initiated too late in disease for effective interventions to be administered. For ante-mortem diagnostic purposes, point-of-care rapid antigen tests using serum or whole blood would perhaps be more useful than RT-qPCR, because of their faster turnaround time. However, the analytical sensitivity of these assays would be critical, particularly early in infection when viraemia is low, based on the viraemia profiles observed in this study. Whether treatment modalities, such as hyperimmune antiserum [58], non-nucleoside inhibitors [59], or other antivirals, would be effective after diagnosis still needs to be investigated. RHDV2-specific IgM reactivity was detectable in some infected animals despite survival times of less than 71 hours in adults and 51 hours in kittens. Importantly, the serological response at these early time points was low and was only detectable through paired sampling, confirming that serology is not useful as a diagnostic tool during acute infection.

After RHDV2 infection we observed non-specific clinical signs that increased in severity over the course of disease. In contrast to previous reports, a subacute disease course was not observed in this study, even after low dose infection [14]. Specifically, we observed pyrexia that progressed to hypothermia (a finding consistent with end stage distributive shock), weight loss (likely through dehydration), and lethargy. Using video camera monitoring, we also observed terminal seizures in all rabbits that were not humanely killed (i.e., died between monitoring points). These seizures were associated with hypoglycaemia. Whether the hypoglycaemia precipitated the seizures (as a consequence of fulminant hepatitis and septic shock) or whether the seizures were triggered by other pathophysiological processes such as hepatic encephalopathy, with subsequent utilisation of glucose during seizure activity, was not determined in this study.

However, previous studies on RHDV1 have reported hypoglycaemia from as early as 24–36 hpi [60, 61], preceding the seizure activity that was only observed terminally in this study. Notably, the duration of welfare impacts (the time from onset of pyrexia to endpoint) ranged from 6–14 hours in kittens and 7–25 hours in adult rabbits. Since many rabbits were humanely killed according to predefined intervention criteria, this may be slightly underestimated; however, this duration of welfare impacts is remarkably short for a pathogen or toxin. For example, pindone poisoning causes functional impairments related to internal haemorrhage for up to 7 days [62].

The use of pet activity trackers, continuous temperature monitors, and continuous video monitoring facilitated detailed observations into the pathogenesis and welfare impacts of RHDV2. While instrumented observation cages and implantable telemetry devices are becoming more readily available, they are not commonly used in rabbits. For example, the Phenocube (Psychogenics, New Jersey, USA) uses home-cage operant conditioning modules and computer vision software to evaluate cognitive performance, motor activity, and circadian rhythms of group-housed mice, but is not compatible with rabbit housing systems [63]. The UID Mouse Matrix system (Unified Information Devices, Illinois, USA) is an RFID-enabled system that allows real-time recording and monitoring of location, movement, and body temperature for multiple animals within a single cage using subcutaneously implanted microchips. While this system appears to be suitable for rabbits, it has only recently entered the market. Telemetry devices such as the PhysioTel Digital L series of implants (Data Sciences International, Minnesota, USA) continuously measure several physiological variables simultaneously, such as blood glucose, body temperature, blood pressure, respiratory parameters, heart rate, and/or activity.

These devices have been used in viral pathogenesis studies, such as studies of influenza virus H5N1 in cynomolgus macaques [64], eastern equine encephalitis virus in mice [65], Ebola virus in rhesus macaques [66], Rift Valley fever virus in African green monkeys and common marmosets [67], and simian immunodeficiency virus in rhesus macaques [68]. However, these devices must be surgically implanted, which is highly invasive. These devices are typically reserved for studies on risk group 3 and 4 pathogens for which routine animal handling poses a considerable risk to handlers. In contrast, the activity trackers, temperature monitors, and video cameras used in this study are non-invasive, relatively inexpensive, and readily accessible, while still providing continuous measurements of key physiological variables.

## Conclusions

Using continuous activity, temperature, and video monitoring, we found that age and infectious dose significantly affected disease progression after RHDV2 infection in naïve laboratory rabbits. Critically, future lagovirus pathogenesis studies should consider differences in disease phenotype due to age and infectious dose and should report the complete genotype of the variant used. It is interesting to speculate that variants with alternate NS proteins may vary in their virulence. While this study was conducted in naïve laboratory rabbits it will be important for future studies to determine whether disease progression after RHDV2 challenge differs in wild rabbits or in rabbits with pre-existing immunity to different lagoviruses.

## Acknowledgements

We wish to acknowledge Michael Frese, Anthony Rowe, and Nagendra Singanallur for critical revision of the manuscript.

## Author contributions

Conceptualization, RNH and TS; methodology, RNH, TS, AJR, JA, KT, and JD; formal analysis, RNH; investigation, RNH, TK and MP; resources, AJR and TOC; data curation, RNH; writing—original draft preparation, RNH; writing—review and editing, RNH, TOC, AJR, JD and TS; visualization, RNH; project administration, RNH and TS; funding acquisition, RNH and TS. All authors have read and agreed to the published version of the manuscript.

## Ethics statements

The study was conducted according to the ‘Australian code for the care and use of animals for scientific purposes’ and was approved by the ‘CSIRO Wildlife, Livestock and Laboratory Animal’ animal ethics committee (permit #2018-06).

