## Supplementary figures for "Age and infectious dose significantly affect disease progression after RHDV2 infection in naïve domestic rabbits"

**Disease course in individual animals.** Bodyweight (green points and green dashed line) was recorded at least twice daily. Each animal was fitted with a monitoring collar. A continuous temperature logger was attached to the collar and held tightly against the skin to record external body temperature (thin red line) every 12 minutes. To smooth noise in the temperature data, a smoothed conditional mean (thick red line) was plotted using local polynomial regression fitting as implemented in ggplot2 (geom_smooth(method = ”loess”,se = T) with span = 0.05). An activity tracker was attached to the same collar as the temperature monitor. In trials 1 to 3, we used Tuokiy activity trackers that generated a reading from the 3D accelerometer every 20 minutes. We encountered repeated technical issues with these trackers, particularly after an app update following trial 3. Consequently, no activity data were able to be recovered from trial 4. We subsequently switched to FitBark2 activity monitors for trials 5 to 8, which generated a 3D accelerometer reading every minute. A moving average of the activity data (orange line) was calculated to smooth noise, with a backwards window length of 255 (FitBark) or 16 (Tuokiy), using the pracma package. The accelerometer readings are in arbitrary units for both devices. Blood samples were collected at several timepoints to monitor viraemia levels (blue points and blue dashed line) over time. Total RNA was extracted from whole blood and virus capsid gene copies per mg blood were quantified by RT-qPCR. Anomalies and dropouts in the collar data (external body temperature and activity) occurred for some animals over some time periods as collars became loose or were scratched off. We have removed these dropouts to improve visualisation and have provided additional information in the caveats below each image.

Adult, control

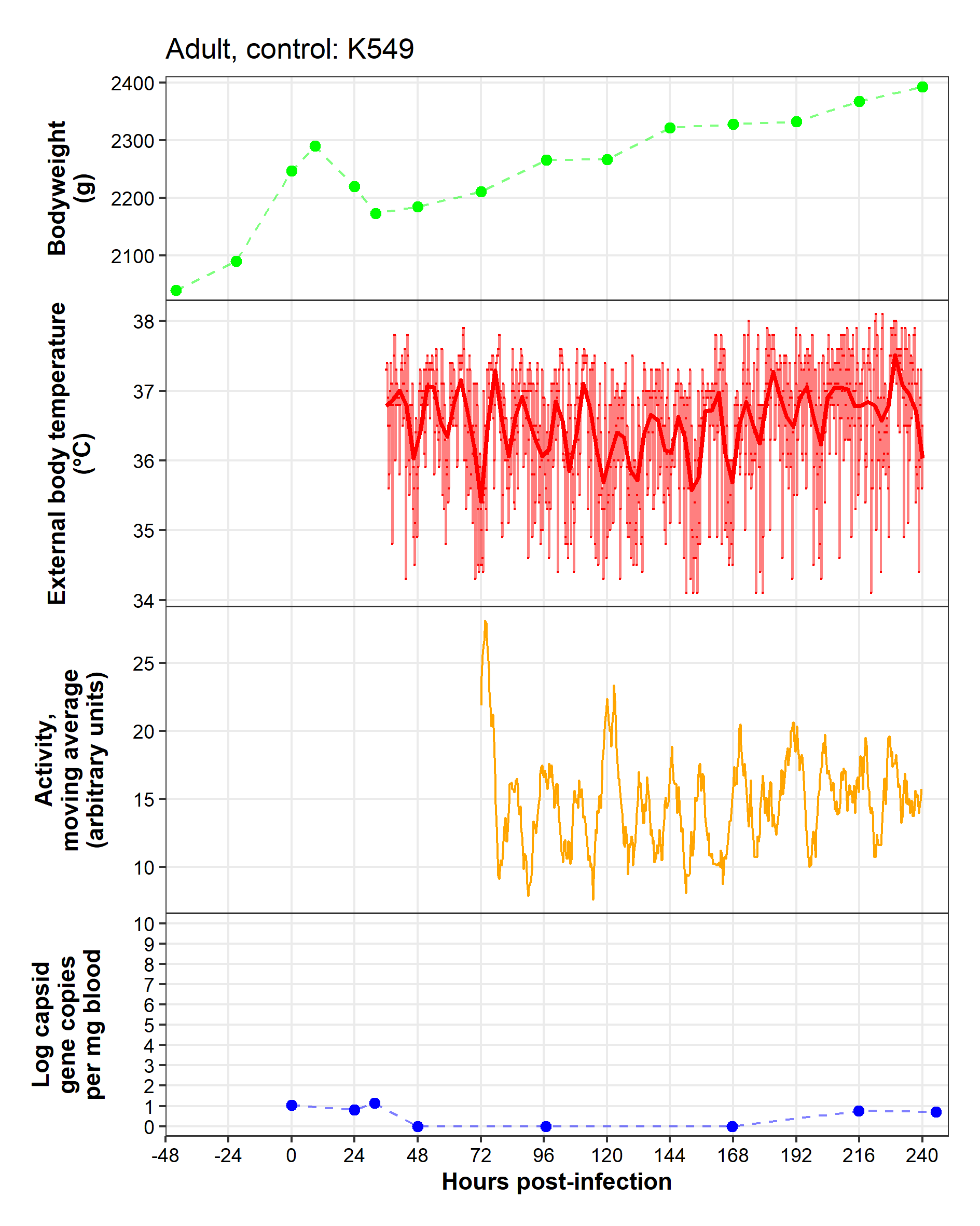

**Caveats**: Temperature monitor fell out of pouch between -48–32 hpi. Experienced technical issues with activity monitor prior to 72 hpi.

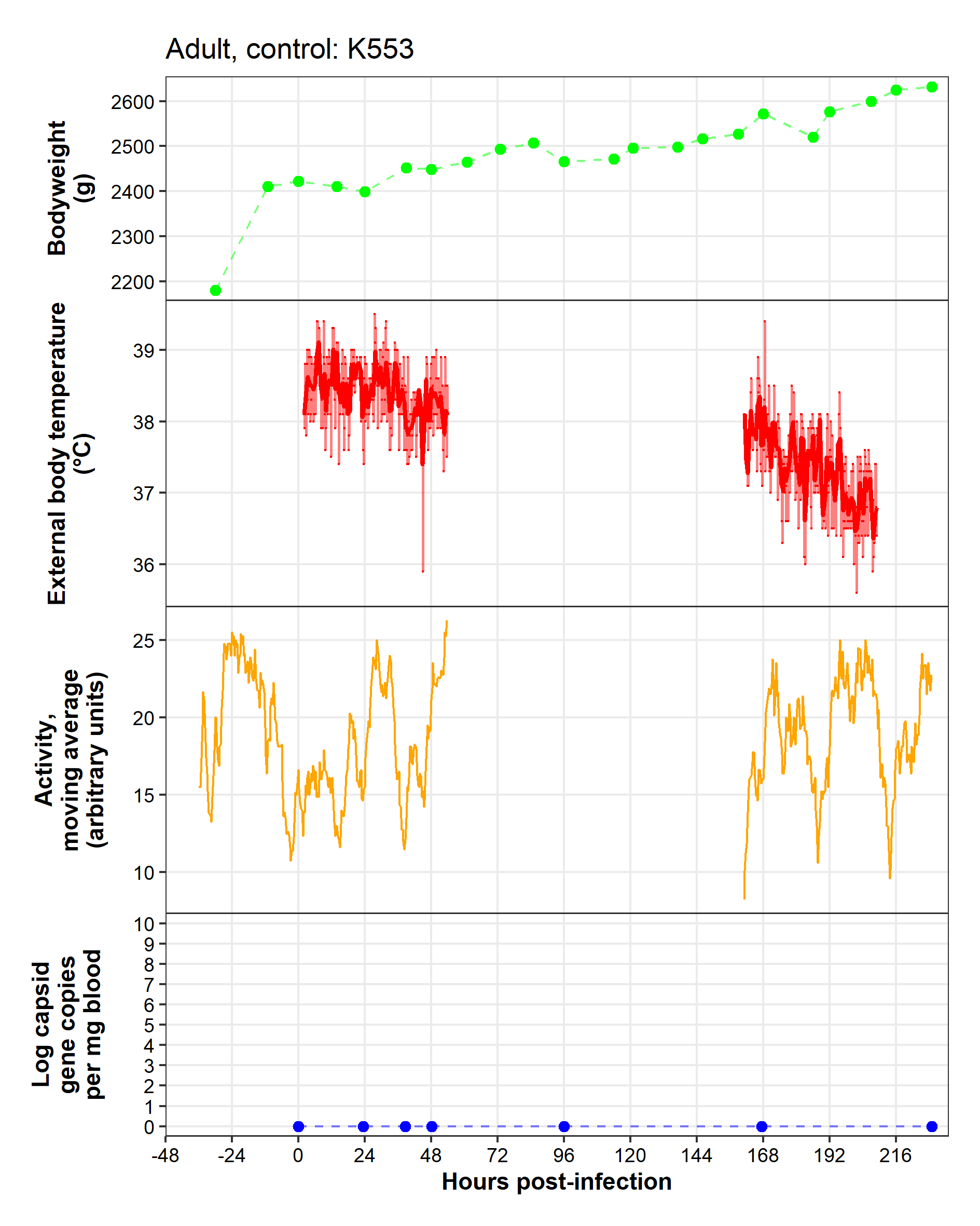

**Caveats:** Temperature monitor fell out of pouch, replaced at 0 hpi. Collar removed between 54–161 hpi. Temperature monitor missing after 209 hpi. Euthanised 229 hpi rather than 240 hpi as per other controls (still 10 dpi).

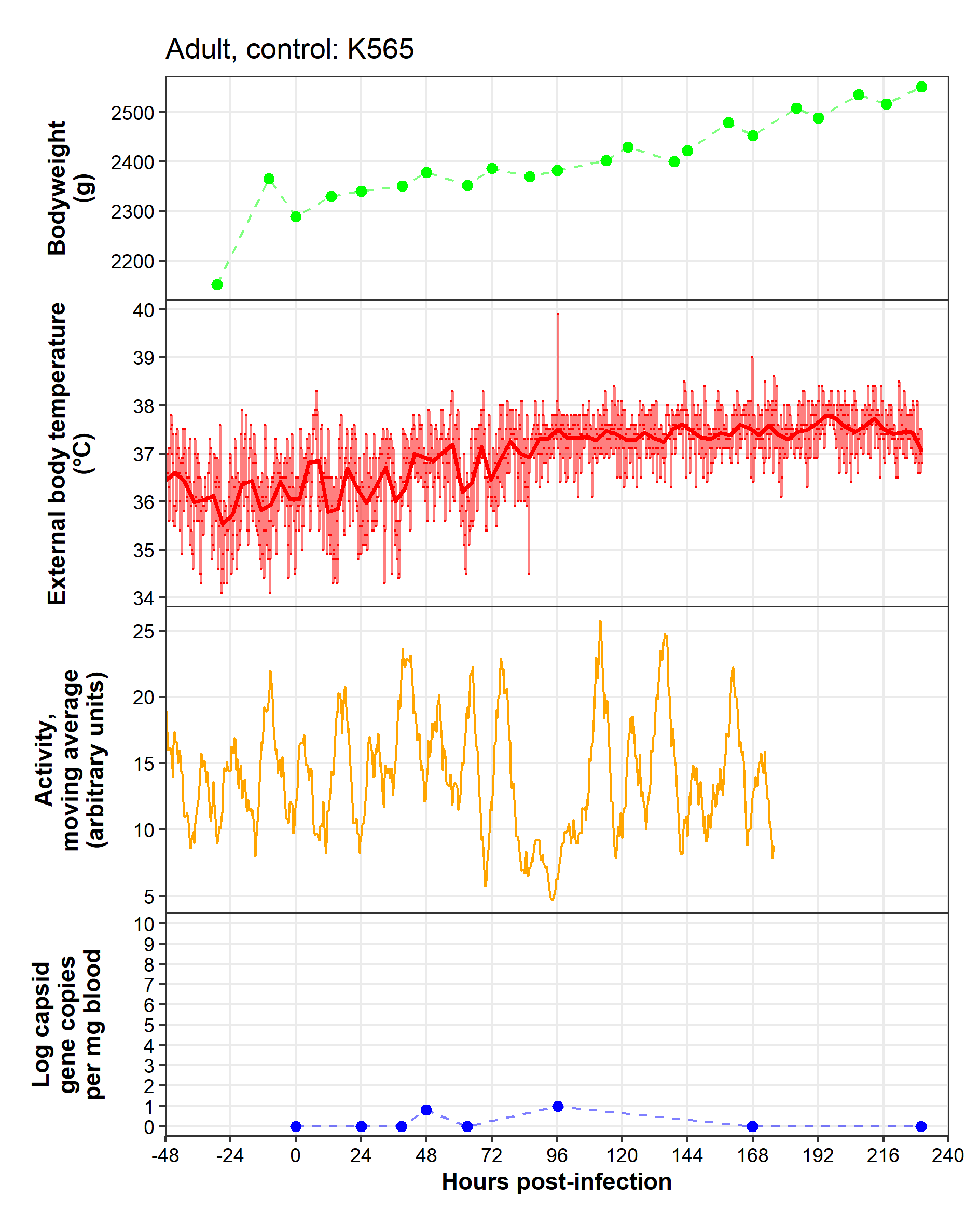

**Caveats:** Technical difficulties with activity monitor after 176 hpi.

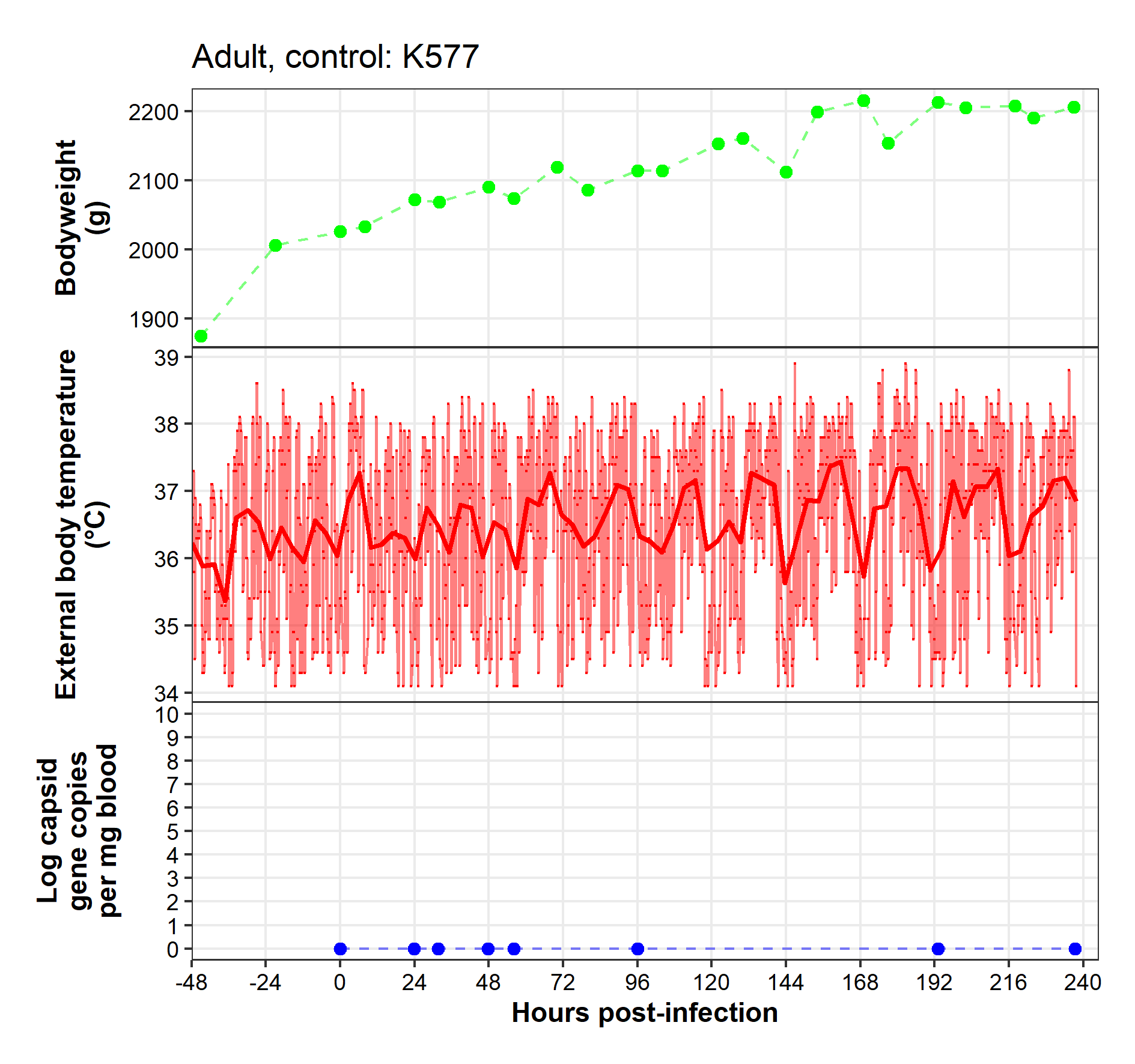

**Caveats:** Activity monitors were not used in trial 4 due to technical issues with the app update.

Adult, high dose

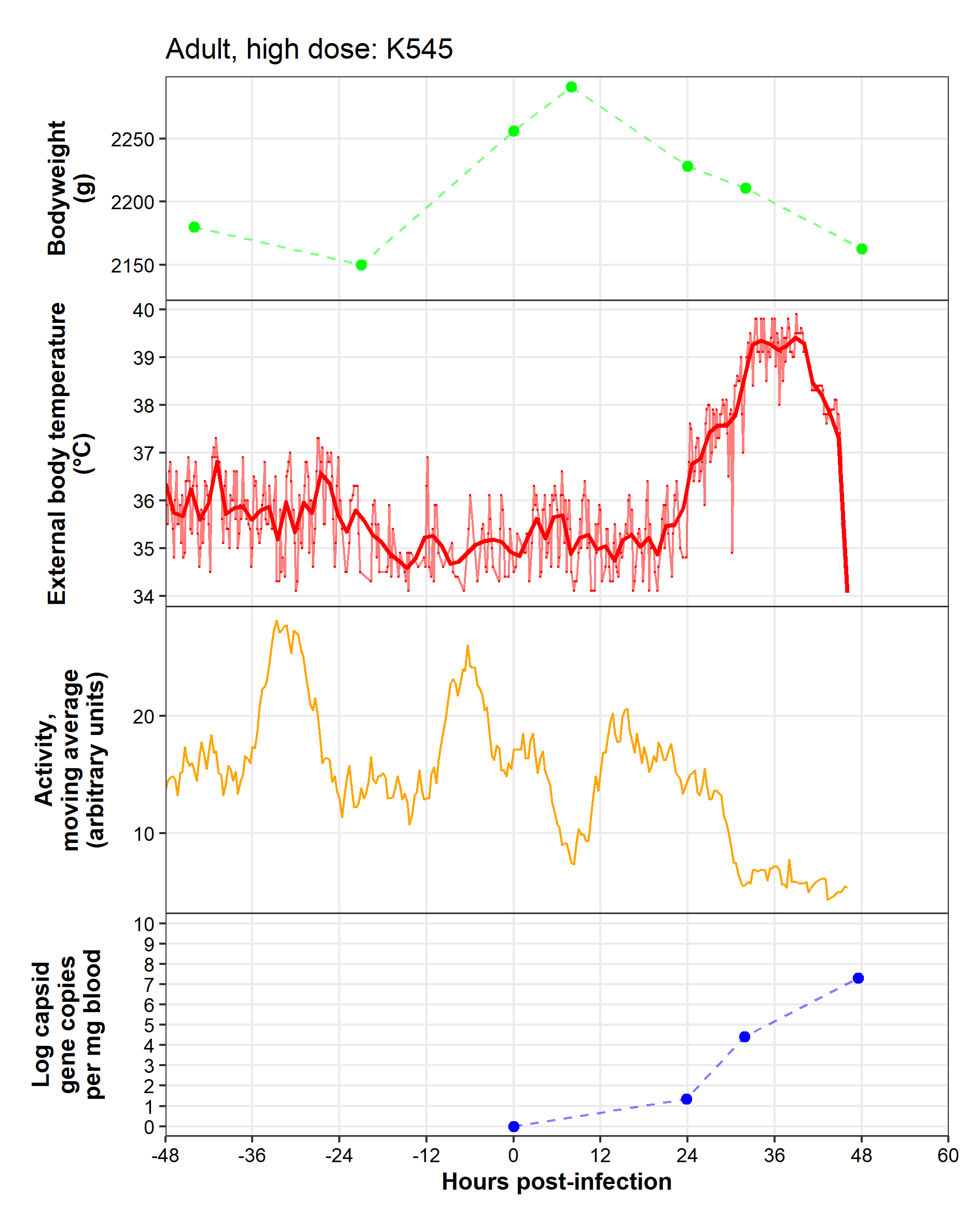

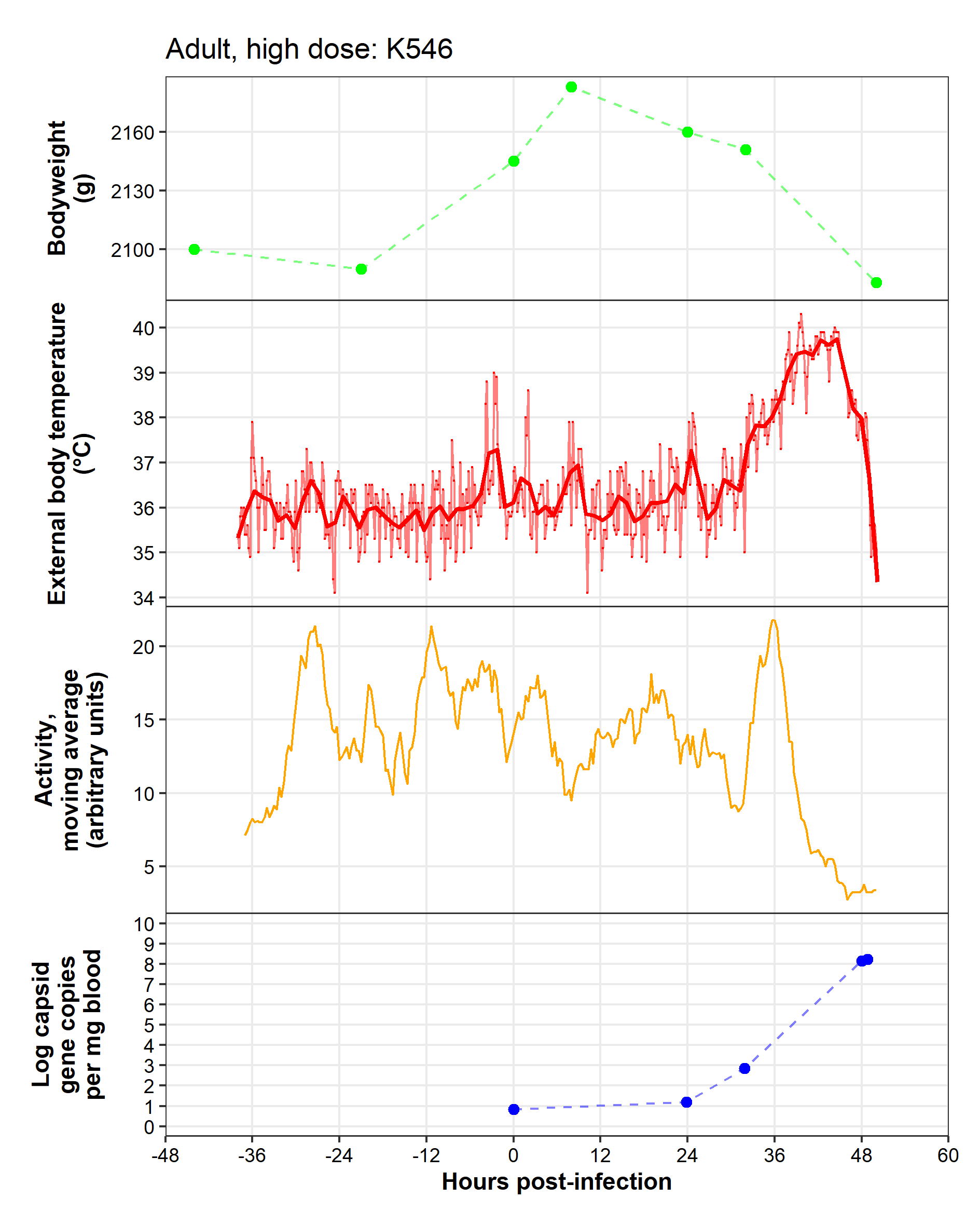

**Caveats:** Technical difficulties with activity monitor before -36 hpi.

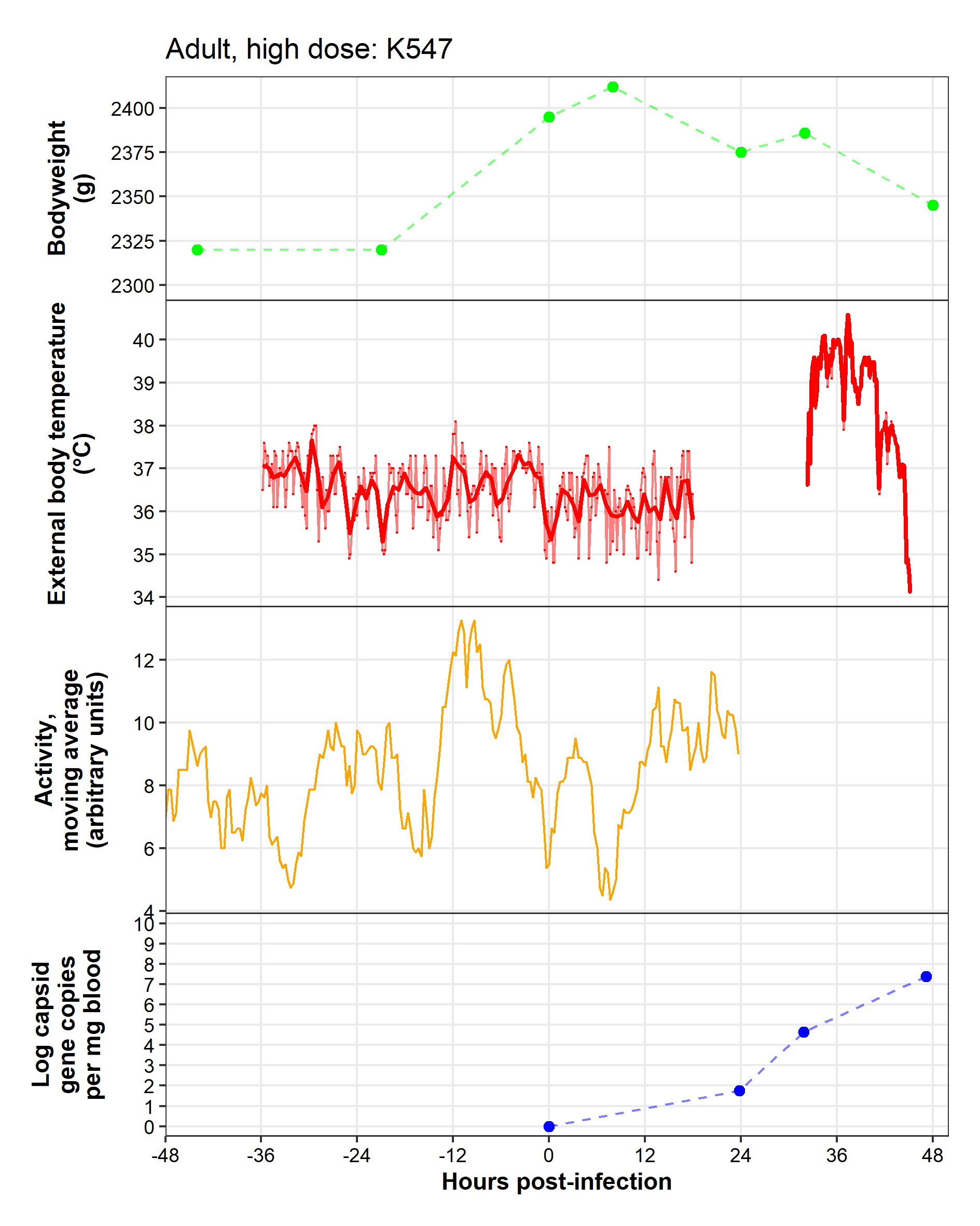

**Caveats:** Technical difficulties with temperature monitor prior to -36 hpi. Collar was twisted so that the temperature logger was not against skin from 18–32 hpi. Technical difficulties with activity monitor after 24 hpi.

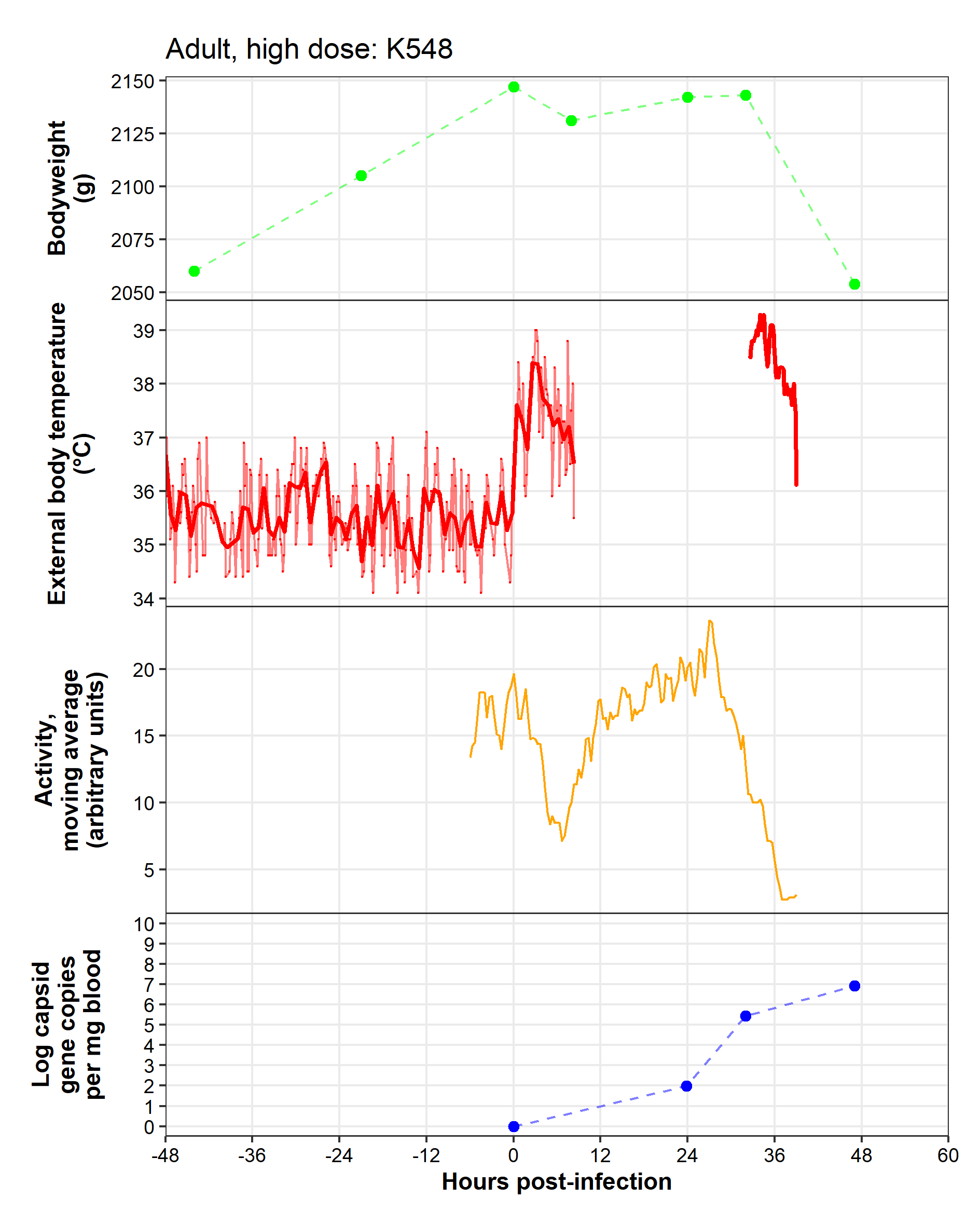

**Caveats:** Temperature monitor fell out of pouch around 10 hpi, replaced at 32 hpi.

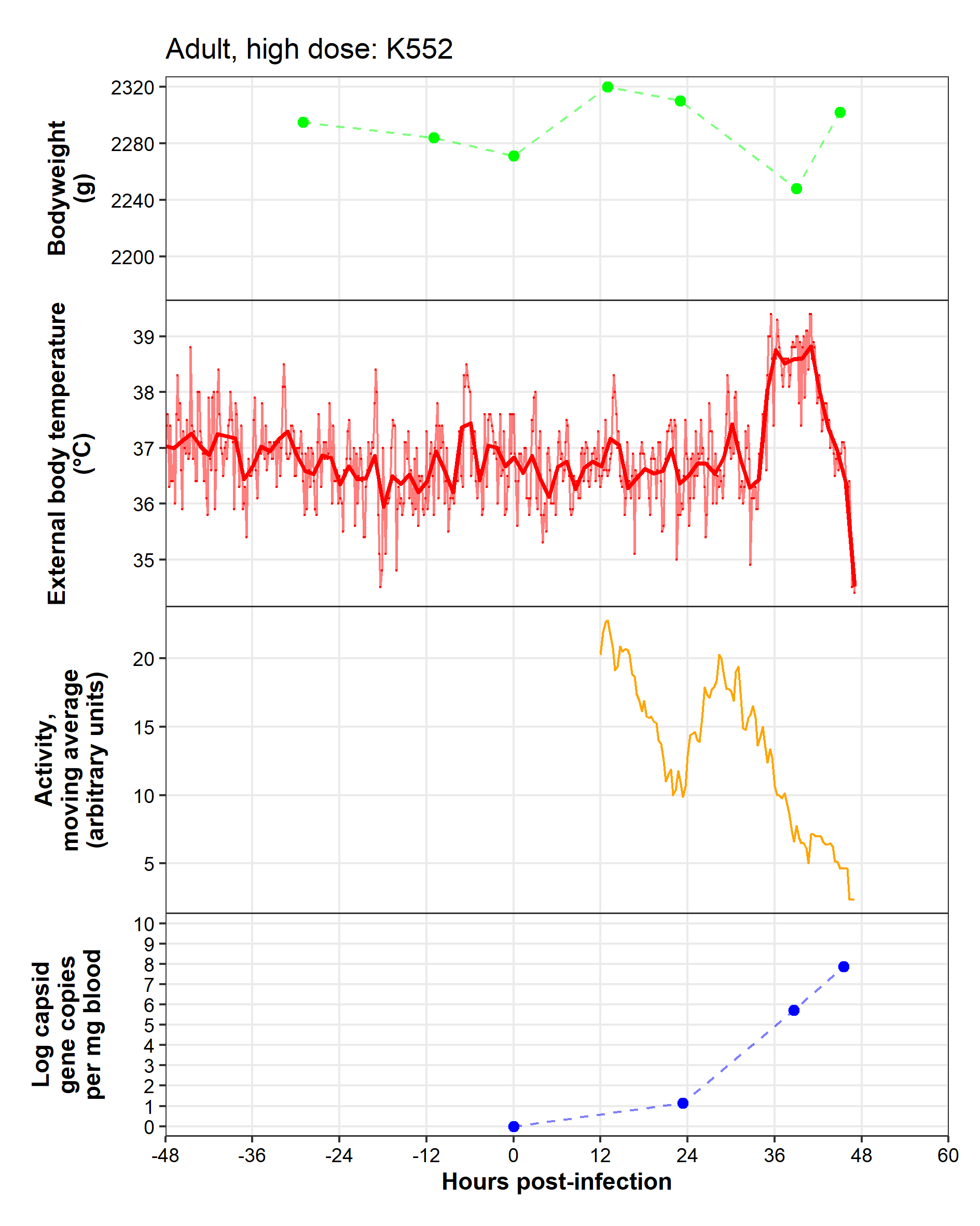

**Caveats:** Technical difficulties with activity monitor prior to 12 hpi.

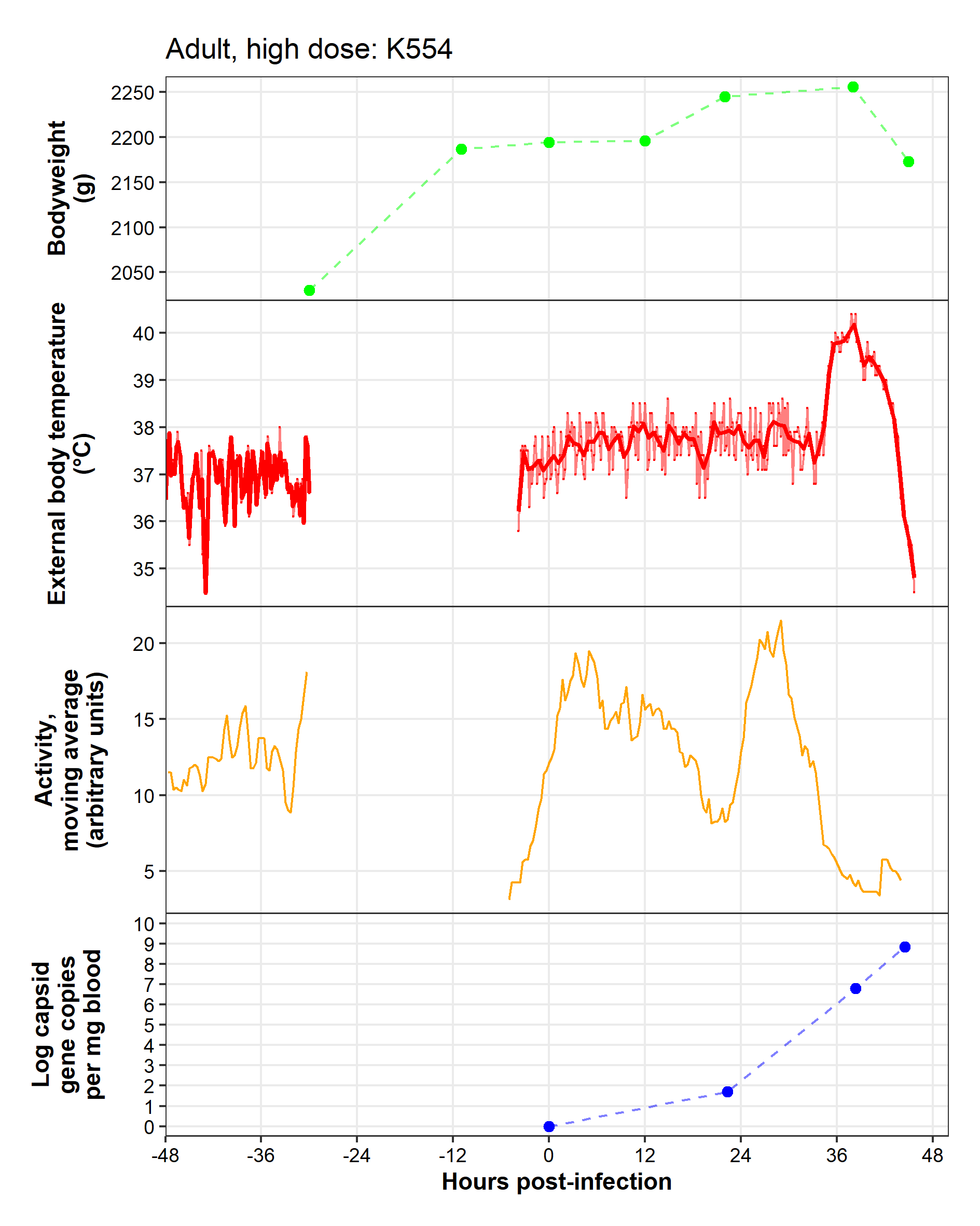

**Caveats:** Collar came off between -30 – -5 hpi.

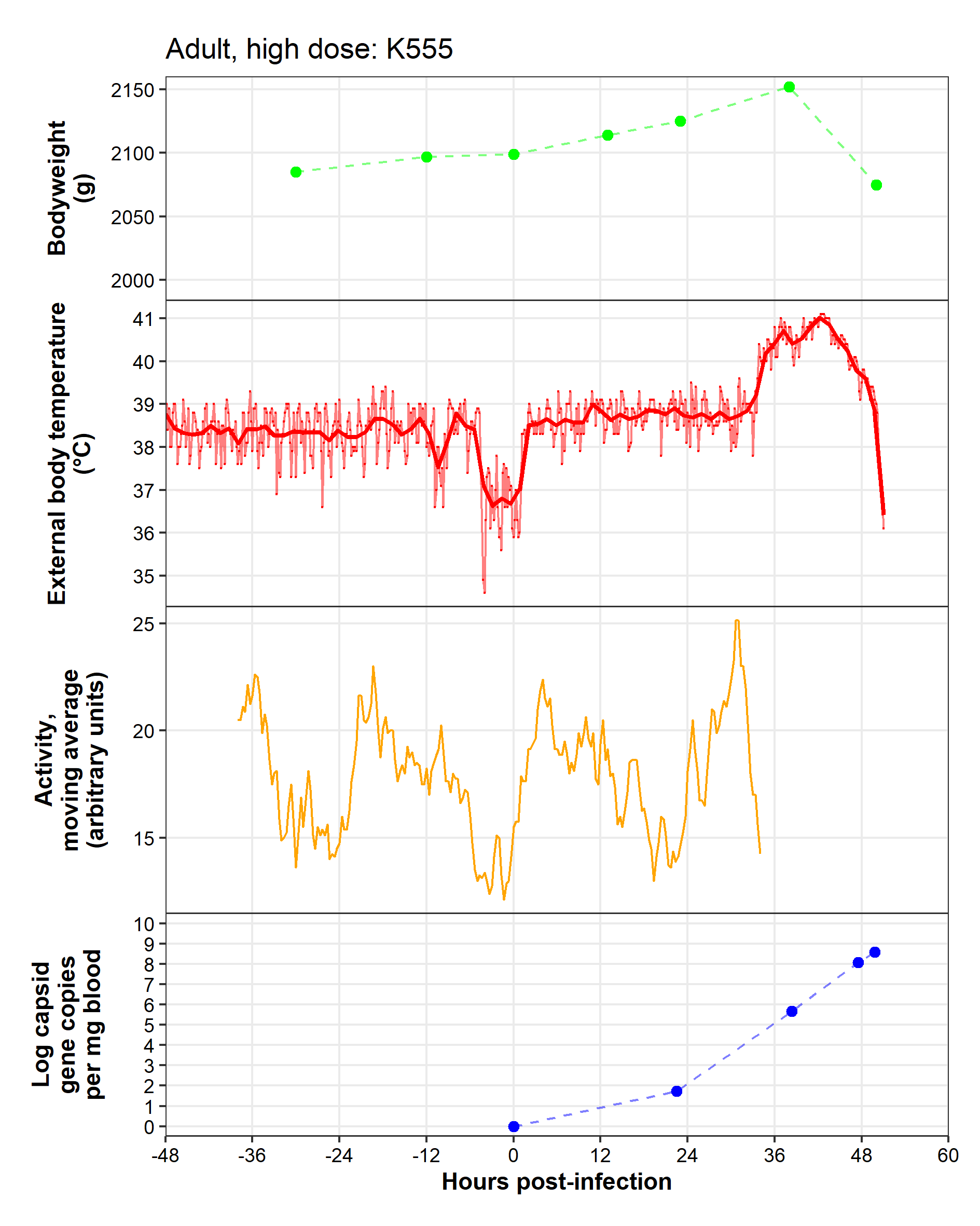

**Caveats:** Technical difficulties with activity monitor after 34 hpi. Temperature monitor replaced at -6 hpi and tightened at 0 hpi.

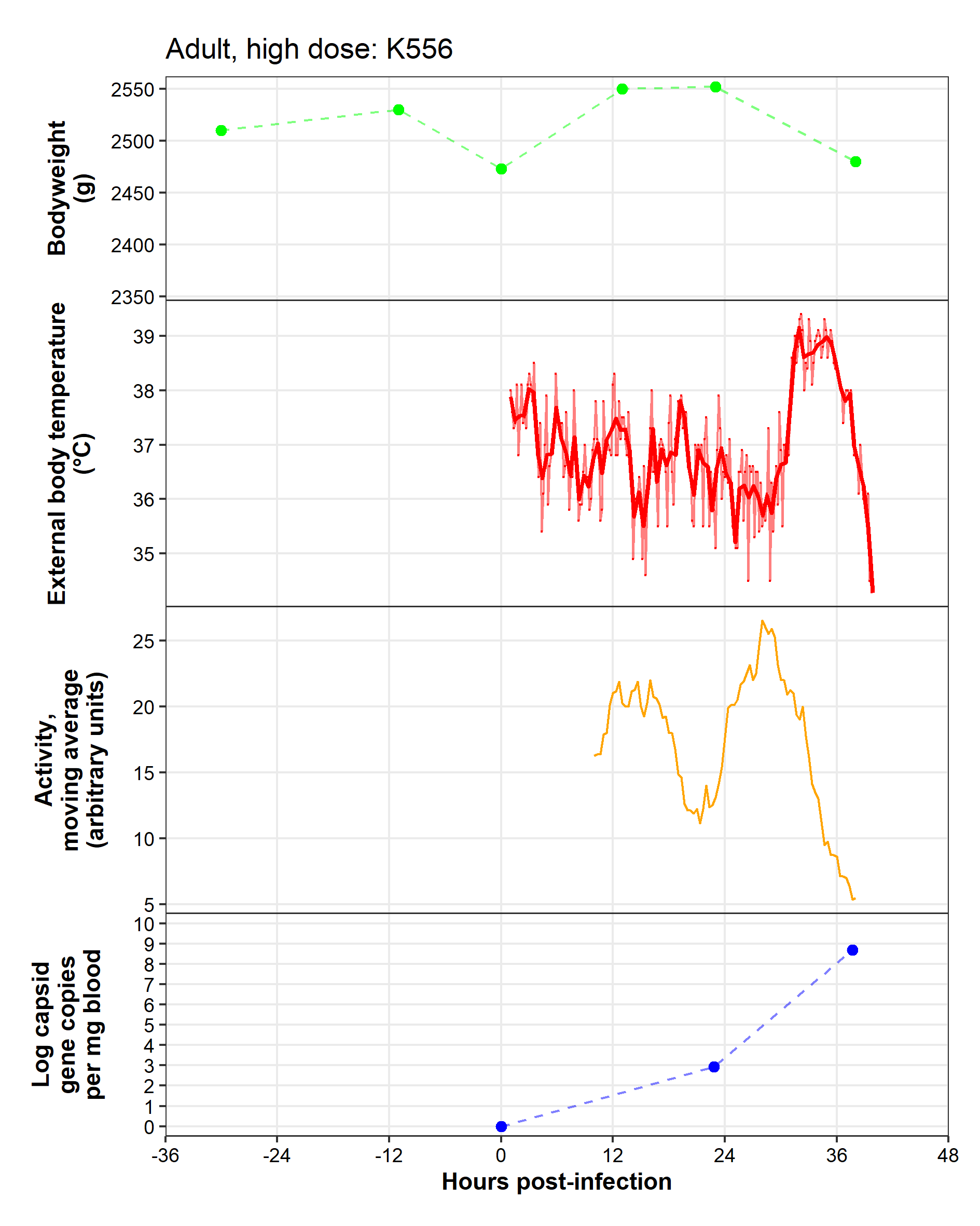

**Caveats:** Technical difficulties with activity monitor prior to 10 hpi and with temperature monitor prior to infection.

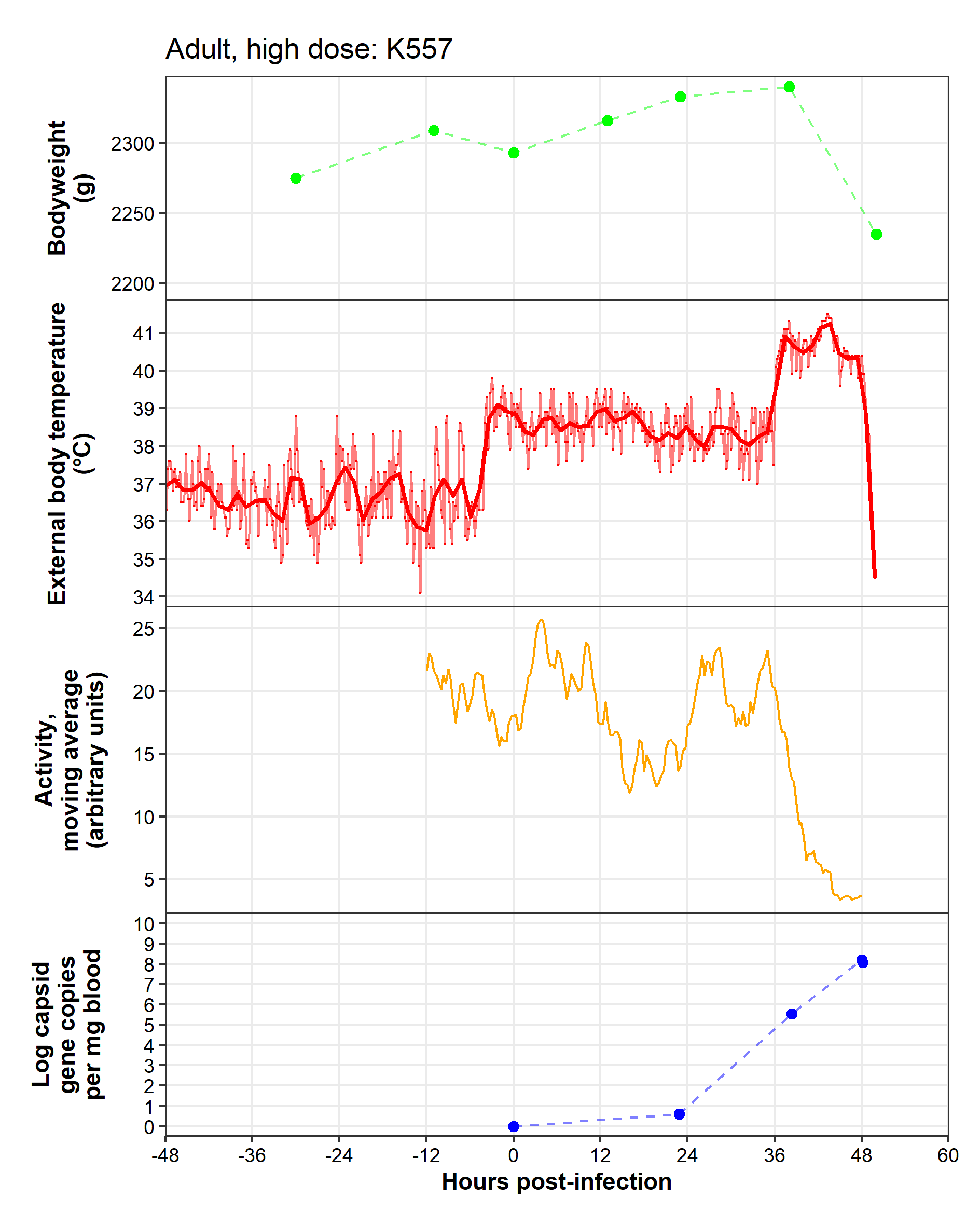

**Caveats:** Technical difficulties with activity monitor prior to -12 hpi. Collar loose prior to -6 hpi.

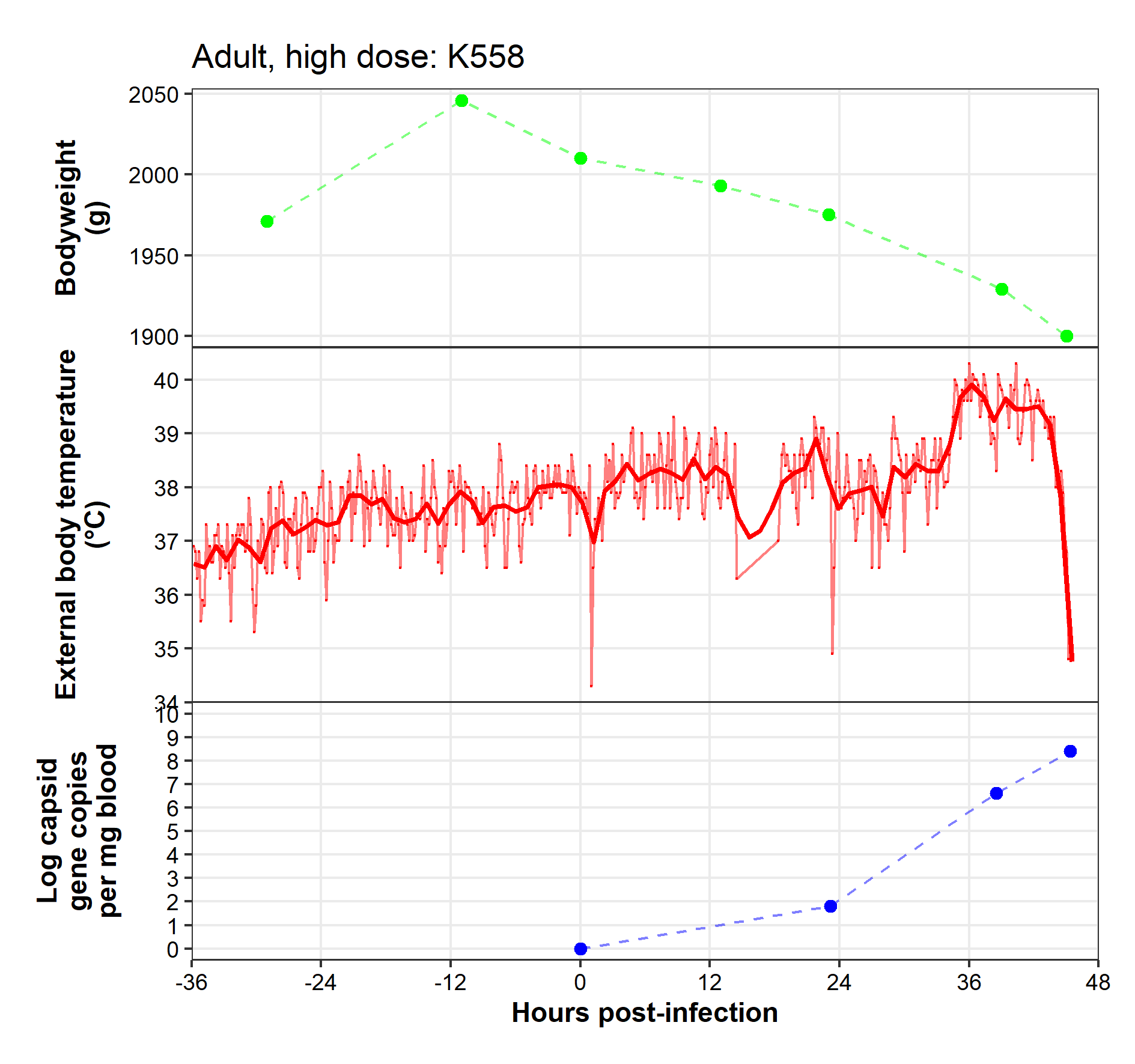

**Caveats:** Technical difficulties with activity monitor. Collar removed between 13–17 hpi.

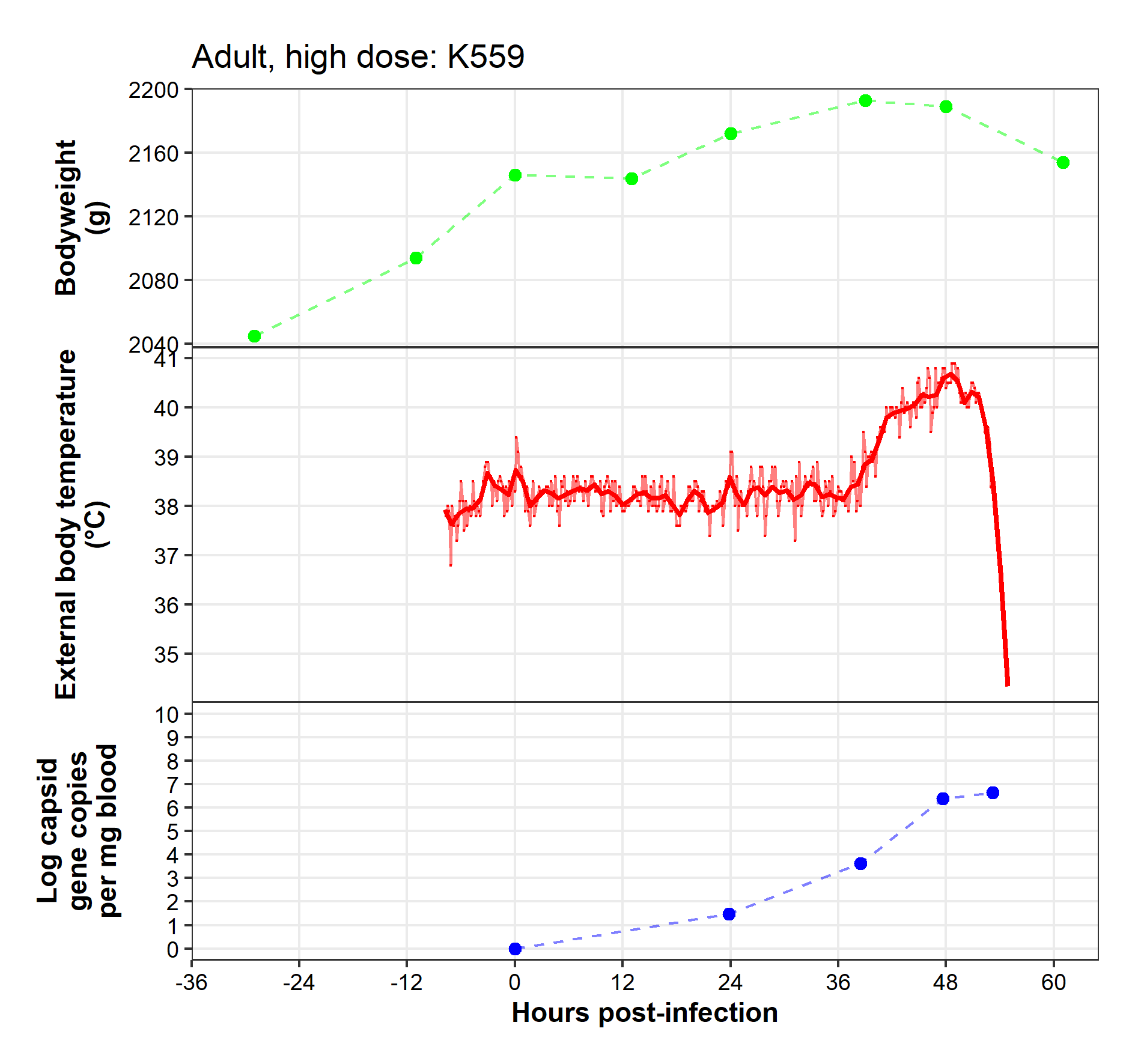

**Caveats:** Technical difficulties with activity monitor. Collar loose prior to -8 hpi.

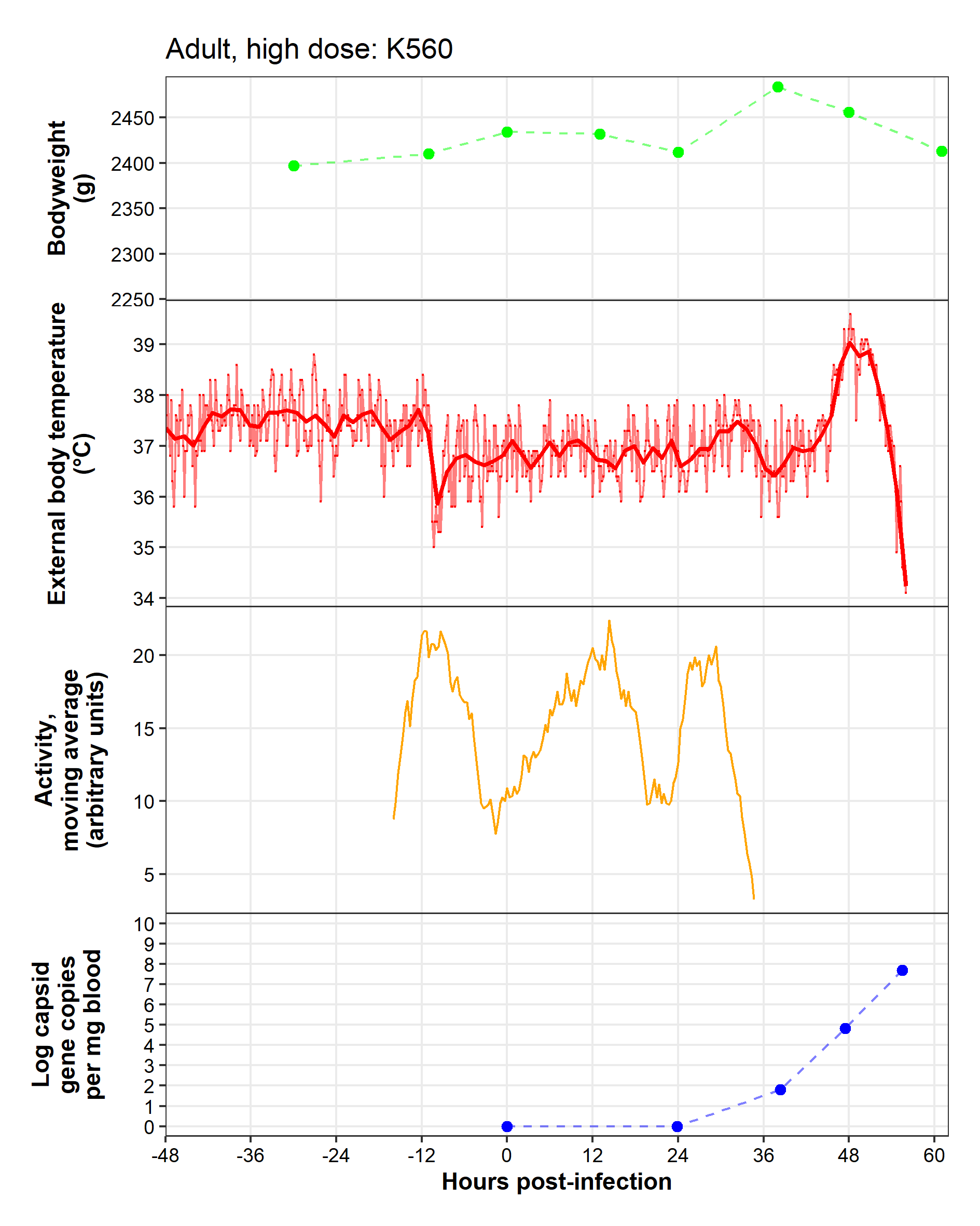

**Caveats:** Technical difficulties with activity monitor prior to -16 hpi and after 35 hpi.

Adult, low dose

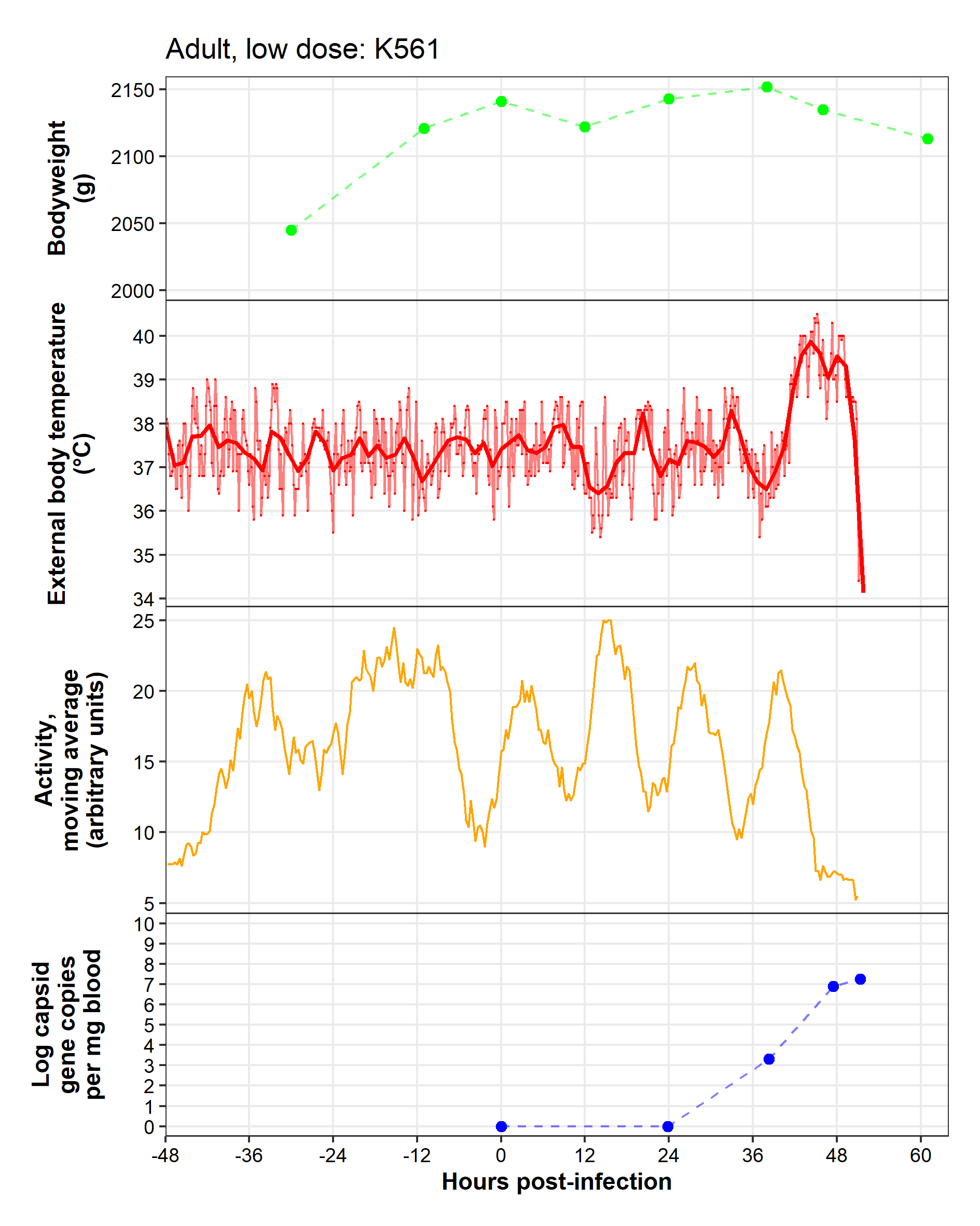

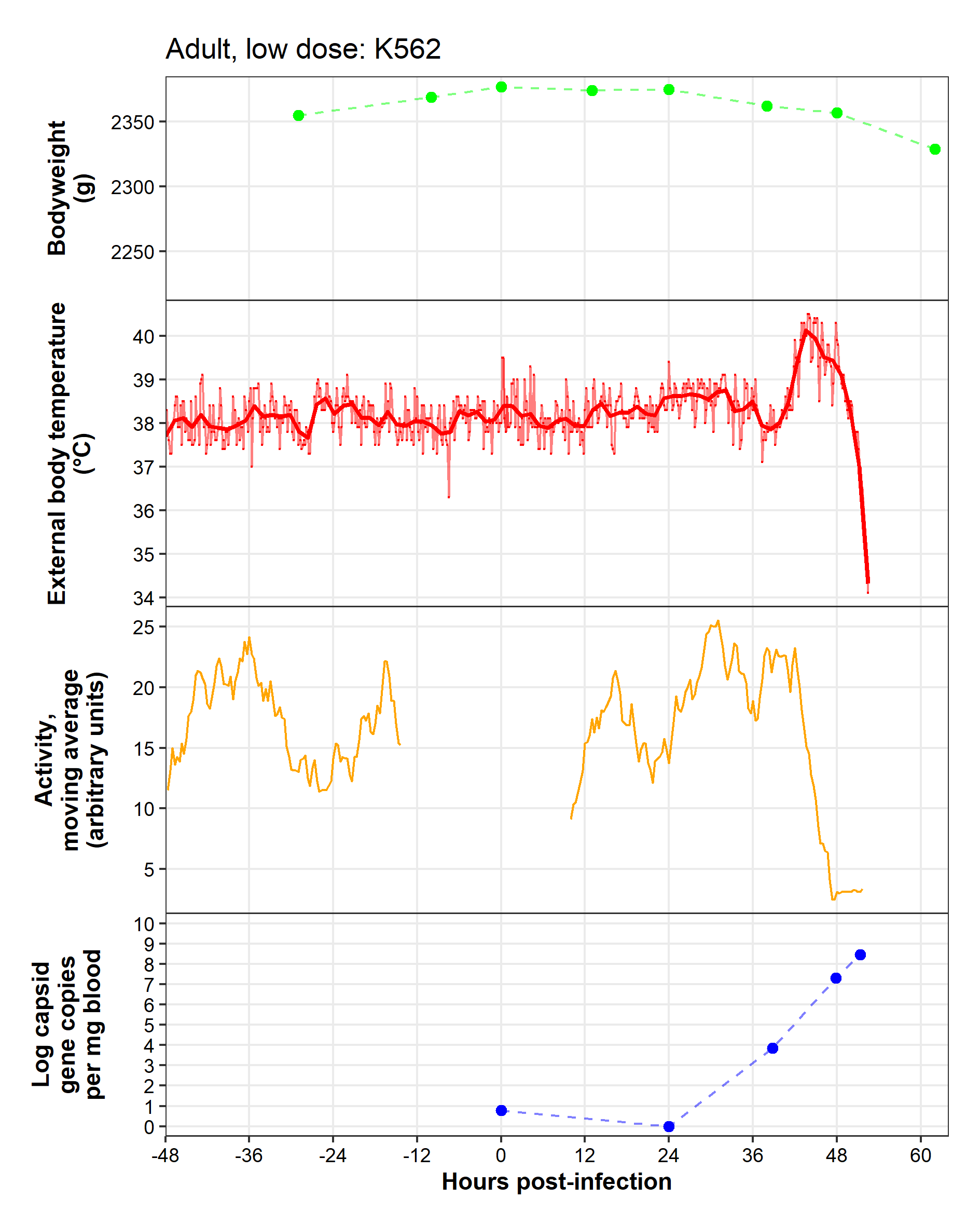

**Caveats:** Technical difficulties with activity monitor from -14–10 hpi.

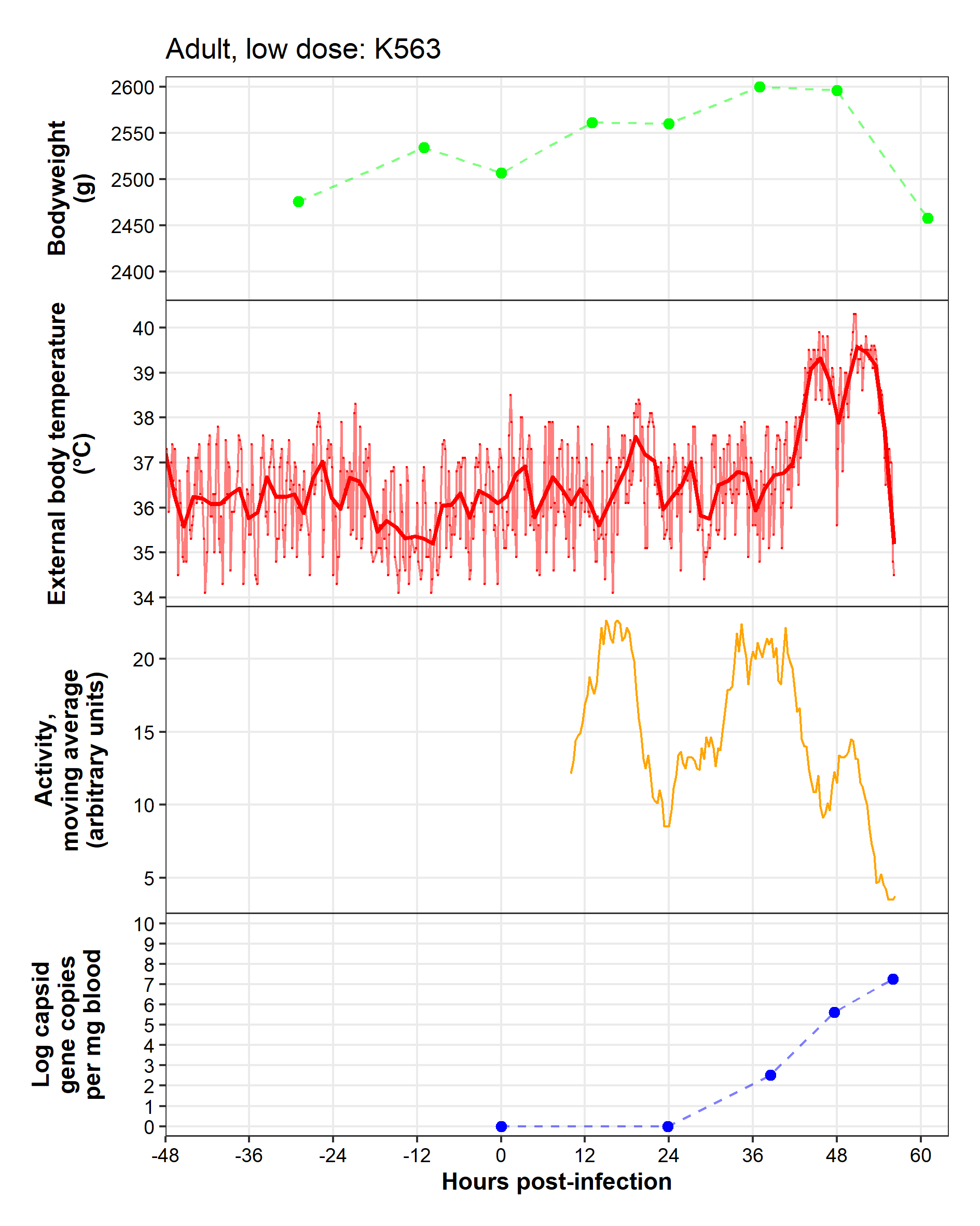

**Caveats:** Technical difficulties with activity monitor prior to -10 hpi.

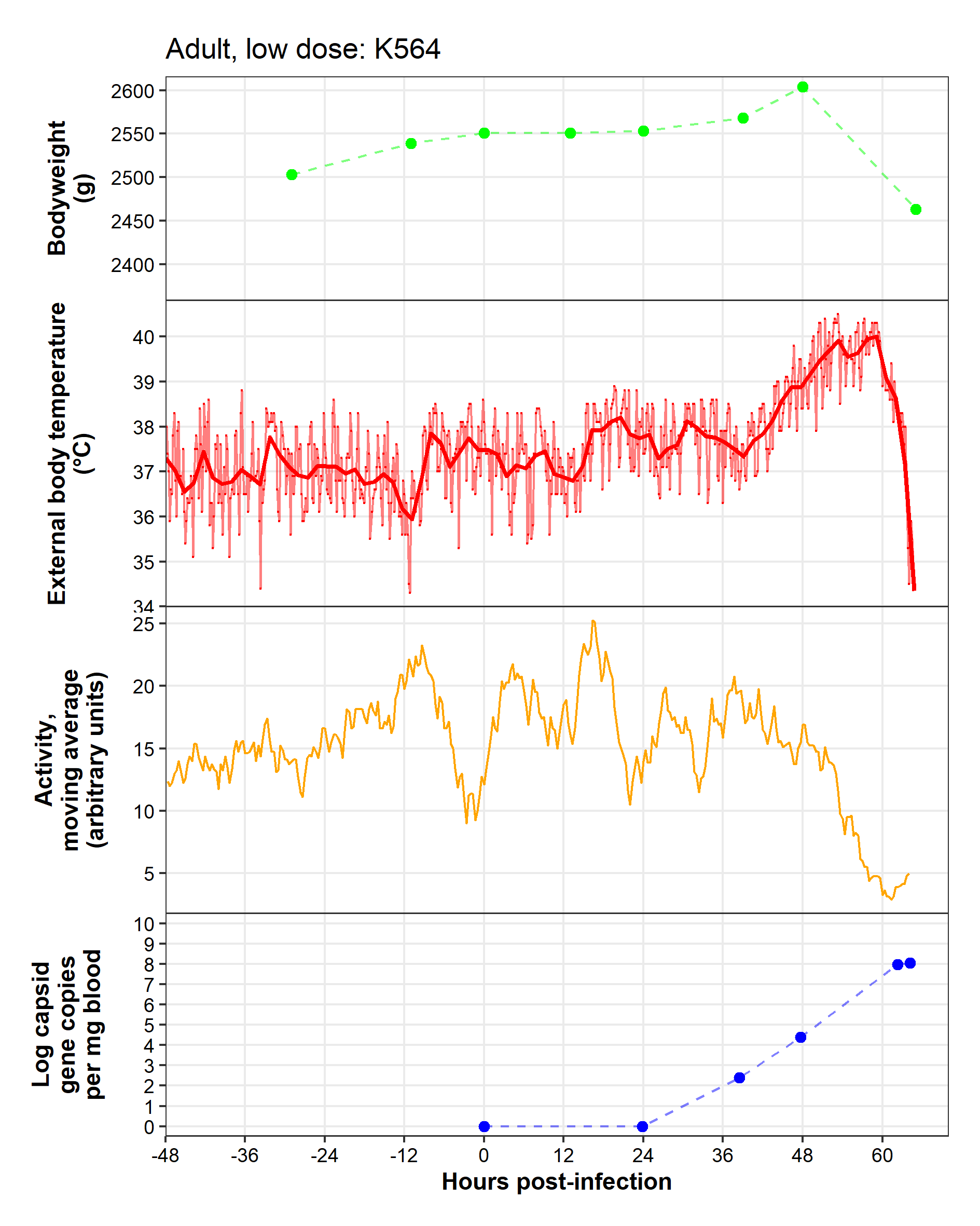

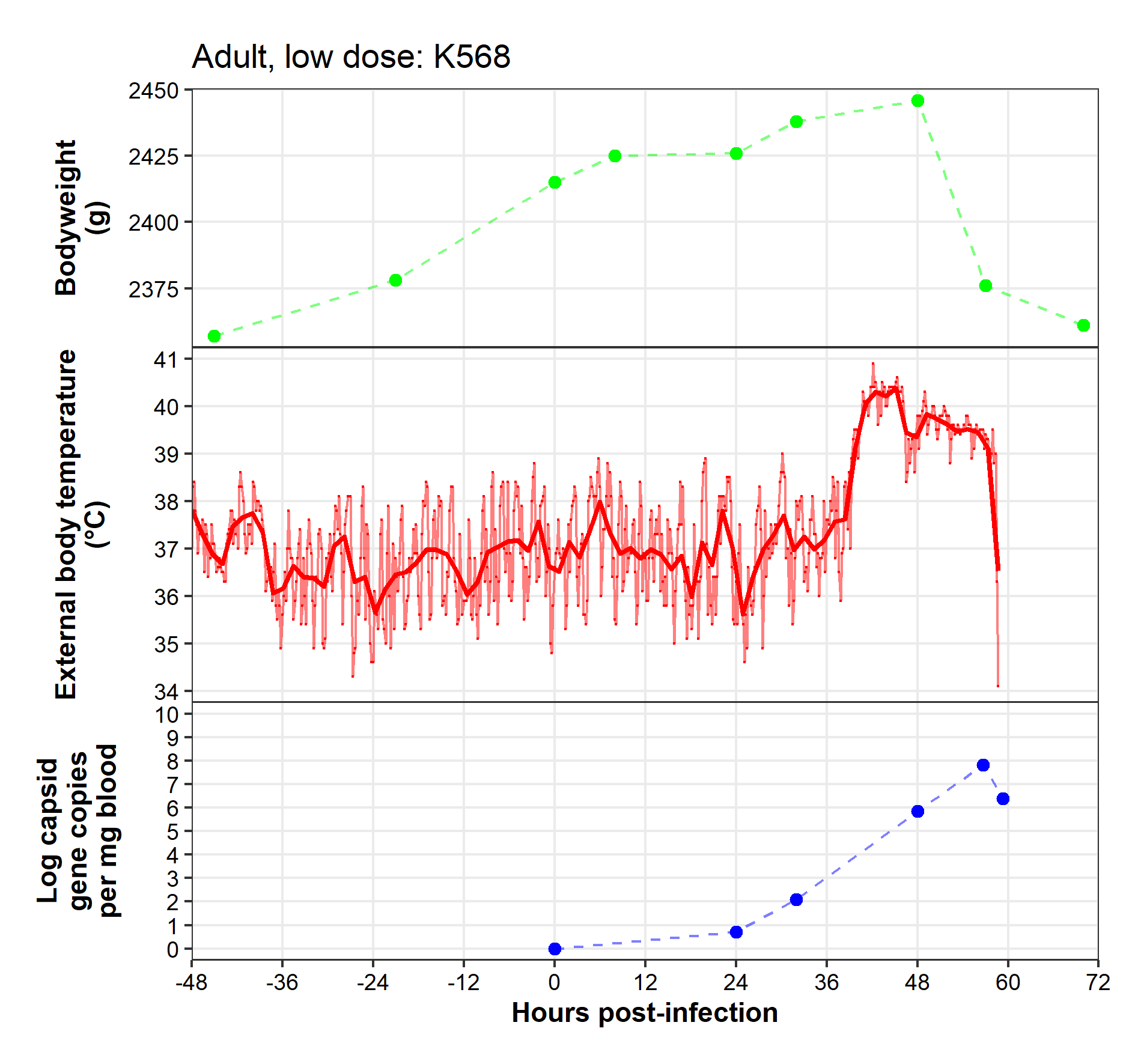

**Caveats:** Activity monitors were not used in trial 4 due to technical issues with the app update.

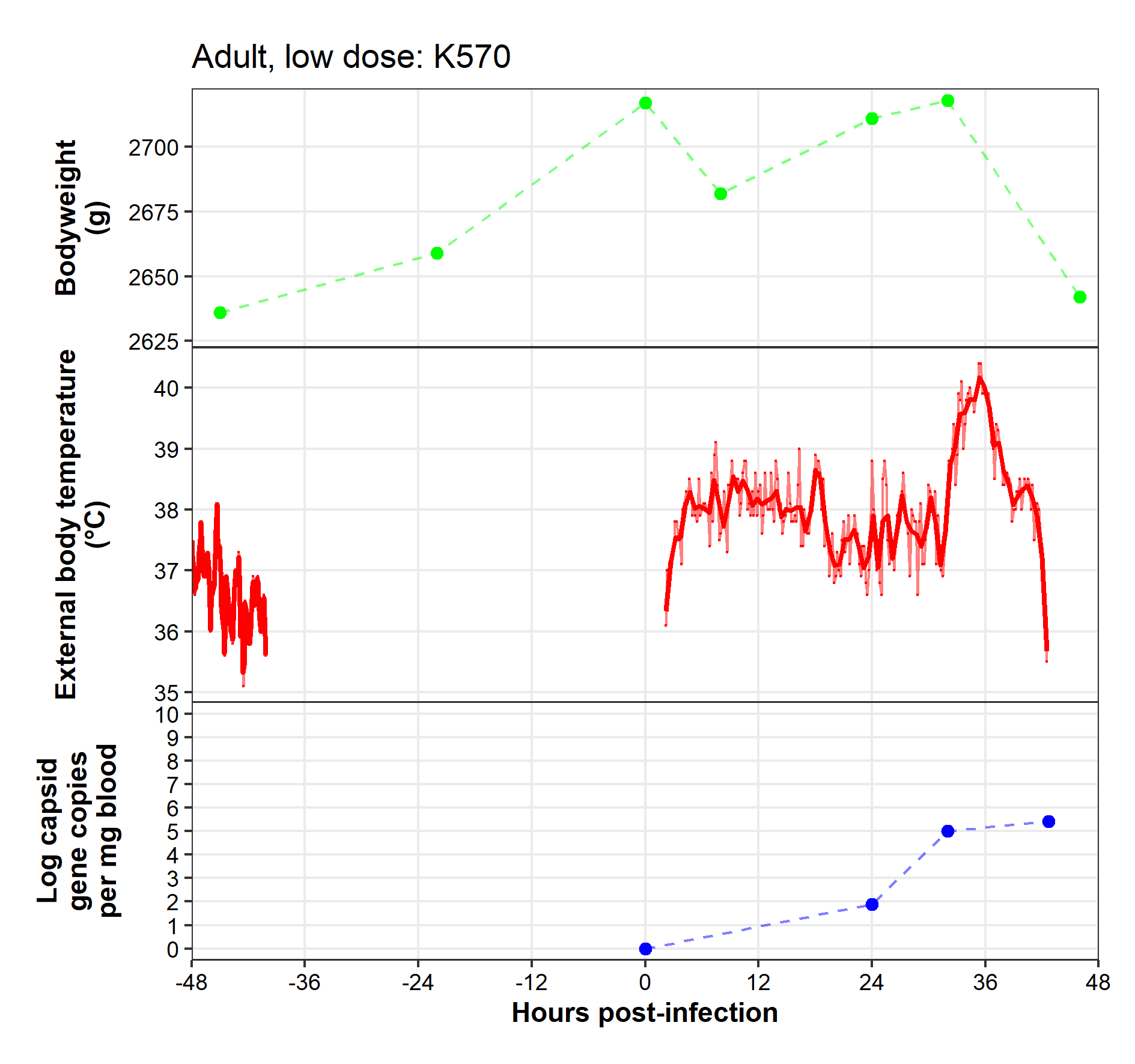

**Caveats:** Activity monitors were not used in trial 4 due to technical issues with the app update. Temperature monitor fell out of pouch around -40 hpi, replaced at 2 hpi.

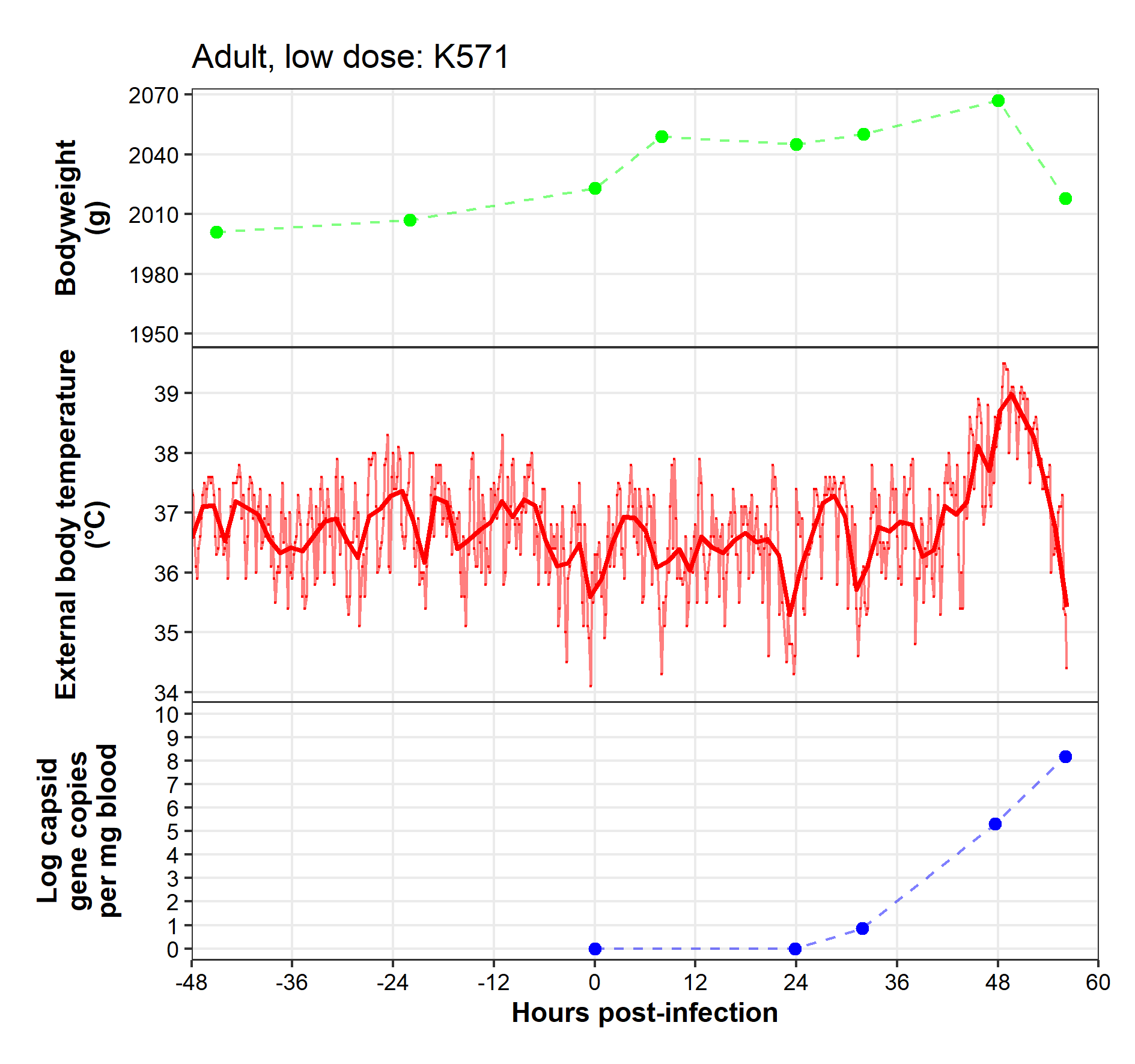

**Caveats:** Activity monitors were not used in trial 4 due to technical issues with the app update.

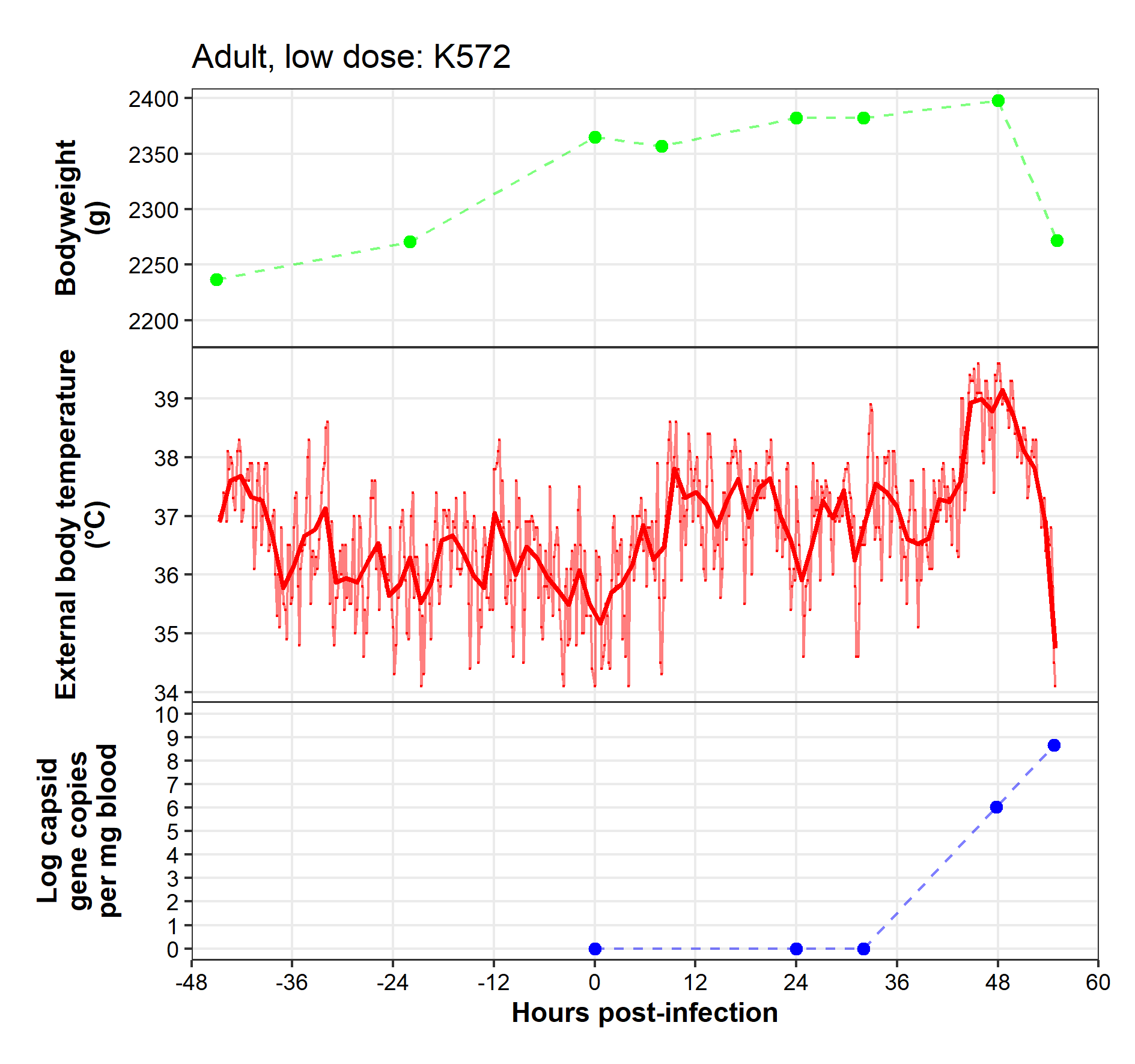

**Caveats:** Activity monitors were not used in trial 4 due to technical issues with the app update.

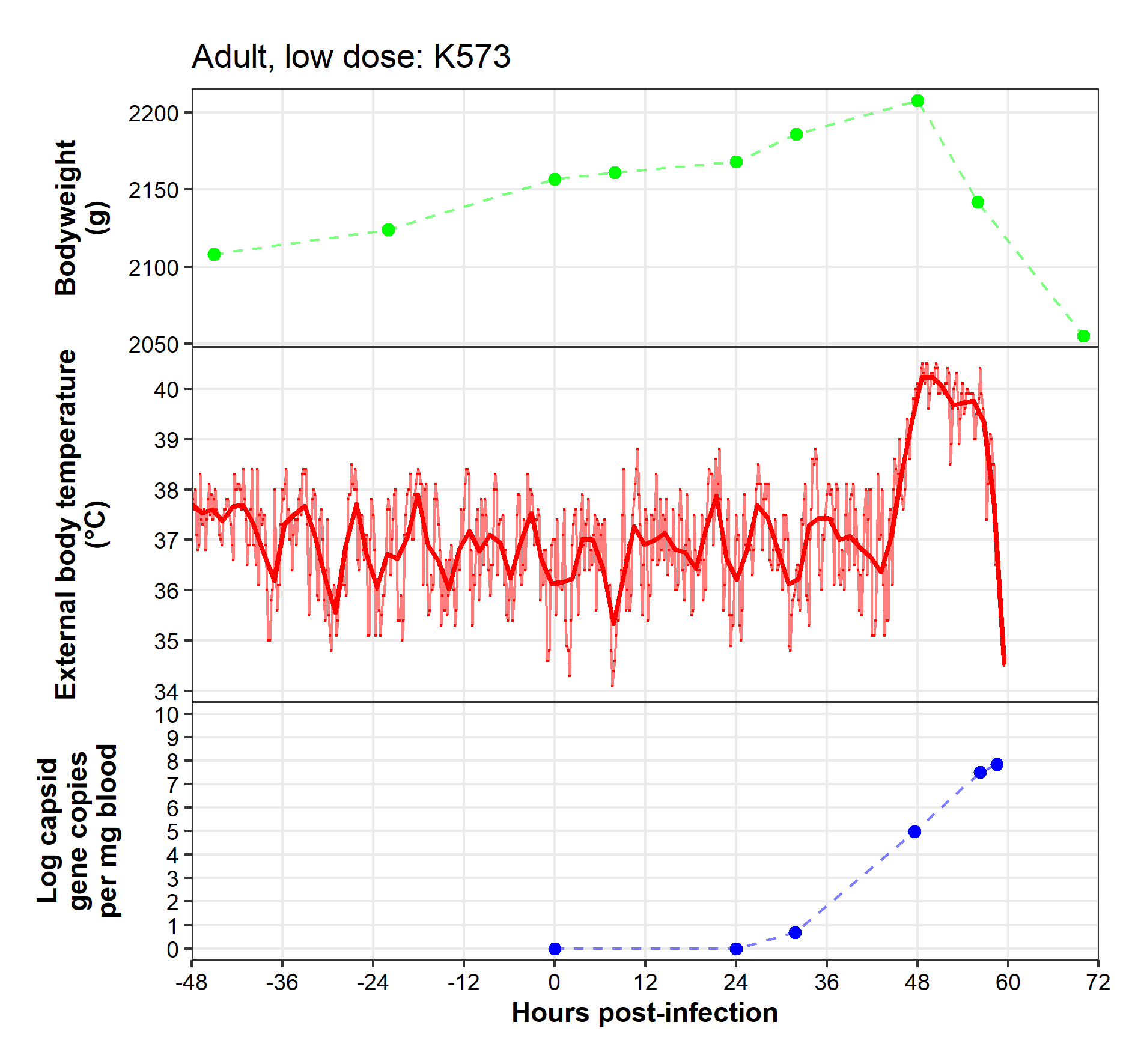

**Caveats:** Activity monitors were not used in trial 4 due to technical issues with the app update.

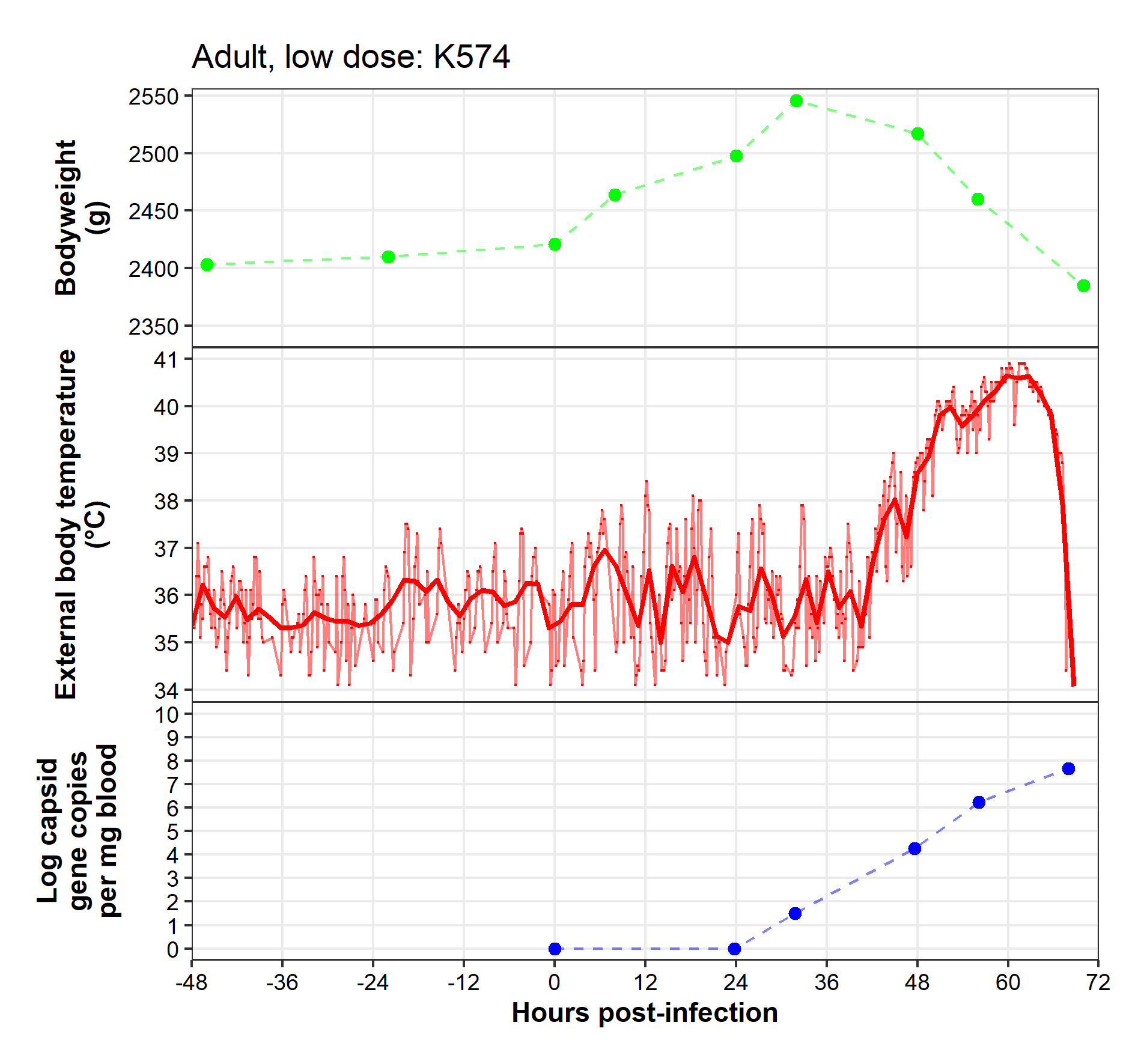

**Caveats:** Activity monitors were not used in trial 4 due to technical issues with the app update.

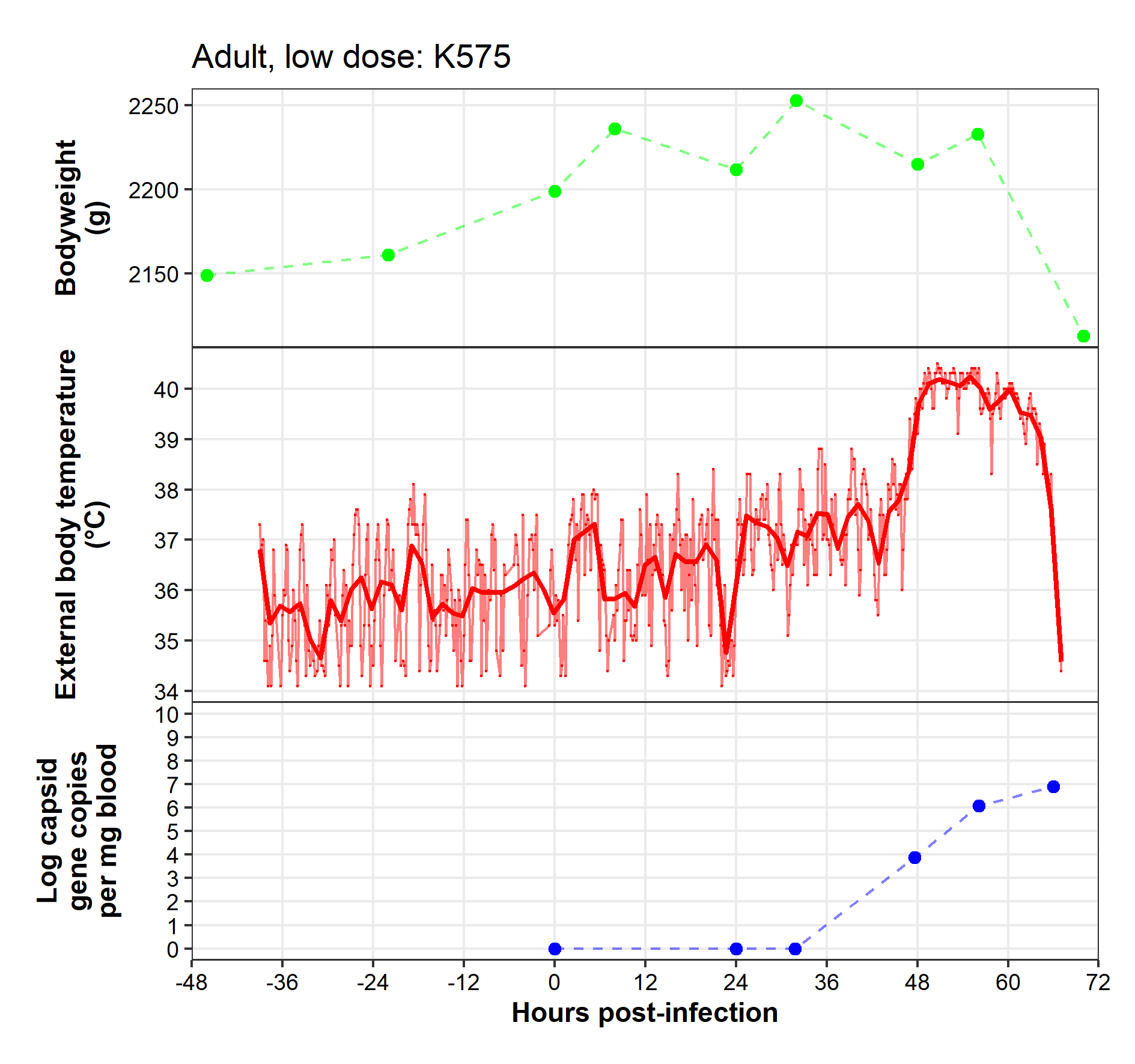

**Caveats:** Activity monitors were not used in trial 4 due to technical issues with the app update. Technical difficulties with collar prior to -38 hpi.

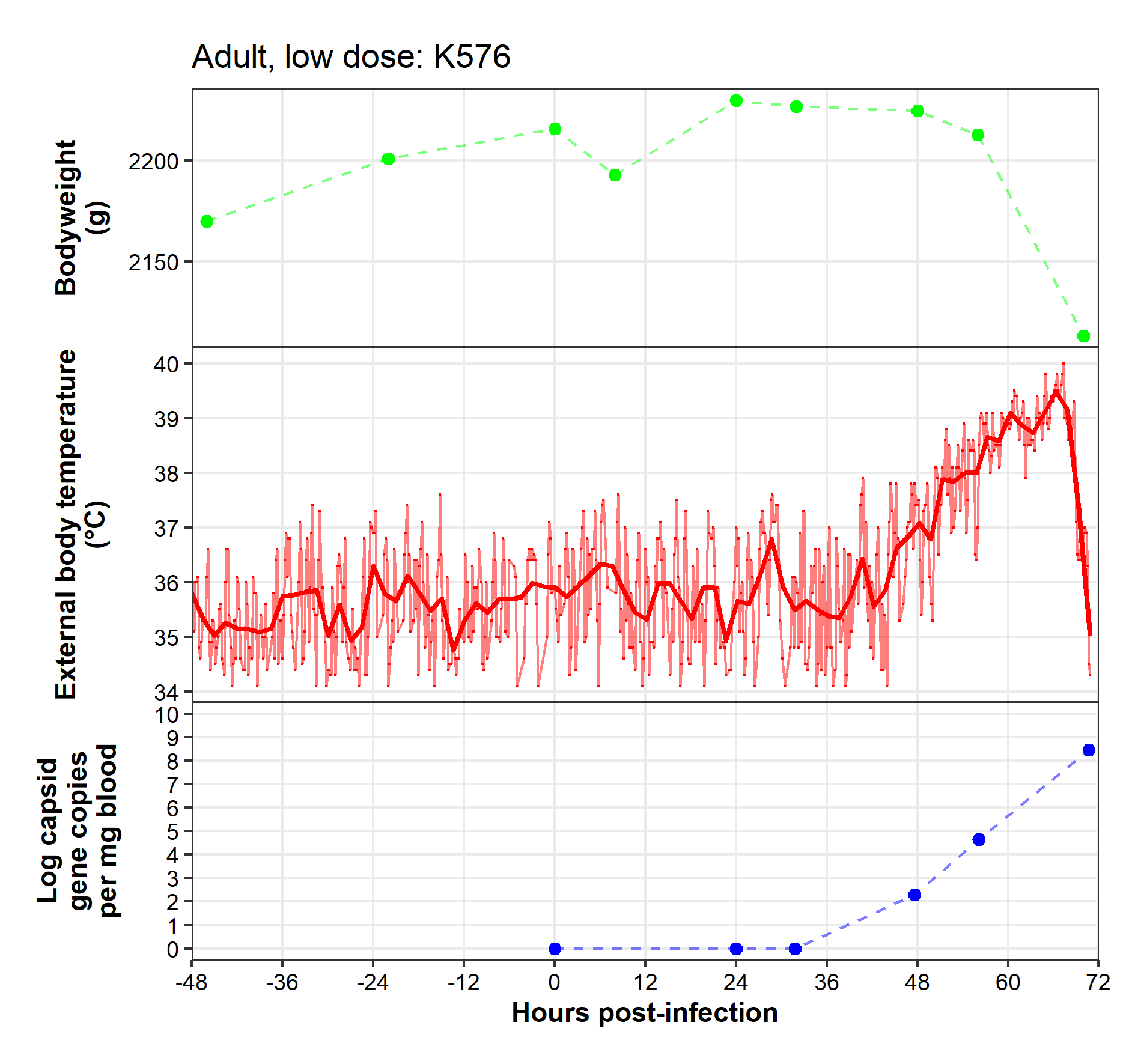

**Caveats:** Activity monitors were not used in trial 4 due to technical issues with the app update.

Kitten, control

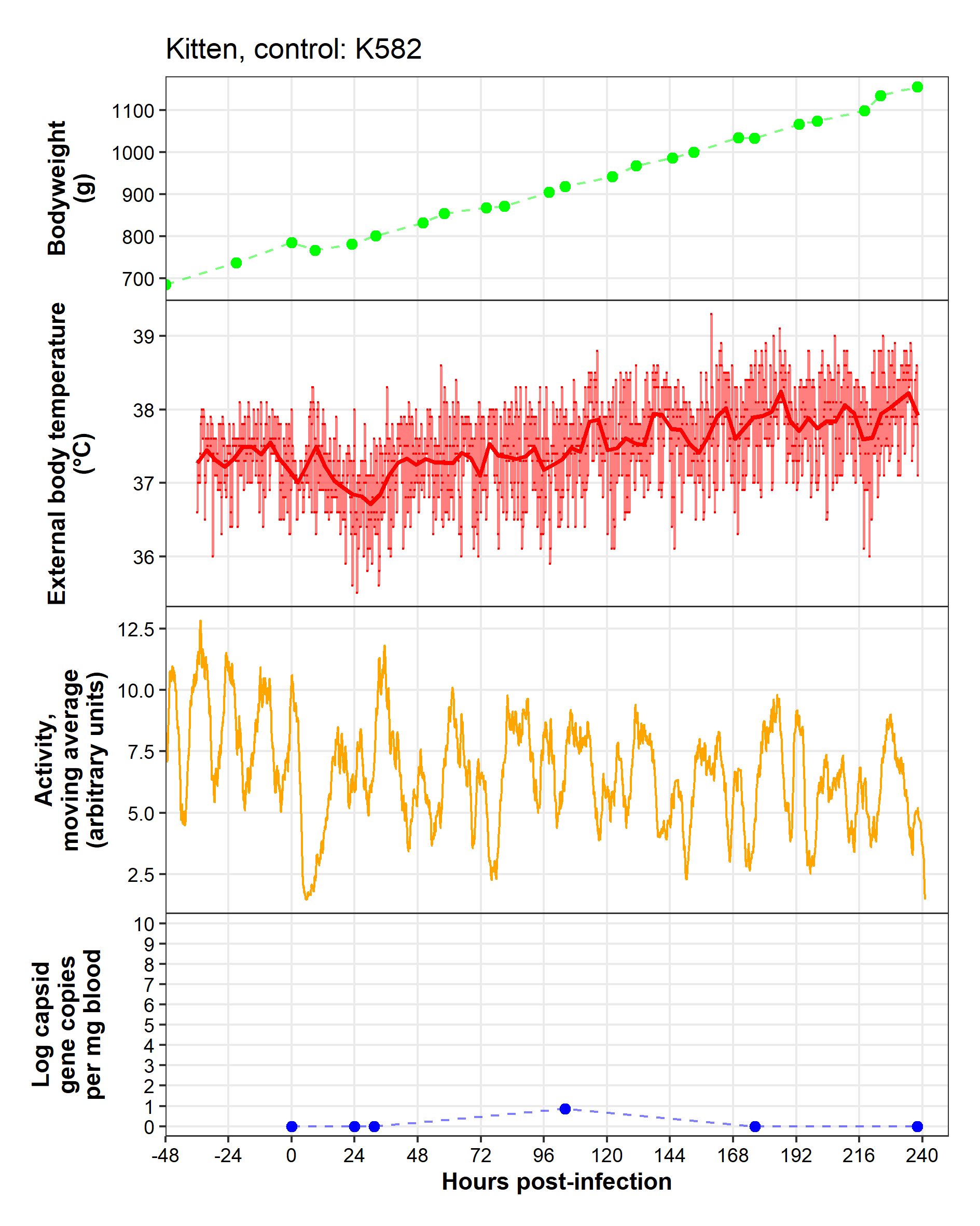

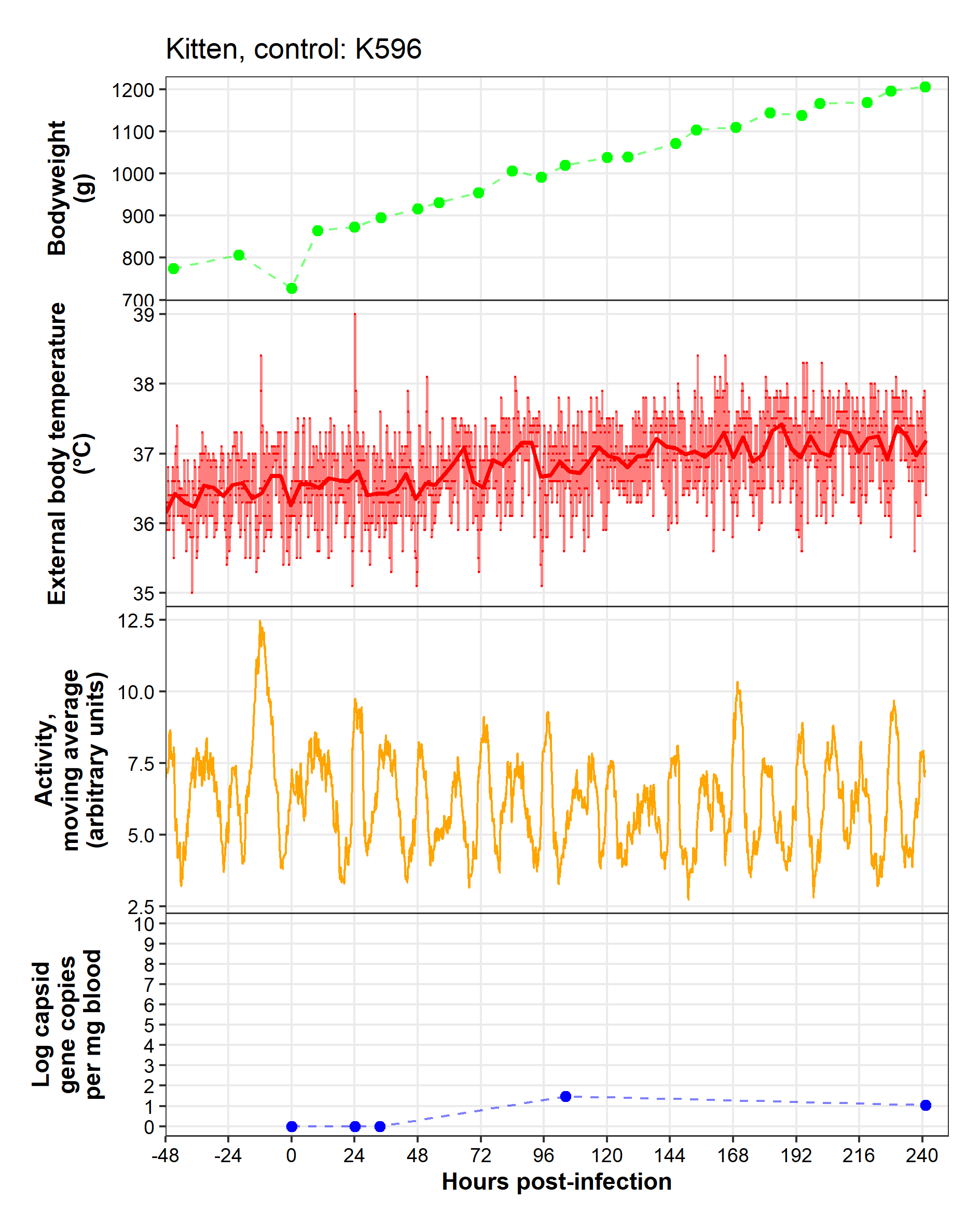

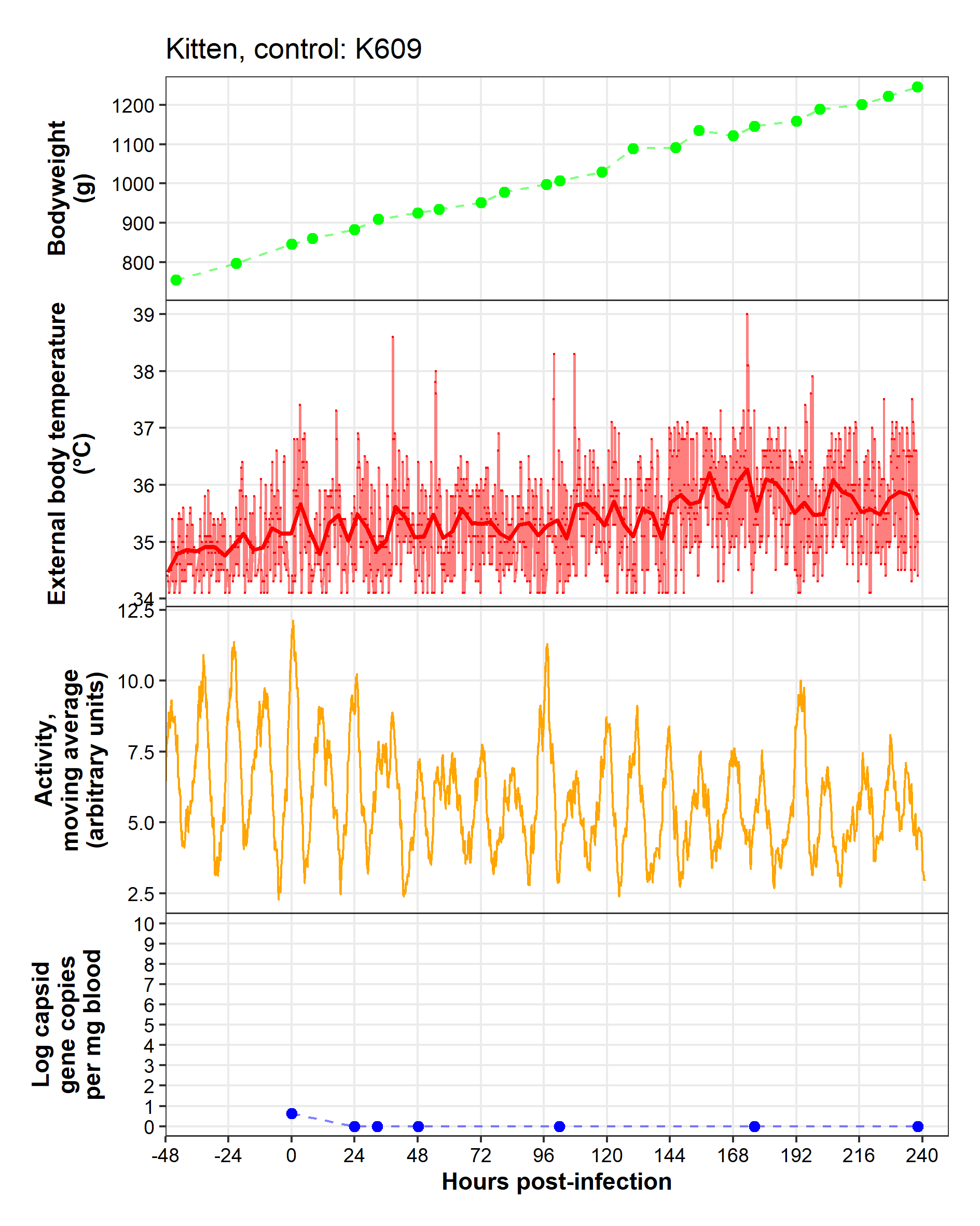

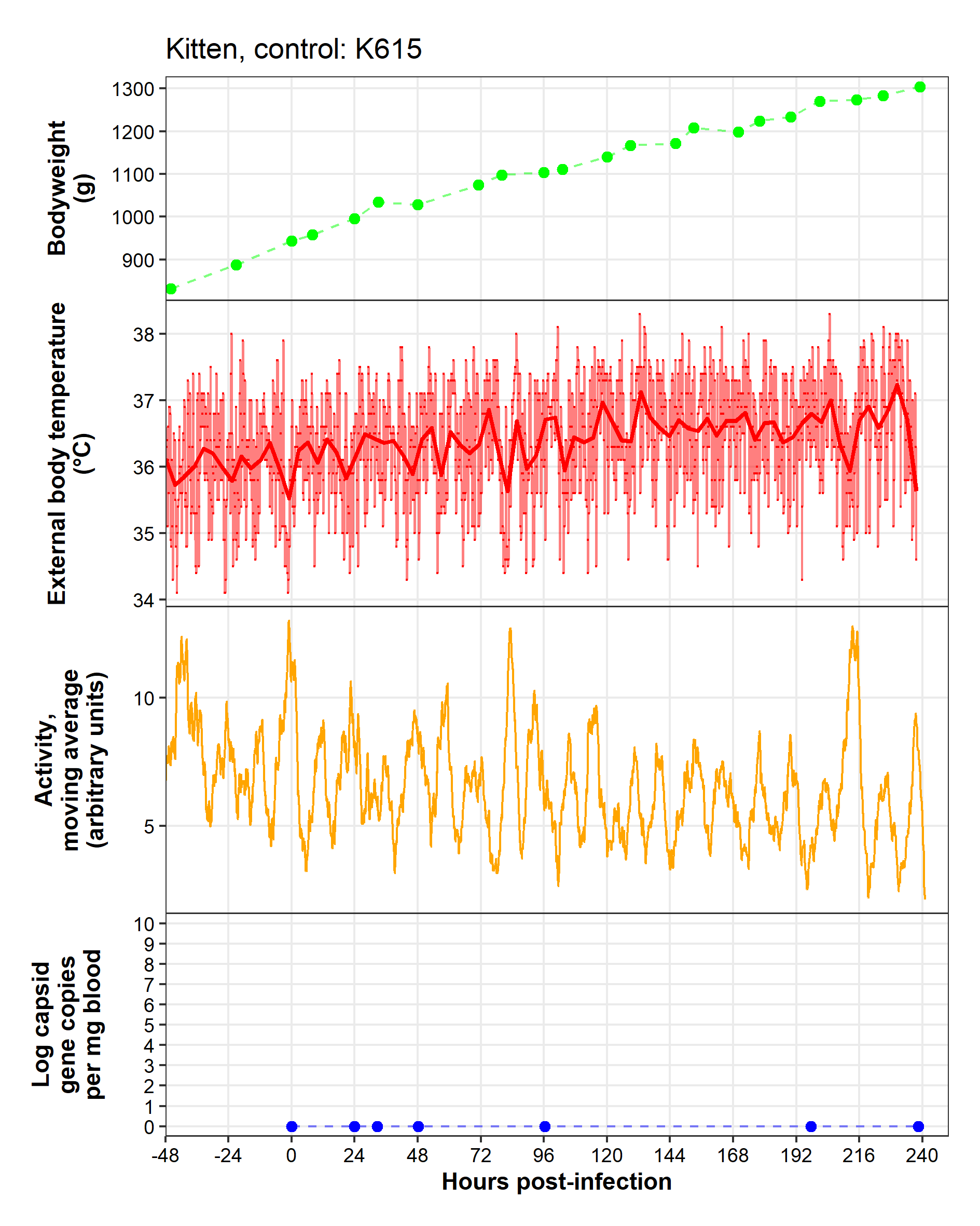

Kitten, high dose

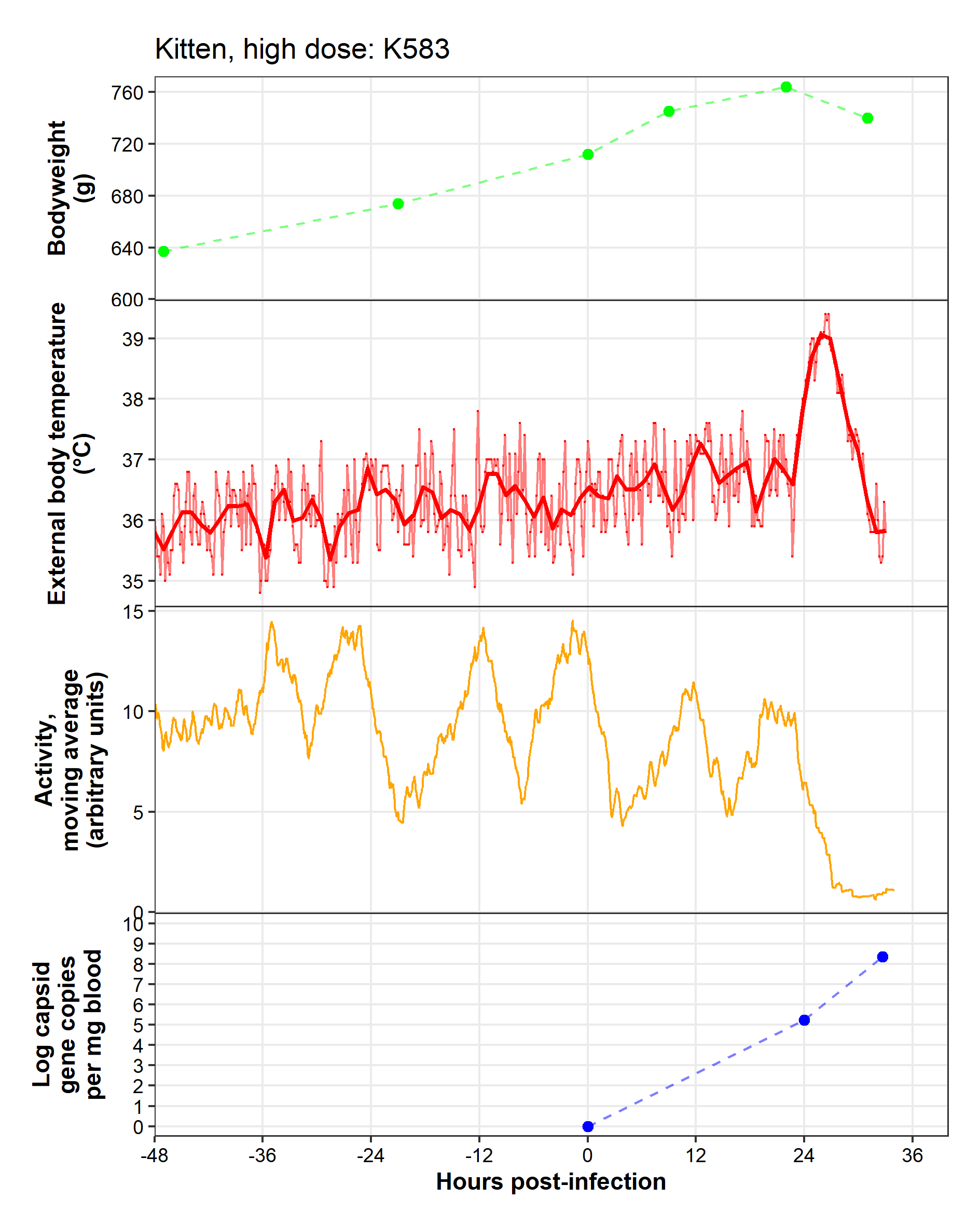

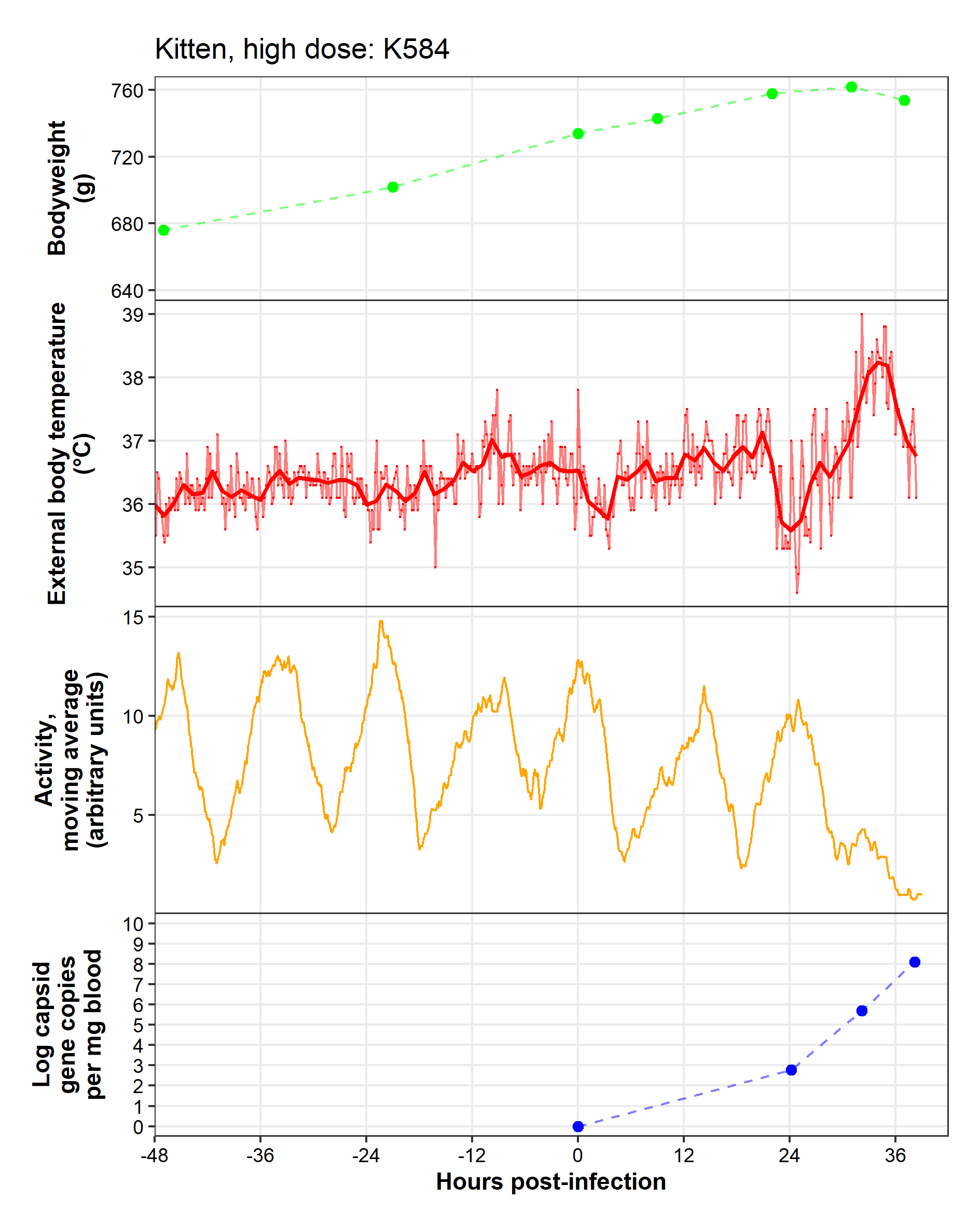

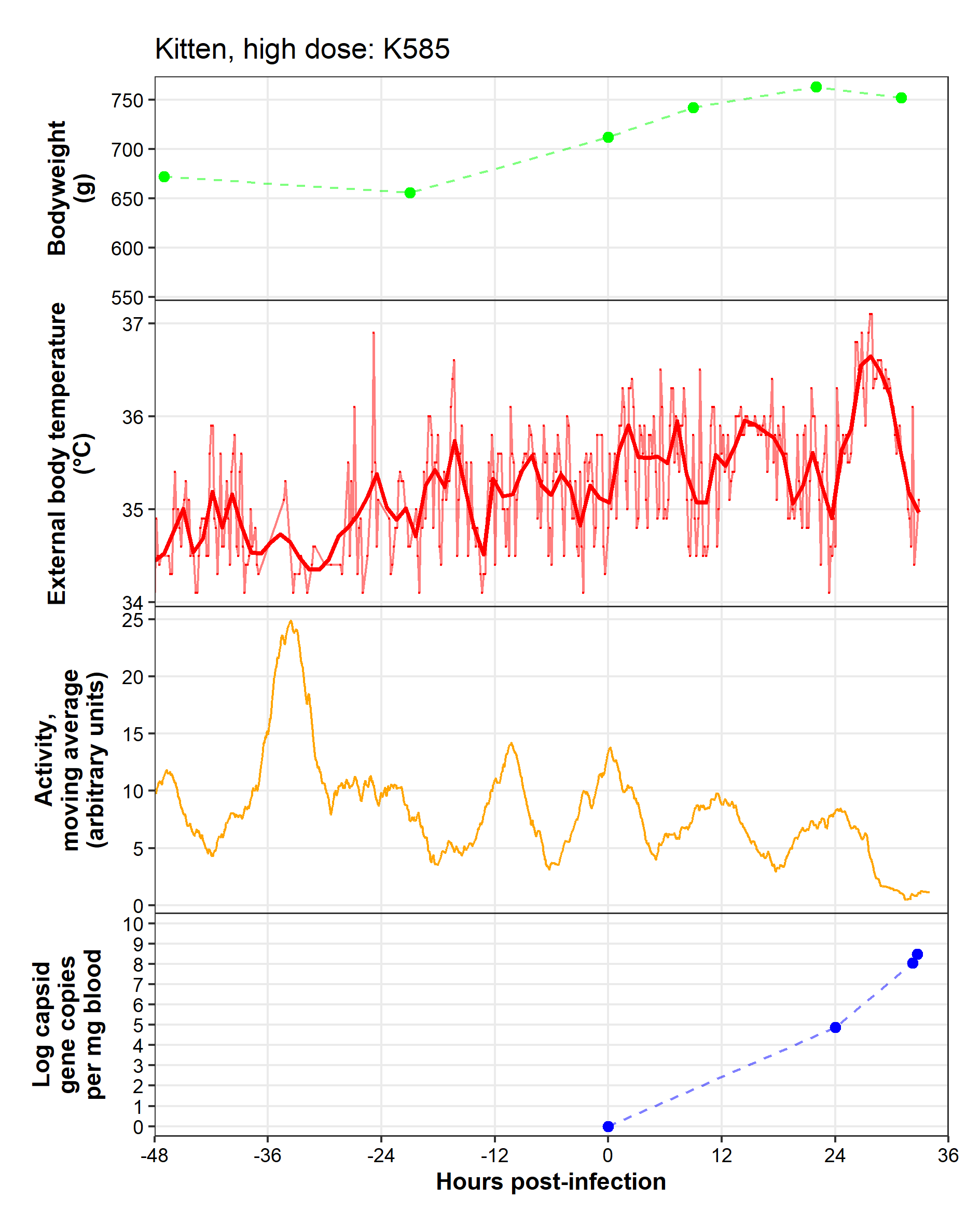

**Caveats**: Glitch in collar between -36 and -20 hpi.

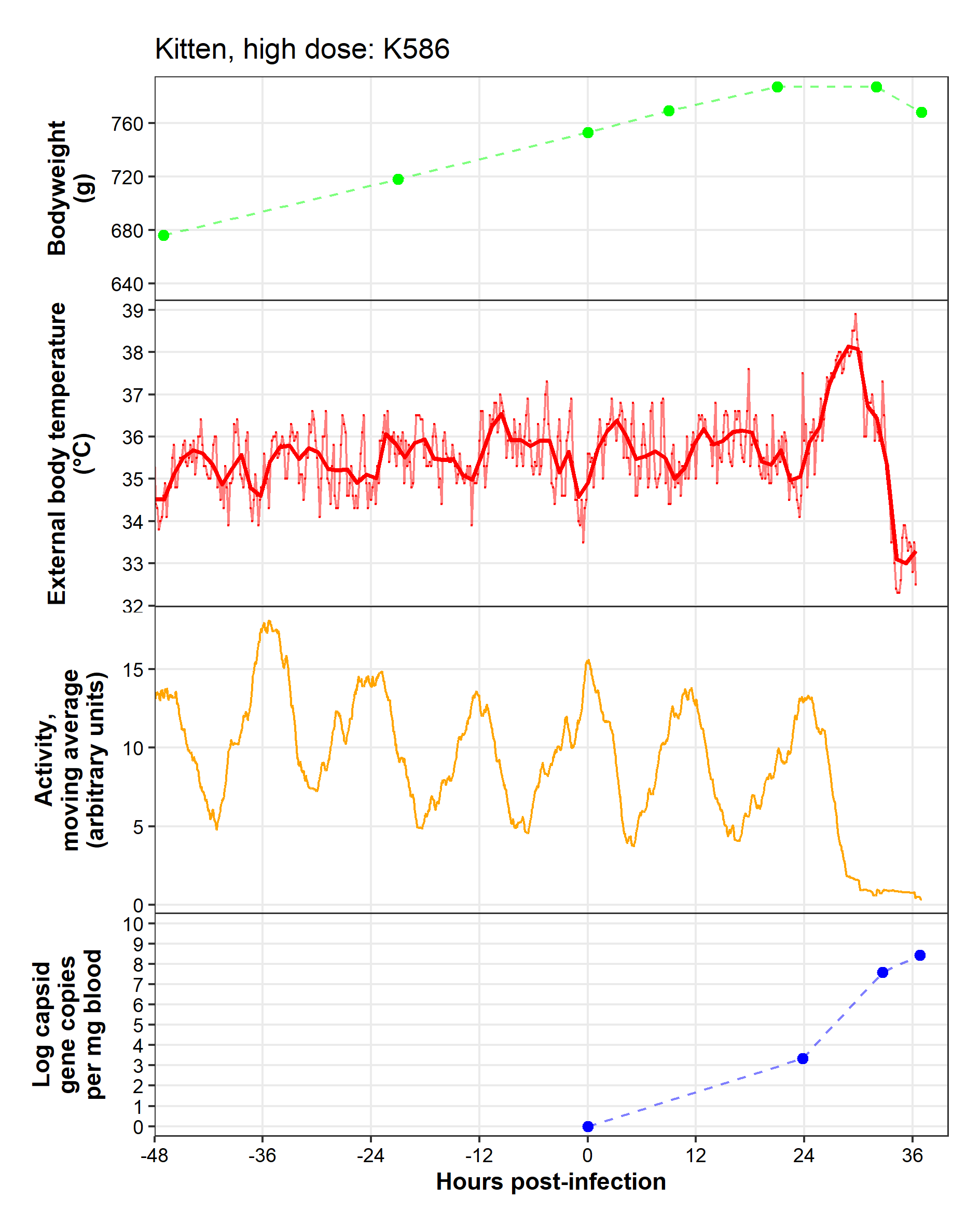

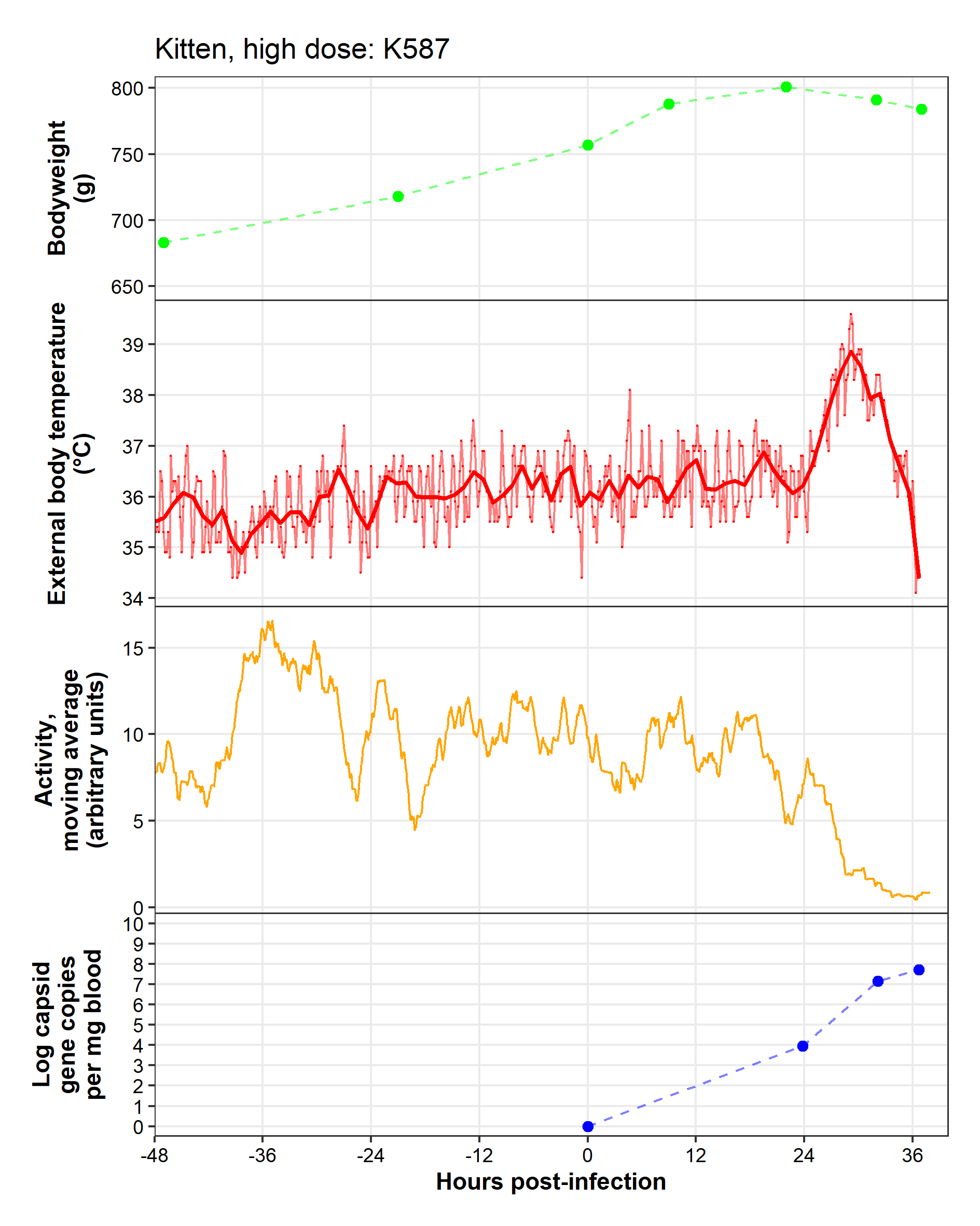

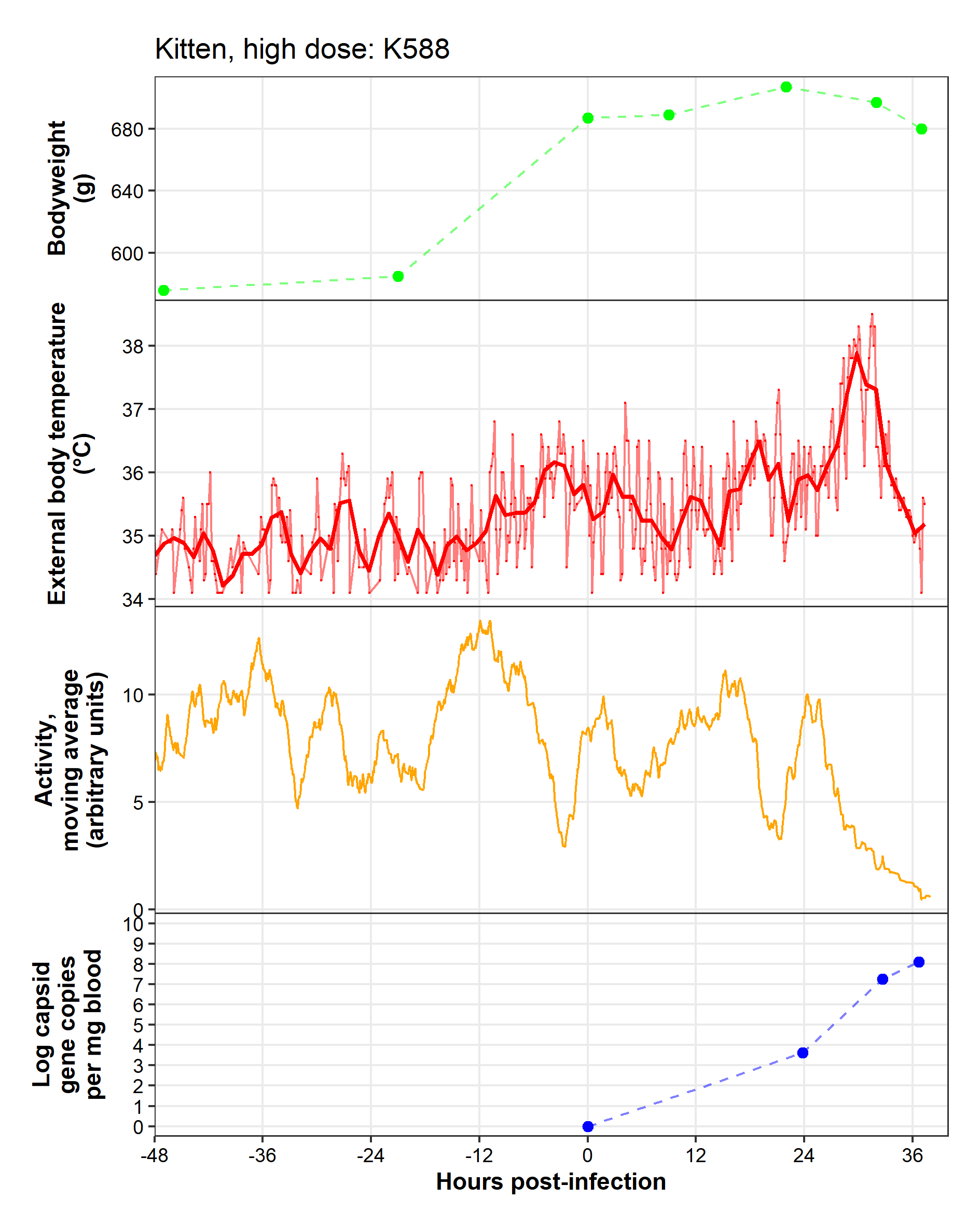

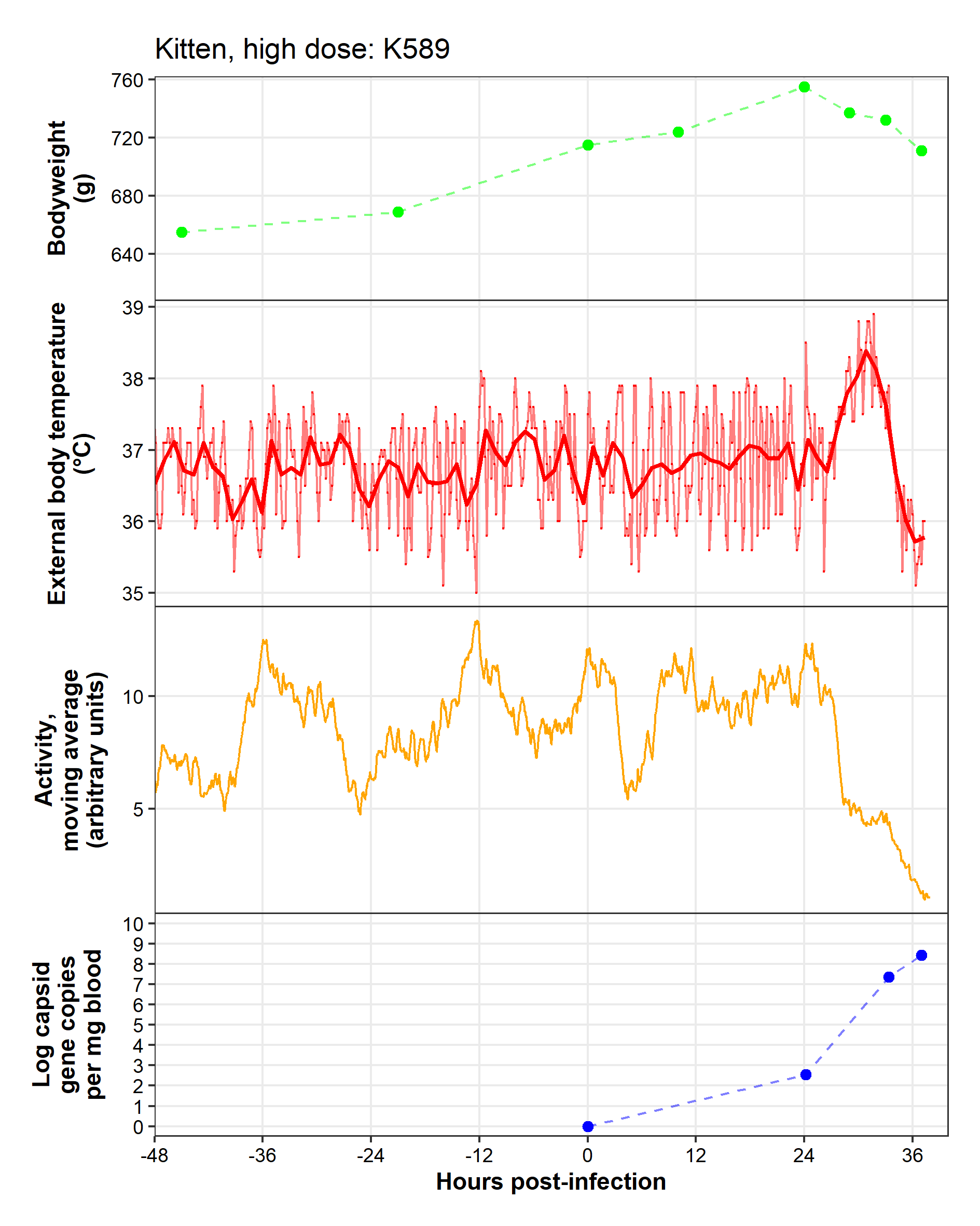

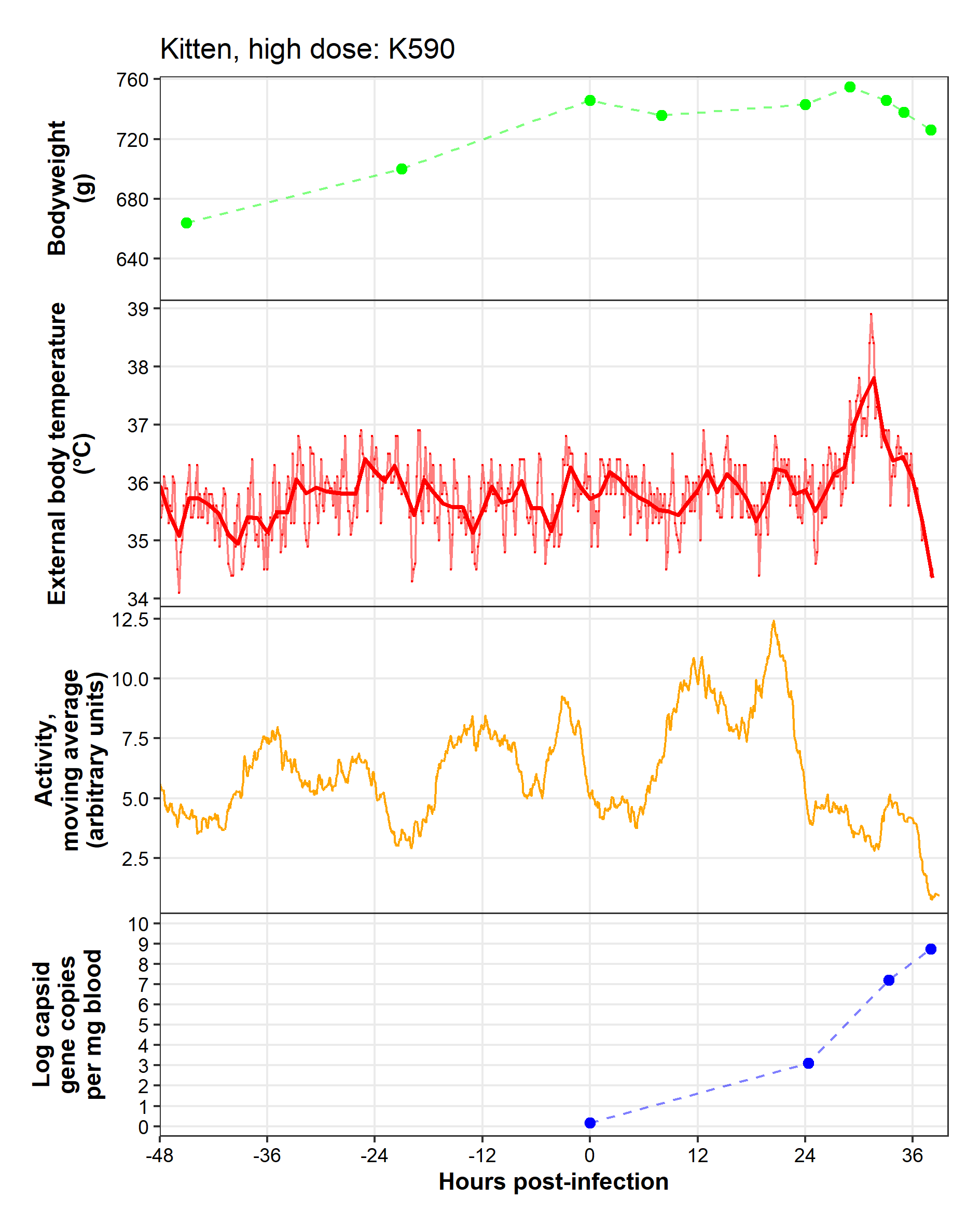

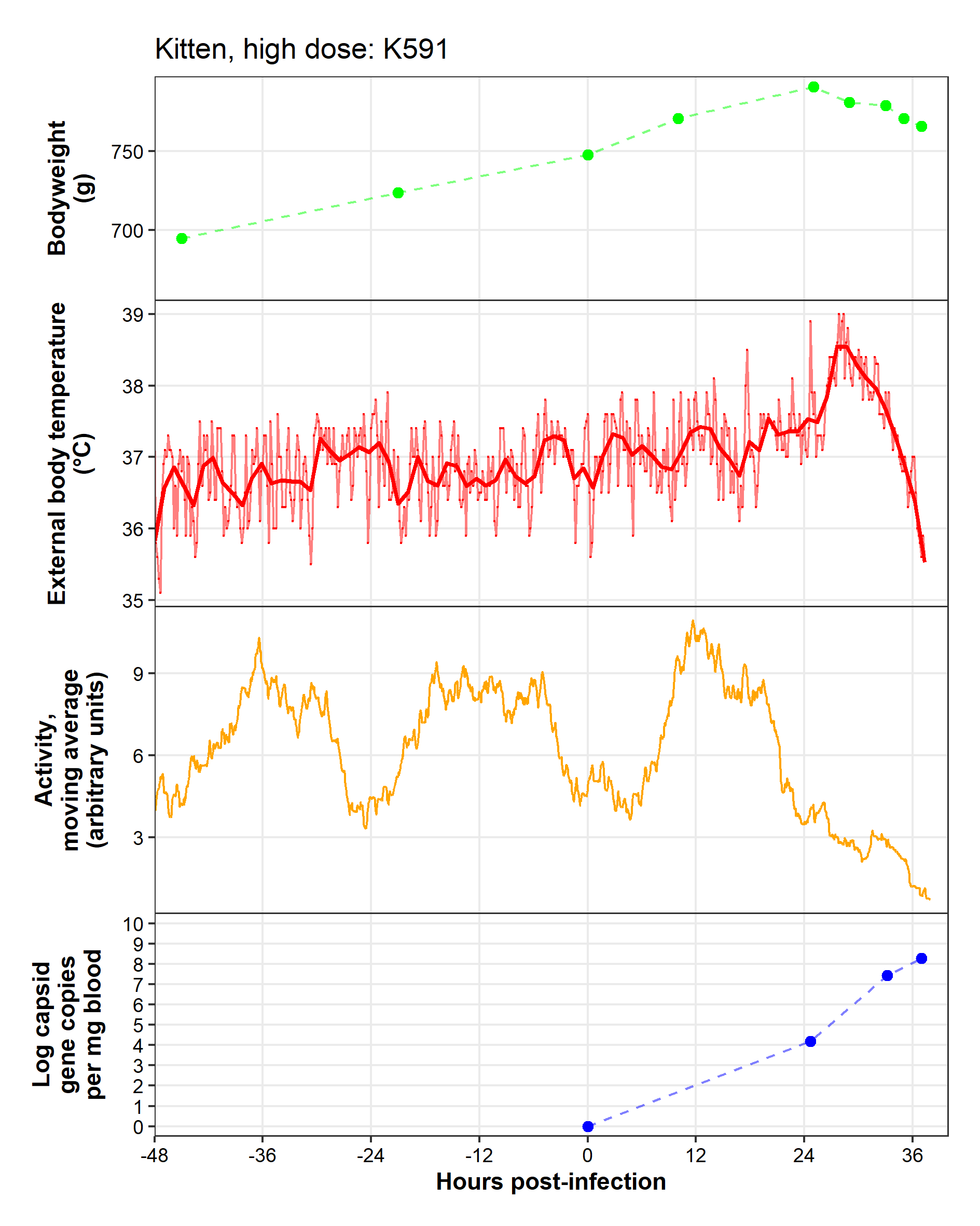

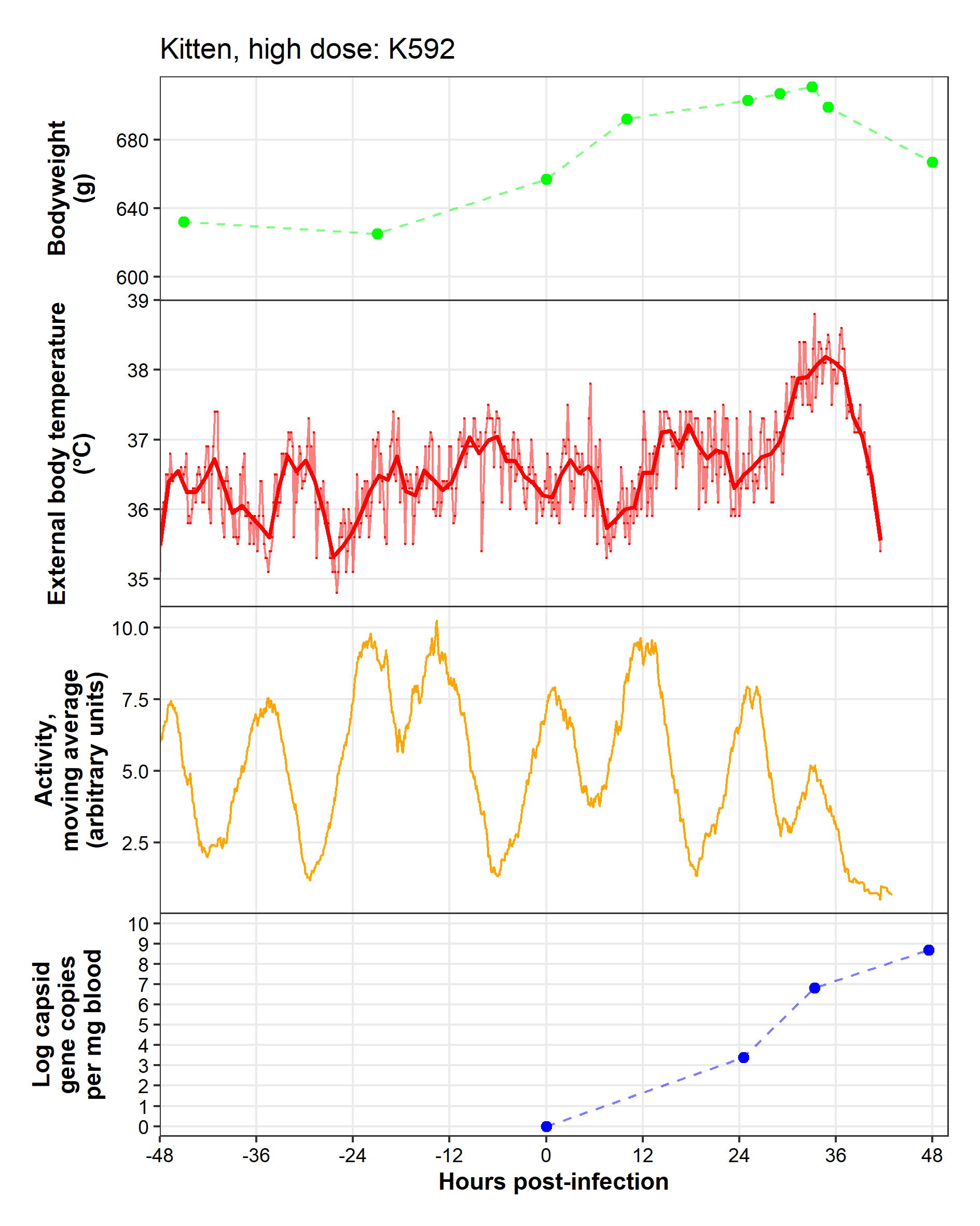

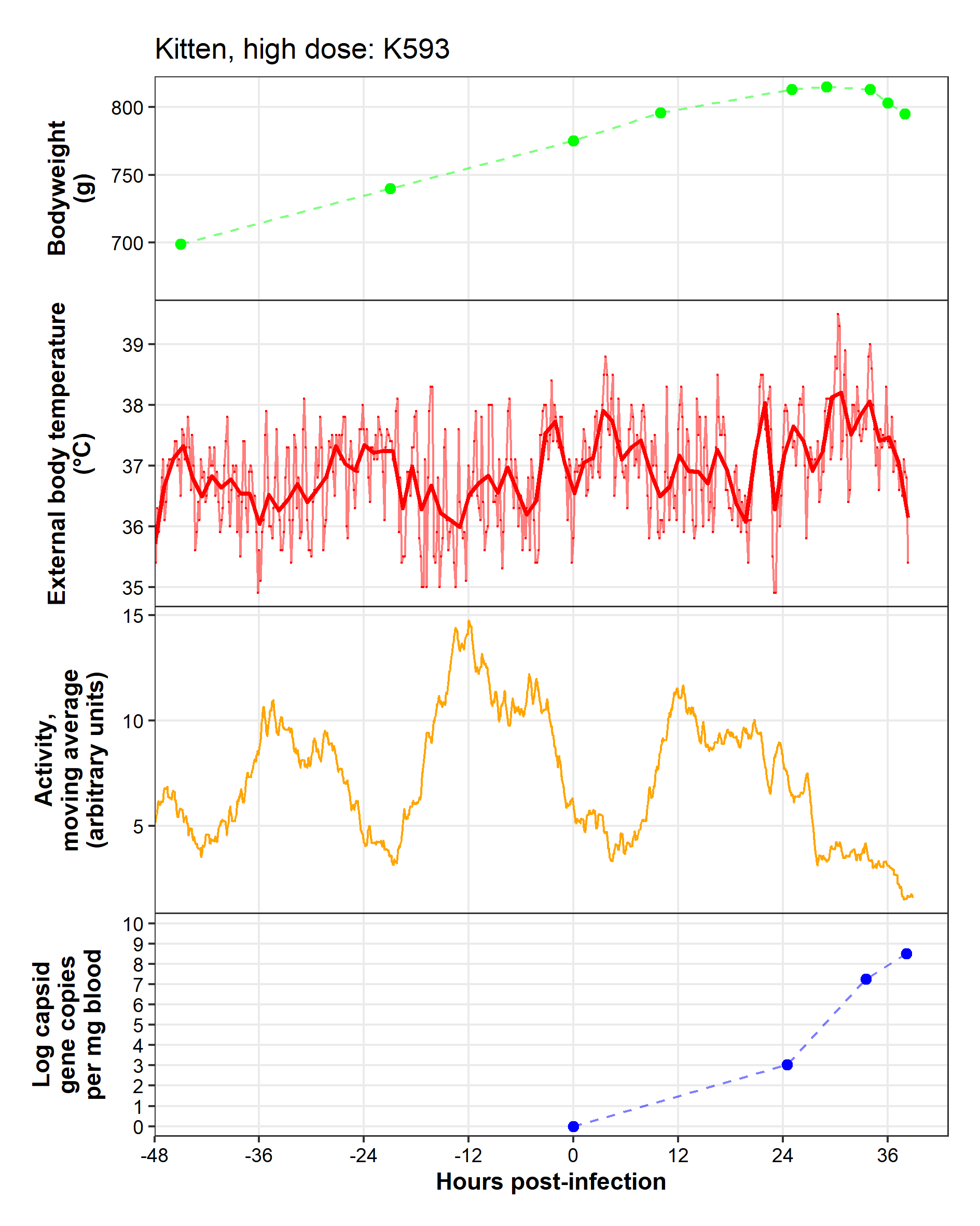

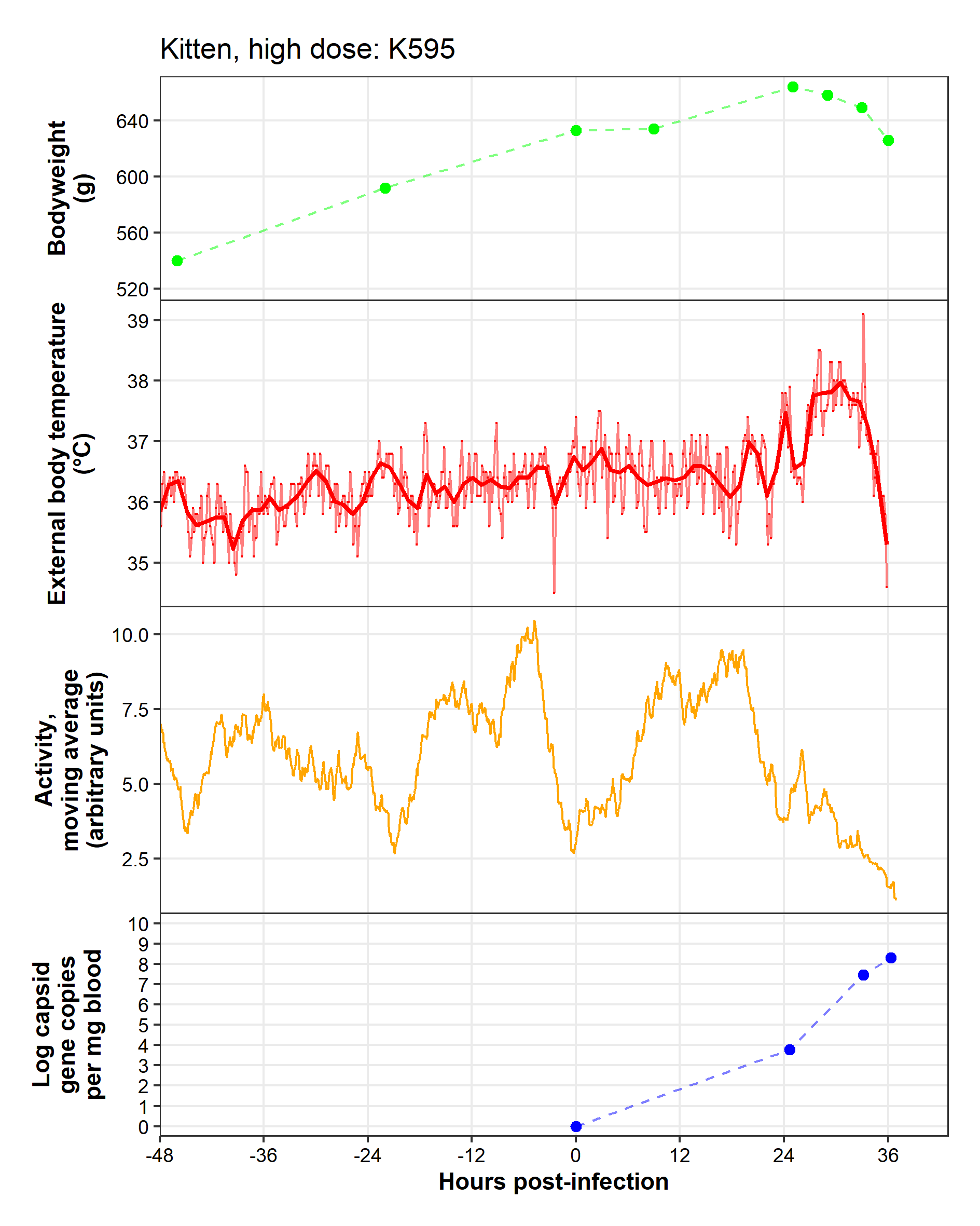

Kitten, low dose
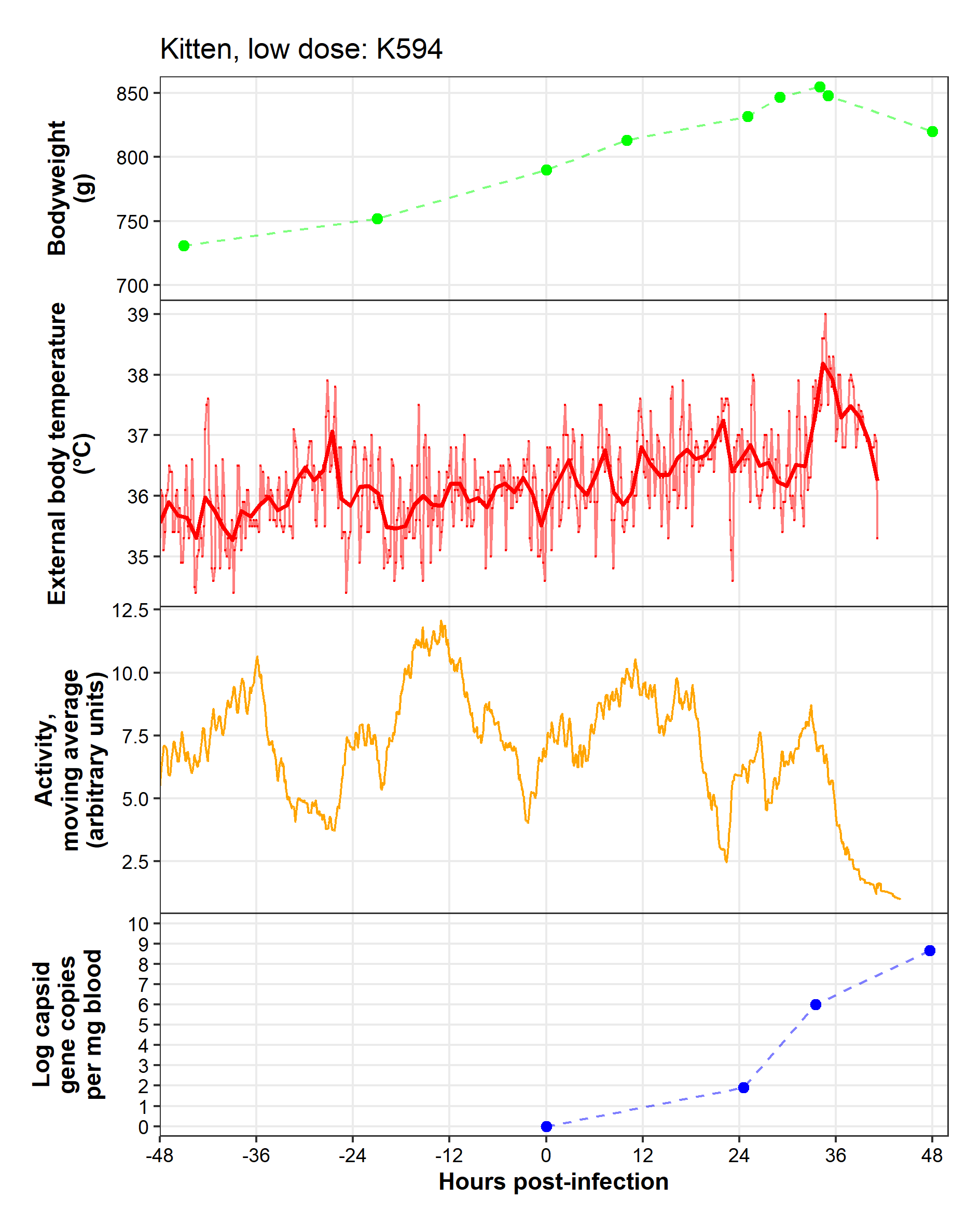

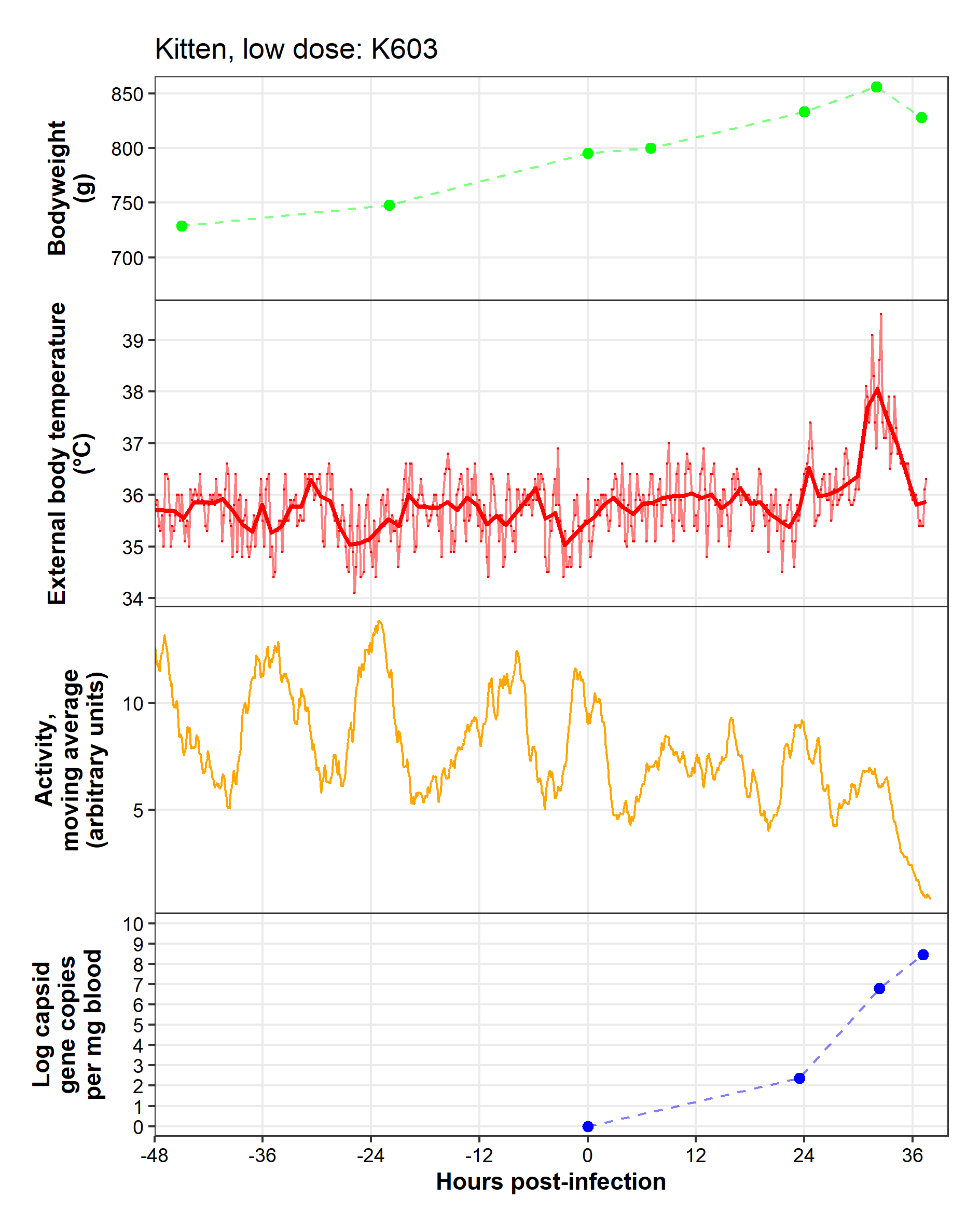

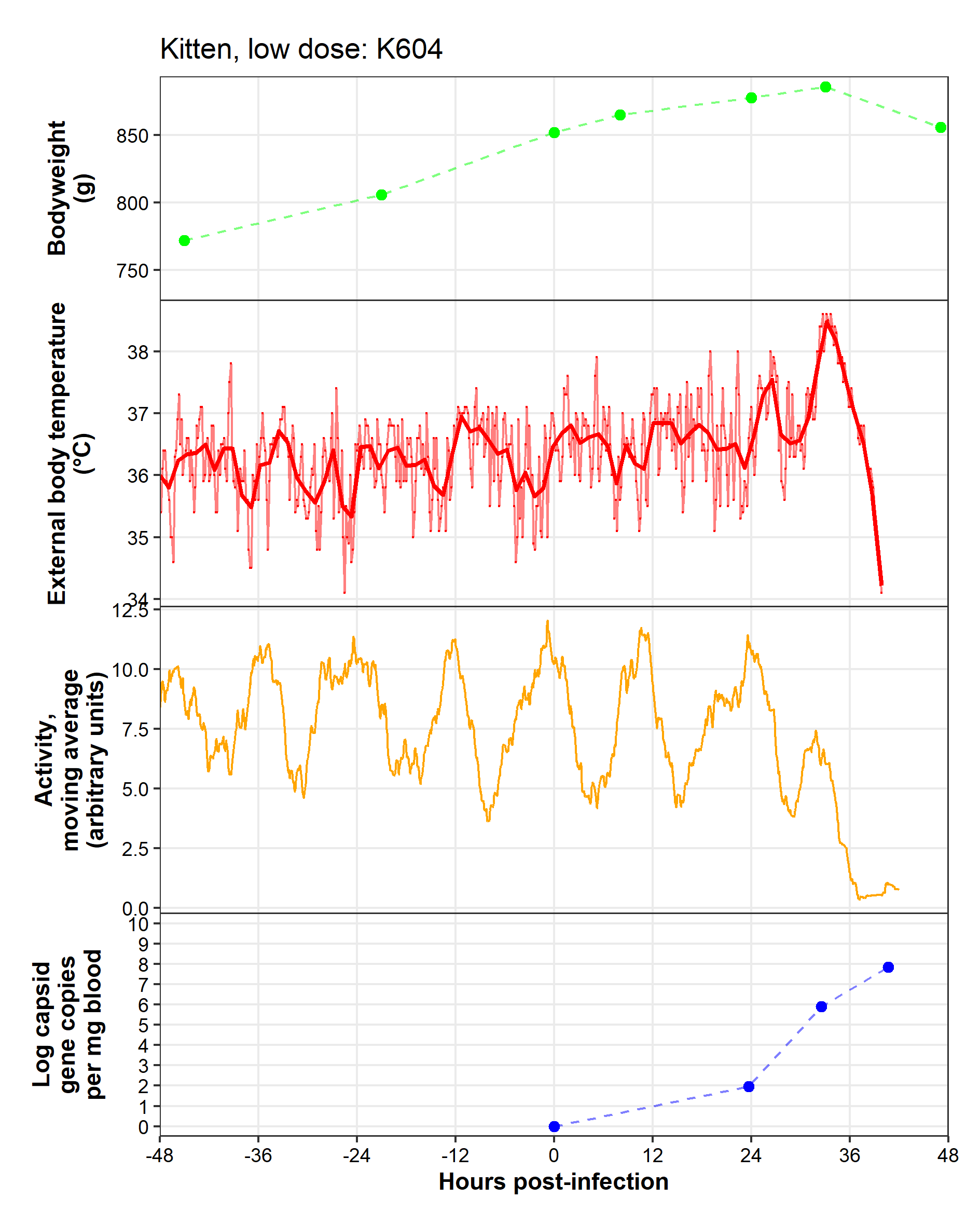

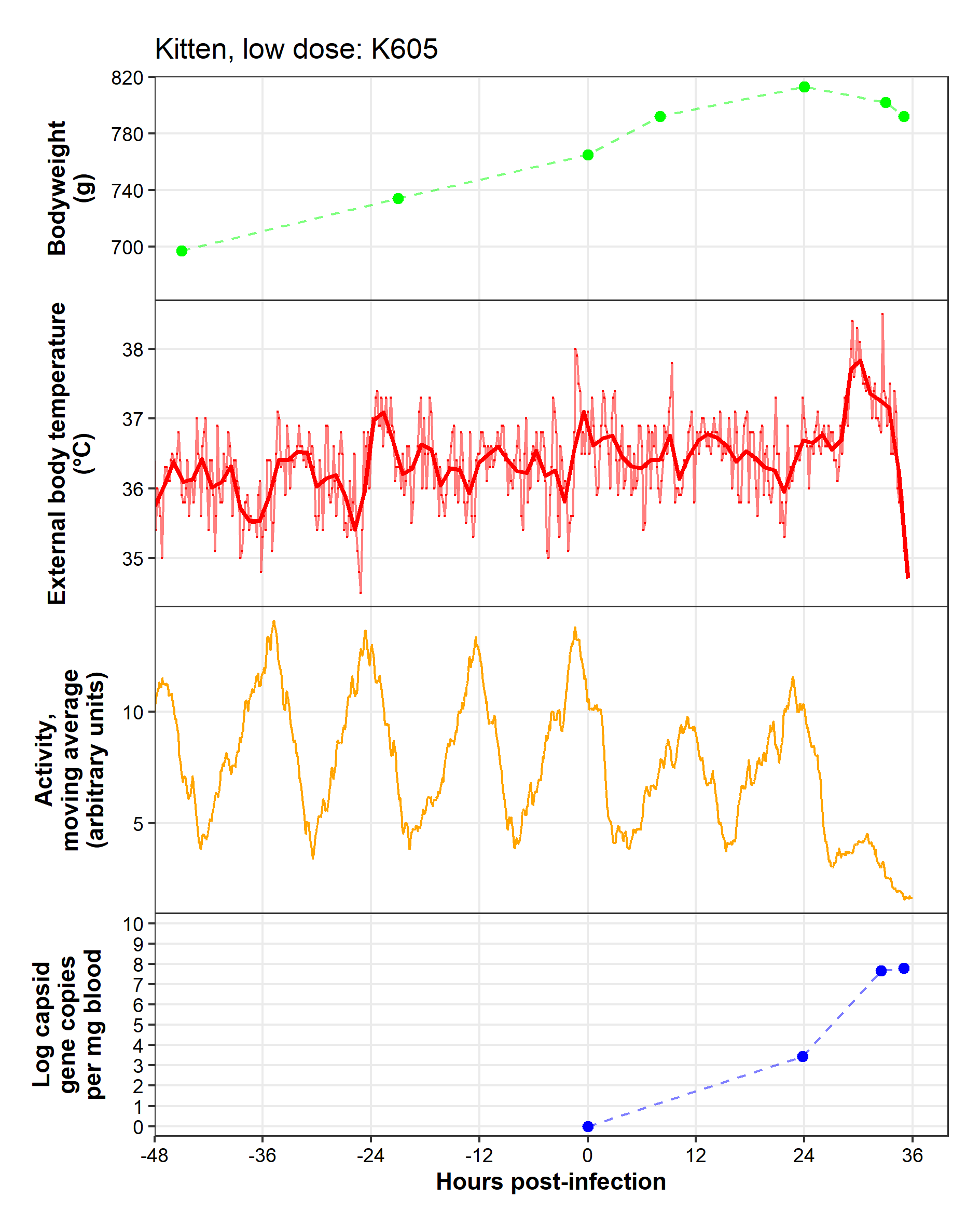

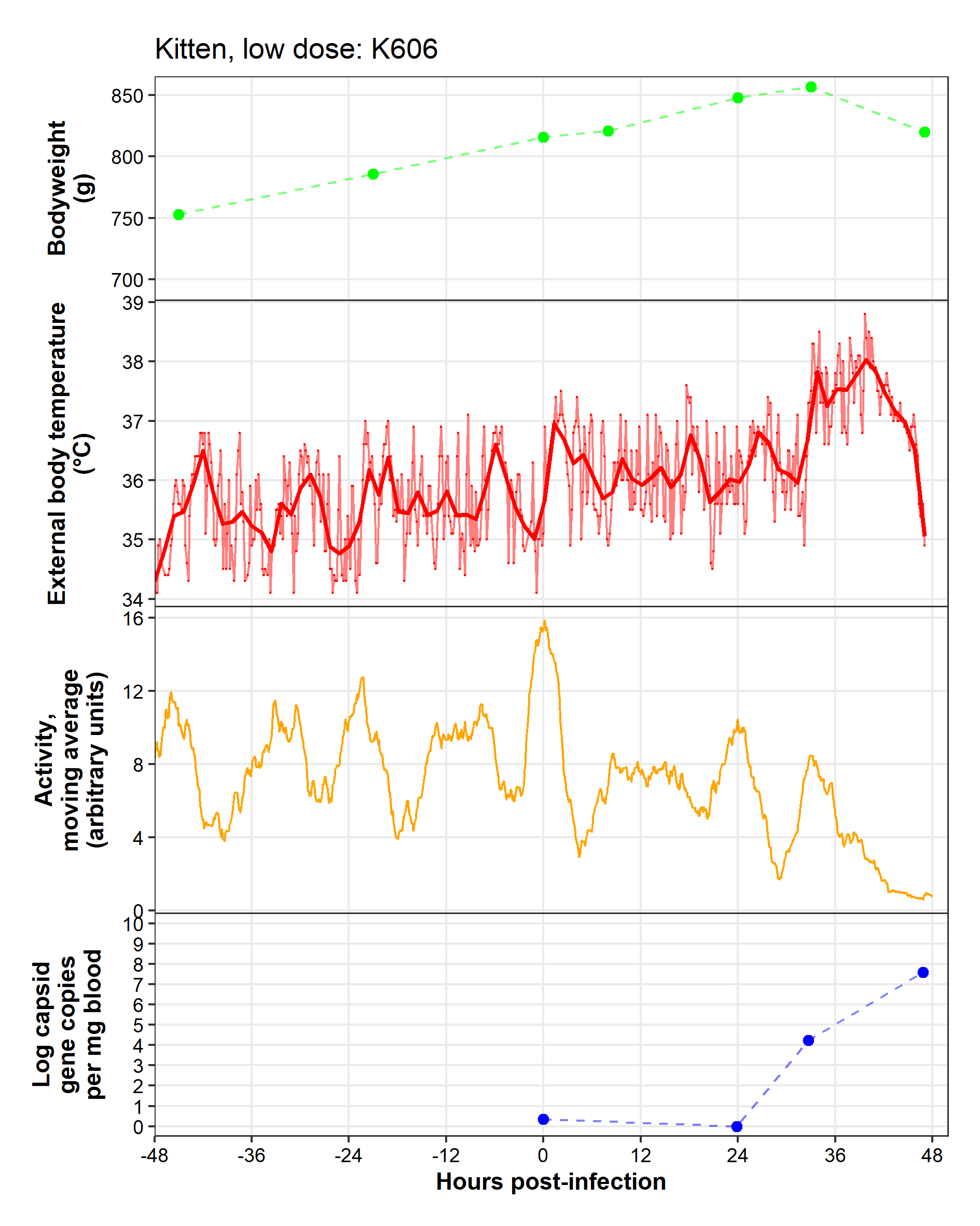

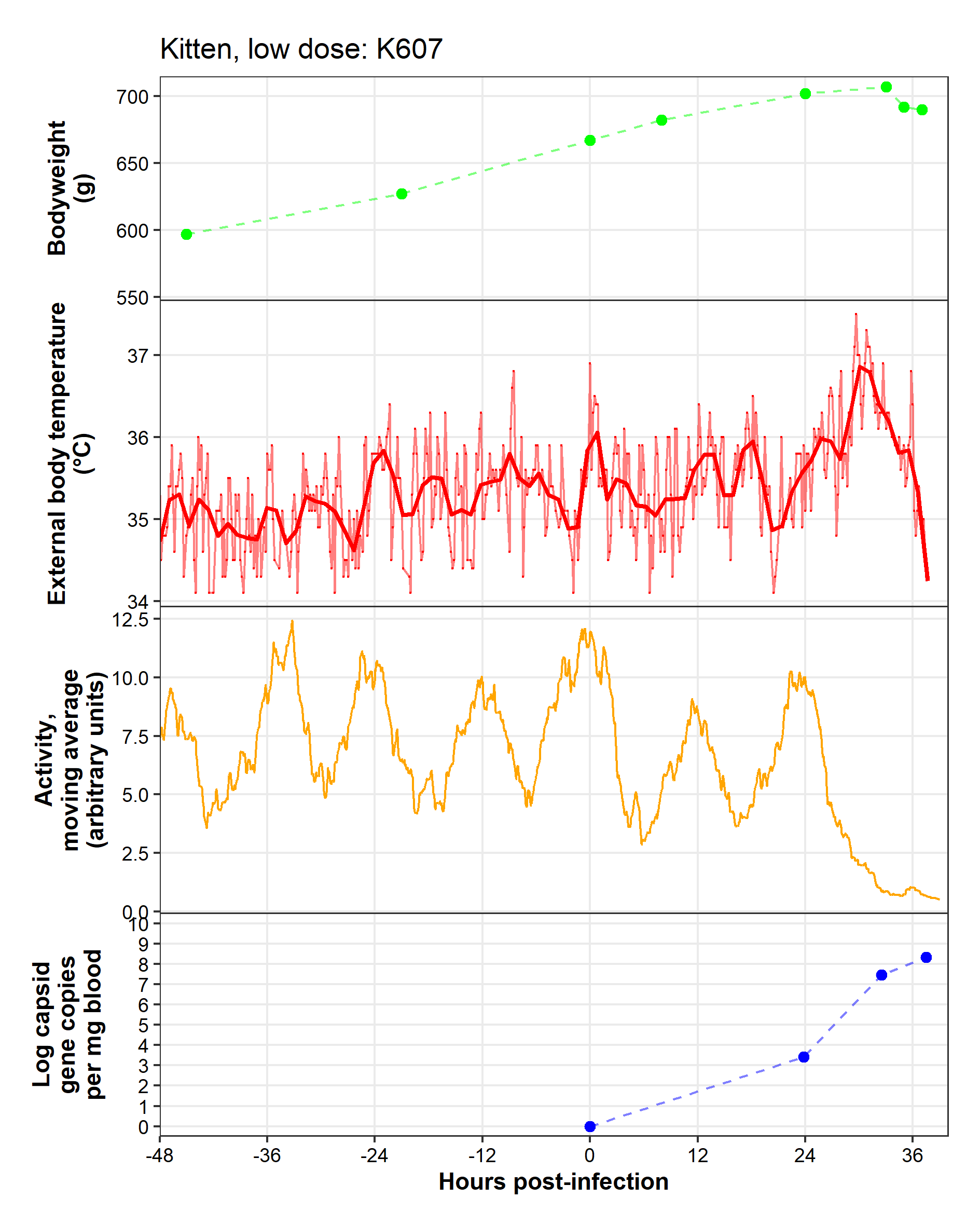

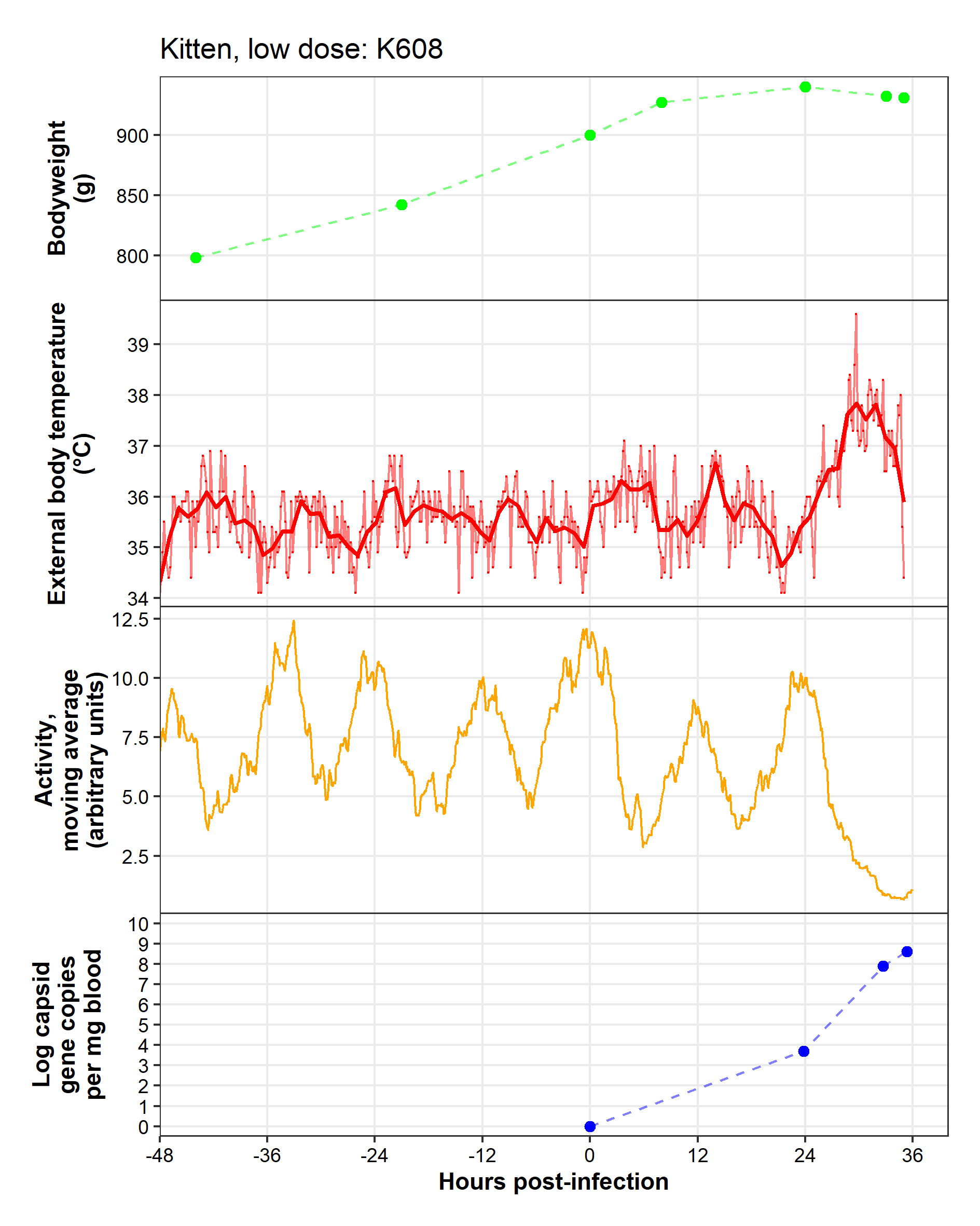

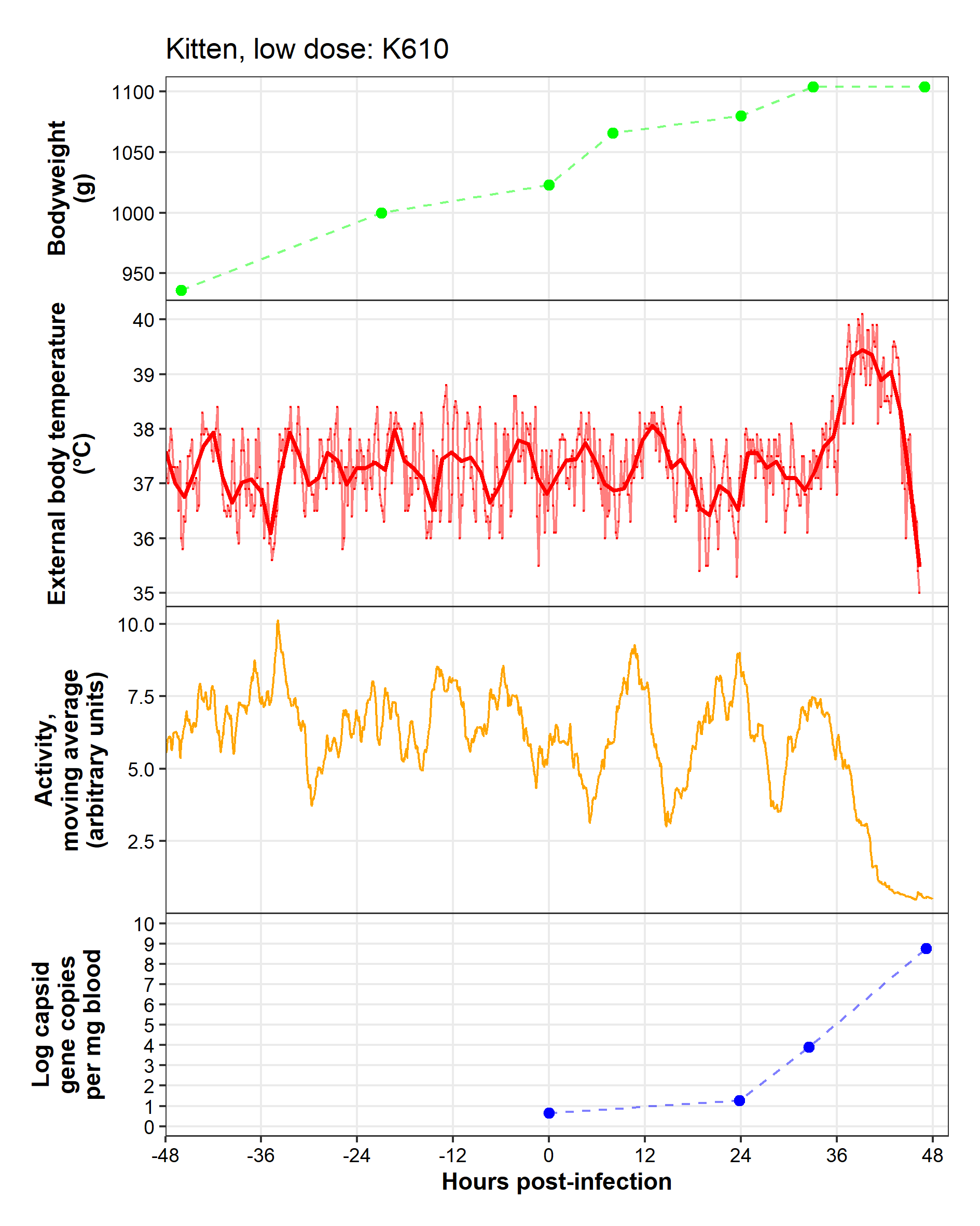

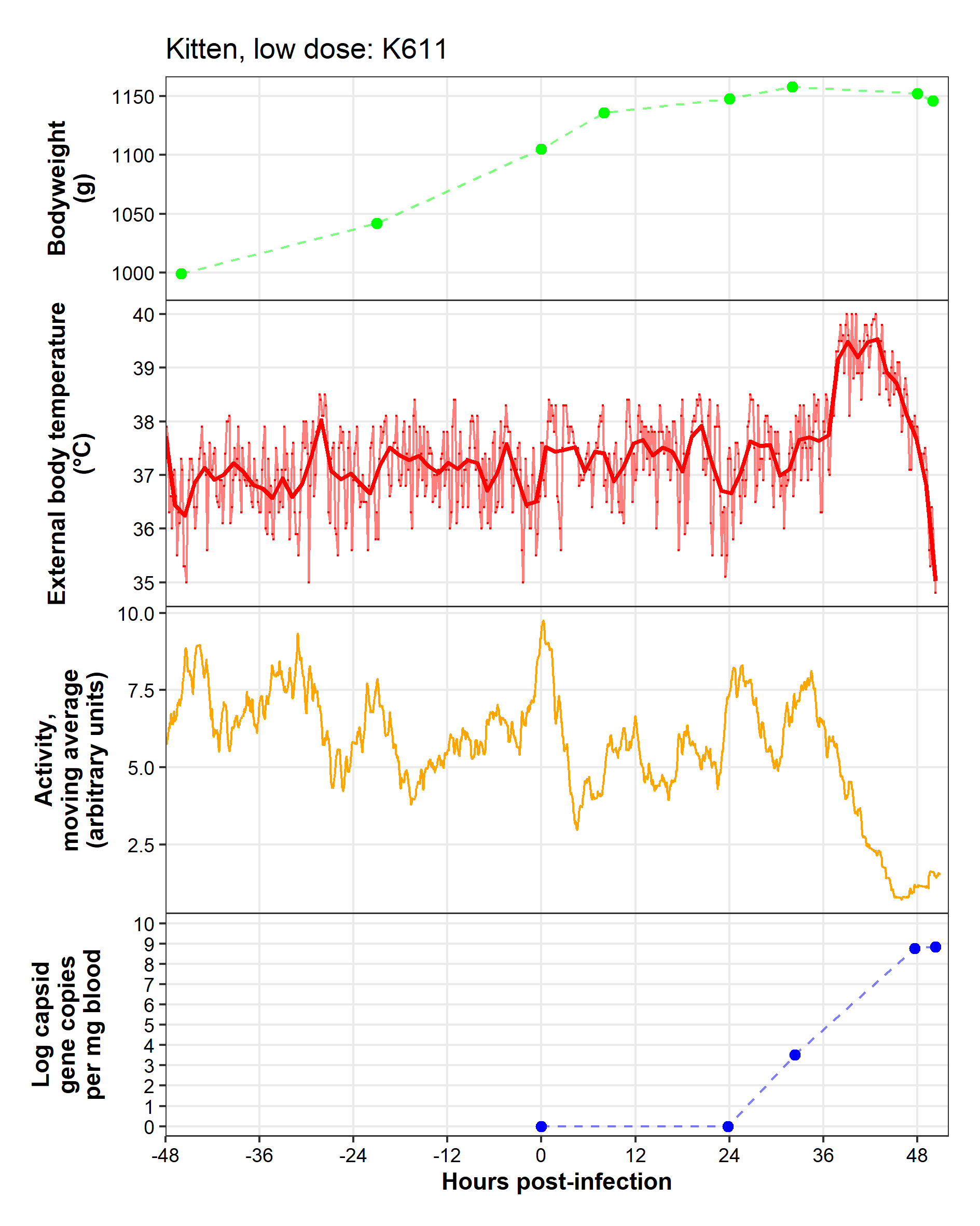

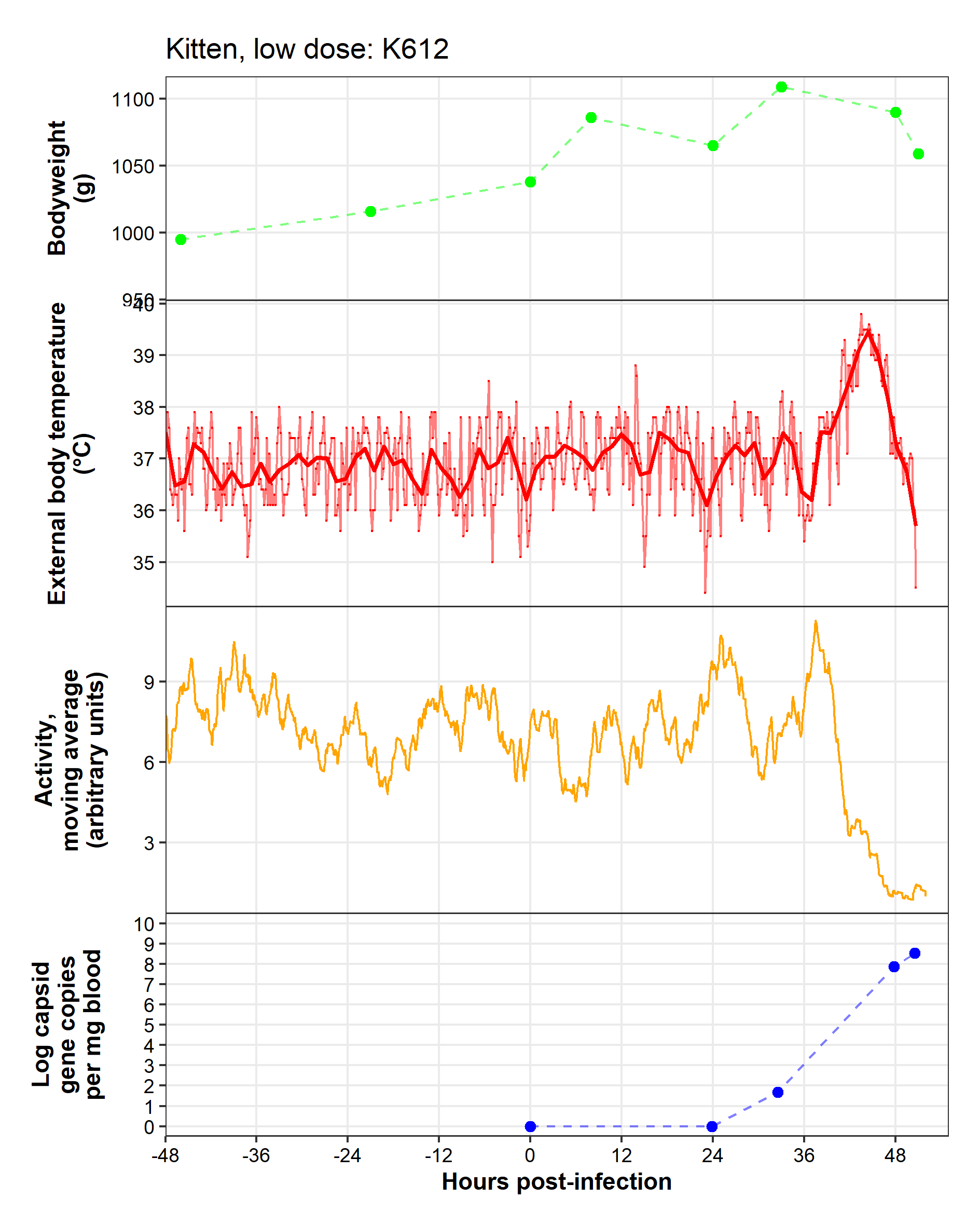

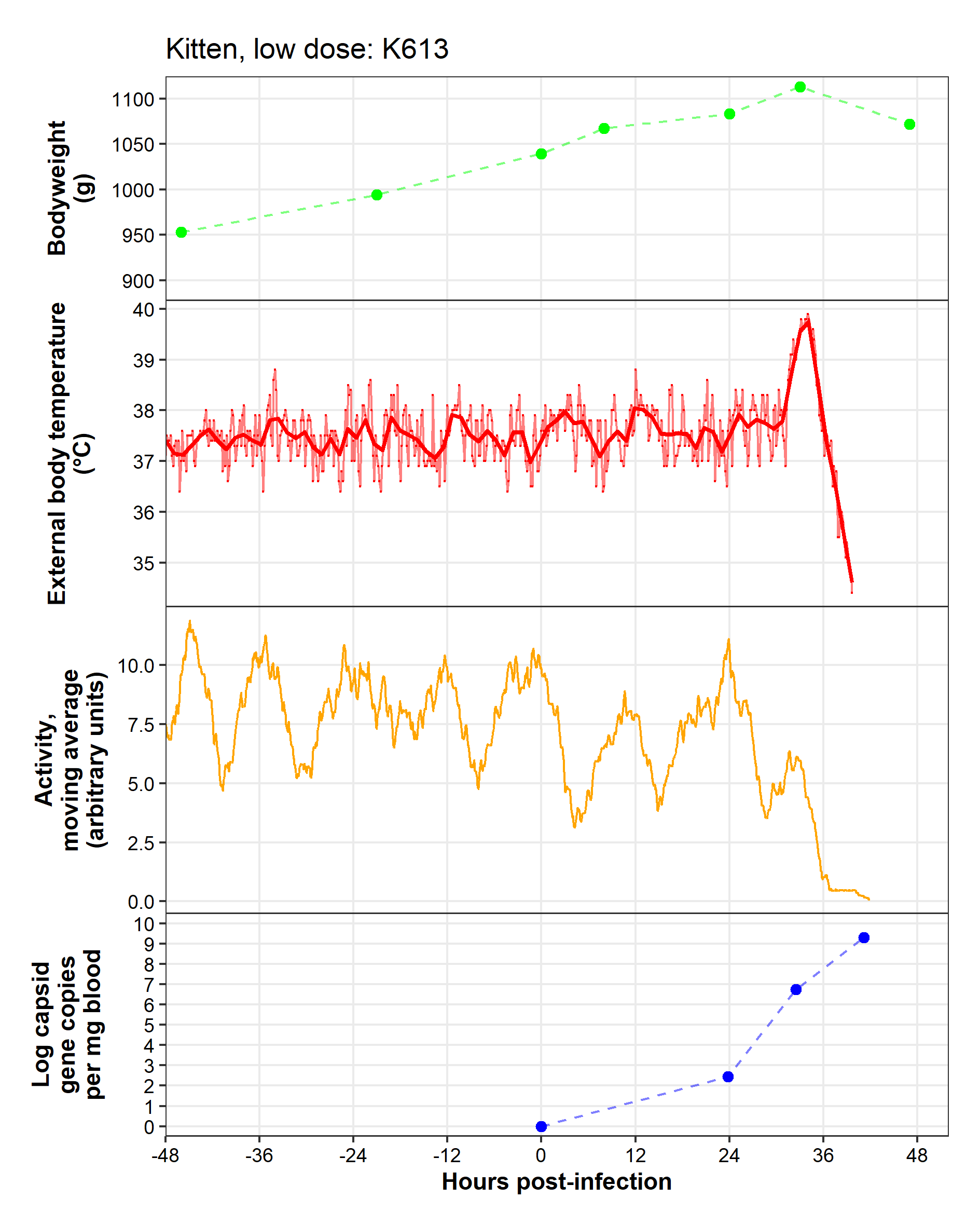

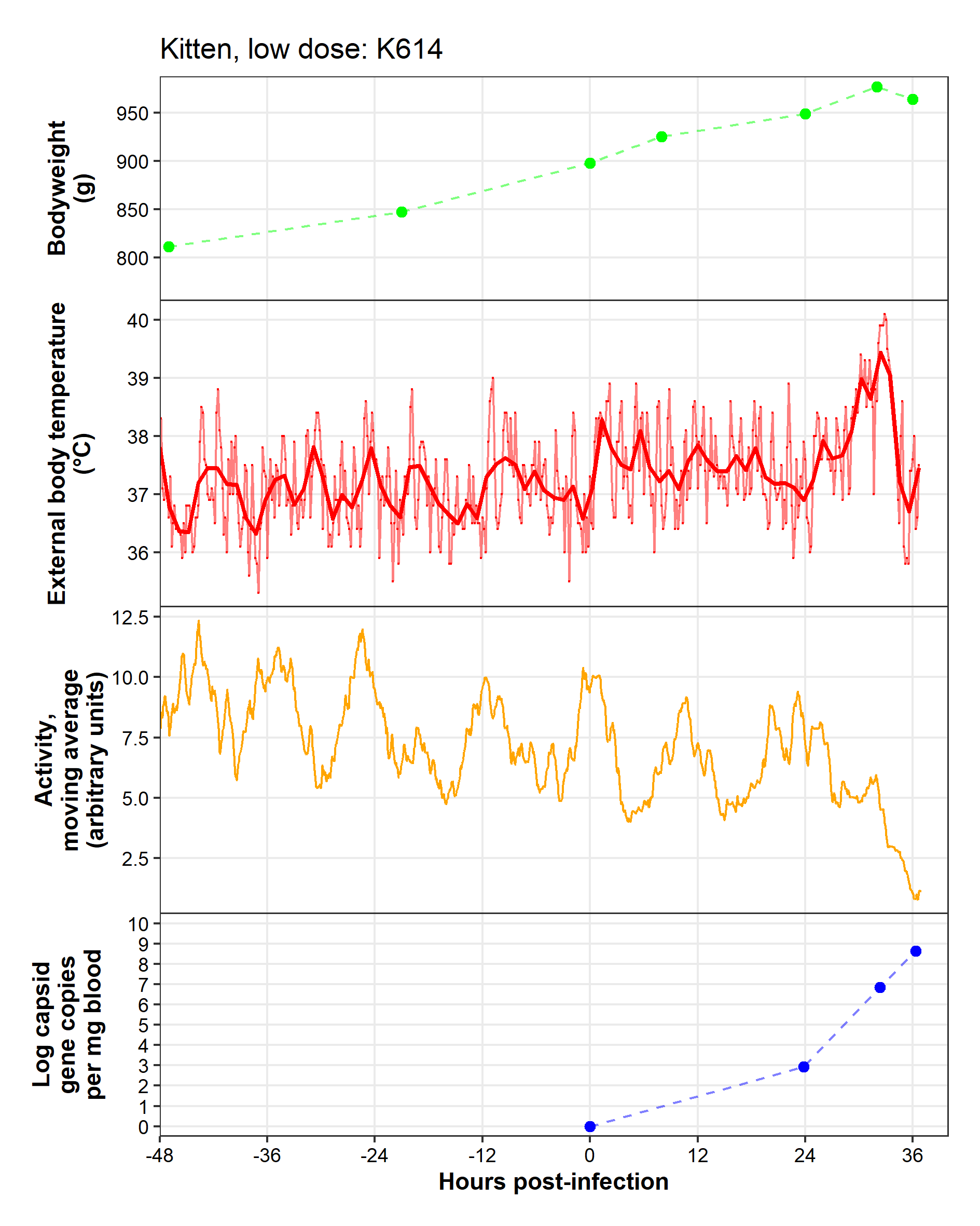
